# Side-Chain Dynamics of the α1B-Adrenergic Receptor determined by NMR via Methyl Relaxation

**DOI:** 10.1101/2023.05.09.539984

**Authors:** Christian Baumann, Wan-Chin Chiang, Renato Valsecchi, Simon Jurt, Mattia Deluigi, Matthias Schuster, Andreas Plückthun, Oliver Zerbe

**Author notes:** **Correspondence to Oliver Zerbe:**, Department of Chemistry, University of Zurich, 8057 Zurich, Winterthurerstrasse 190, Switzerland.

## Abstract

G protein-coupled receptors (GPCRs) are medically important membrane proteins that sample inactive, intermediate, and active conformational states characterized by relatively slow interconversions (∼μs– ms). On a faster timescale (∼ps–ns), the conformational landscape of GPCRs is governed by the rapid dynamics of amino acid side chains. Such dynamics are essential for protein functions such as ligand recognition and allostery. Unfortunately, technical challenges have almost entirely precluded the study of side-chain dynamics for GPCRs. Here, we investigate the rapid side-chain dynamics of a thermostabilized α_1B_-adrenergic receptor (α_1B_-AR) as probed by methyl relaxation. We determined order parameters for Ile, Leu, and Val methyl groups in the presence of inverse agonists that bind orthosterically (prazosin, tamsulosin) or allosterically (conopeptide ρ-TIA). Despite the differences in the ligands, the receptor’s overall side-chain dynamics are very similar, including those of the apo form. However, ρ-TIA increases the flexibility of Ile176^4x56^ and possibly of Ile214^5x49^, adjacent to Pro215^5x50^ of the highly conserved P^5x50^I^3x40^F^6x44^ motif crucial for receptor activation, suggesting differences in the mechanisms for orthosteric and allosteric receptor inactivation. Overall, increased Ile side-chain rigidity was found for residues closer to the center of the membrane bilayer, correlating with denser packing and lower protein surface exposure. In contrast to two microbial membrane proteins, in α_1B_-AR Leu exhibited higher flexibility than Ile side chains on average, correlating with the presence of Leu in less densely packed areas and with higher protein-surface exposure than Ile. Our findings demonstrate the feasibility of studying receptor-wide side-chain dynamics in GPCRs to gain functional insights.

## Introduction

G protein-coupled receptors (GPCRs) are medically important eukaryotic membrane proteins characterized by seven transmembrane (TM) α-helices. GPCRs detect extracellular stimuli, ranging from photons to small molecules and proteins, and transduce them into intracellular signals via conformational changes (Hilger et al. 2018; Kobilka 2013). To fulfill this function, GPCRs sample complex conformational landscapes. Therefore, a comprehensive picture of their conformational dynamics is vital to decipher the molecular mechanisms of receptor activation and inactivation. X-ray crystallography and single-particle cryo-electron microscopy (cryo-EM) are pivotal in providing GPCR structures in various activation states at high resolution (Danev et al. 2021; García-Nafría and Tate 2020; Grisshammer 2017). However, X-ray and cryo-EM structures are static snapshots, with only very few and limited exceptions (Gruhl et al. 2023; Matsumoto et al. 2021).

In contrast, nuclear magnetic resonance (NMR) can be used to study protein dynamics over a broad range of timescales, from the slow reorientations of domains (∼μs to s) to the fast motions of side chains (∼ps to ns). Dynamics in the range of ps to ns are interpreted using order parameters (S^2^_axis_). Order parameters range between zero (fully flexible) and one (completely rigid), reporting on how much bond vectors, such as the C-C bond vector of side-chain methyl groups, move on timescales faster than the overall tumbling of the protein. Based on their order parameters, side chains of increasing rigidity can be assigned to the motional classes J’, J, α, and ω (O’Brien et al. 2020; Sharp et al. 2014). The class J’ is only present in membrane proteins and contains side chains that are more dynamic than those found in soluble proteins (O’Brien et al. 2020). Side-chain dynamics are directly linked to the conformational entropy of a protein (Hoffmann et al. 2022), thereby influencing ligand binding, allosteric communication, and potentially providing a mechanism for allostery that does not require conformational changes (Cooper and Dryden 1984; Igumenova et al. 2006).

Methyl order parameters have been obtained for several soluble and, so far, three microbial membrane proteins, i.e., the β-barrel outer membrane protein W (OmpW), sensory rhodopsin II (pSRII), and bacteriorhodopsin (bR) (Kooijman et al. 2020b; O’Brien et al. 2020). Compared to soluble proteins, these three microbial membrane proteins showed increased side-chain dynamics and the absence of very rigid side chains. In the case of GPCRs, Clark et al. (Clark et al. 2017) determined relative side-chain dynamics of isoleucine residues in the agonist- and inverse agonist-bound adenosine A_2A_ receptor (A_2A_R), and found that the inverse agonist suppressed fast side-chain dynamics at the G protein-binding site. However, no methyl order parameters have yet been reported for GPCRs that allow comparisons with other proteins.

Unfortunately, GPCRs pose many technical challenges for NMR studies. The poor stability of most native GPCRs in membrane mimetics precludes extended measurements at the temperatures required to obtain NMR spectra of sufficient quality (typically, hours to weeks at 20–50°C). In addition, native GPCRs often have poor expression levels that complicate the isolation of sufficient amounts (up to several mg) of isotopically labeled receptors (Kim et al. 2009). In particular, isotopic labeling and especially the deuteration of GPCRs expressed in eukaryotic cells is difficult, which led the NMR community to resort to individual ^19^F reporter probes placed strategically within the receptors (Didenko et al. 2013; Liu et al. 2012; Picard and Prosser 2021). While stunning results have been obtained with this approach, the full potential of NMR is not yet exploited, as only individual probes have been used to examine receptor dynamics. In principle, NMR can provide receptor-wide dynamics by probing relaxation properties of many amide or methyl groups simultaneously, as is successfully done for soluble proteins (Palmer 2004).

To provide a more comprehensive view of GPCR side-chain dynamics, we report here on the dynamics of methyl groups, including their order parameters, of a thermostabilized human α_1B_-adrenergic receptor (or α_1B_ -adrenoceptor, α_1B_-AR) bound to three inverse agonists as well as in the apo form. The thermostabilized α_1B_-AR construct has been optimized for expression in the inner membrane of *E. coli* (Schuster et al. 2020), allowing us to apply established protocols for ^13^C,^1^H labeling of δ-methyl groups in Ile and Leu side chains, and γ-methyl groups in Val side chains (Figure 1) (Kerfah et al. 2015; Tugarinov et al. 2006; Tugarinov et al. 2007). The crystal structure of a related α_1B_-AR construct has been determined bound to the inverse agonist (+)-cyclazosin, which is a close analog of prazosin, which was used in this study (Deluigi et al. 2022). Since the mutations harbored by the thermostabilized α_1B_ - AR construct used for NMR and crystallography lock the receptor in a signaling-inactive state (Deluigi et al. 2022), we focused our NMR study on ligands that stabilize the inactive state(s) of α_1B_-AR, i.e., inverse agonists (Berg and Clarke 2018).

**Figure 1.**
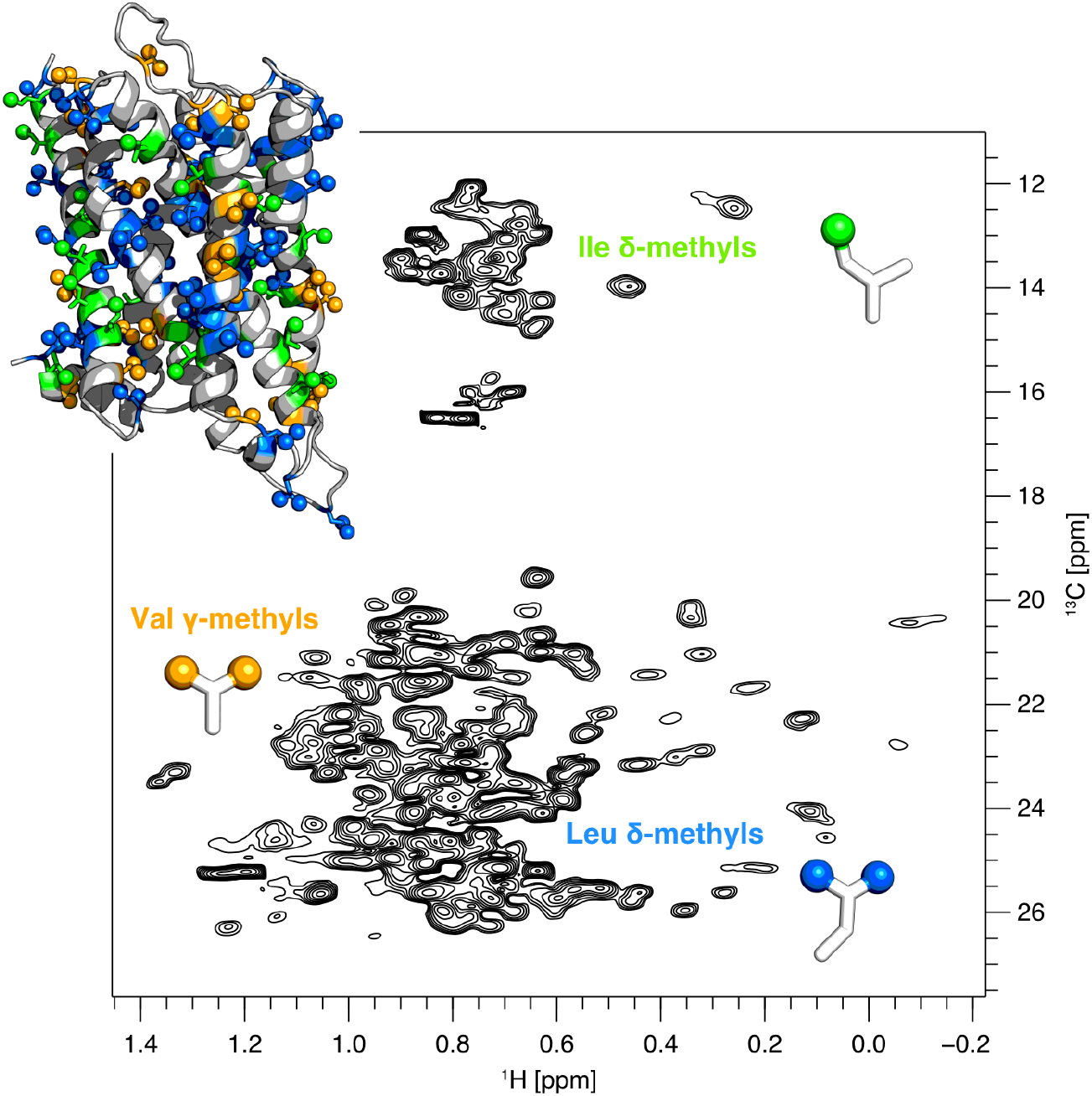
Structural model and [^13^C,^1^H]-HSQC spectrum of the thermostabilized α_1B_-AR denoted α_1B_-AR-B1D1 bound to prazosin. The receptor was expressed in *E. coli*, enabling ^13^C and ^1^H labeling of δ-methyl groups in Ile (green) and Leu residues (blue), and γ-methyl groups in Val residues (orange). These methyl groups are depicted as spheres on a homology model of α_1B_-AR-B1D1 and on the side chains within the spectrum. The receptor is probed globally using these three amino acids, constituting roughly a third of the entire amino acid content (24 Ile, 47 Leu, and 28 Val in 303 residues).

The inverse agonists prazosin and tamsulosin (Figure 2a) are clinically prescribed small molecules that bind to the orthosteric ligand-binding site of α_1B_-AR (Rossier et al. 1999), i.e., the site located within the transmembrane bundle targeted by the endogenous agonists adrenaline and noradrenaline, as well as to adjacent regions (Deluigi et al. 2022). In contrast, the inverse agonist ρ-TIA, a toxin produced by the cone snail *Conus tulipa* to hunt fish, is a 19-amino acid peptide that binds to a distinct, allosteric site primarily located within the extracellular surface of α_1B_-AR (Figure 2a) (Chen et al. 2004; Ragnarsson et al. 2013; Sharpe et al. 2001; Sharpe et al. 2003). Besides their different chemical structures and either orthosteric or allosteric binding modes, the three ligands also display different selectivities for adrenergic receptor subtypes (Chen et al. 2004; Michel et al. 2020; Proudman et al. 2020).

**Figure 2.**
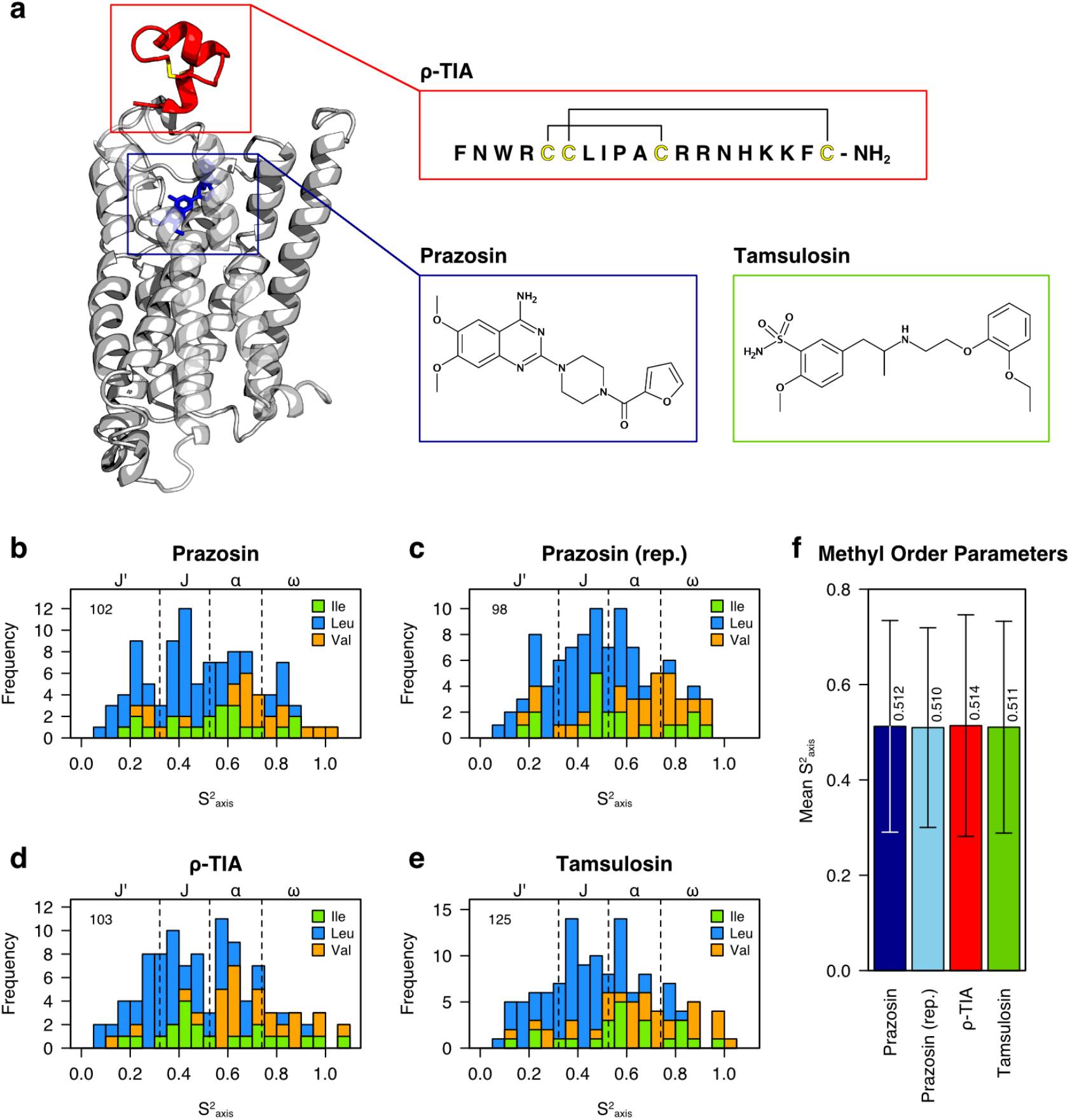
Structural model of α_1B_-AR-B1D1 depicting the binding sites of the three investigated inverse agonists and the obtained side-chain dynamics. **(a)** Prazosin and tamsulosin are small-molecule inverse agonists that bind within the transmembrane helical bundle and fill the orthosteric binding pocket, while ρ-TIA is a conopeptide that binds allosterically at the extracellular receptor surface. The binding of ρ-TIA to the structural model of α_1B_-AR-B1D1 was based on Ragnarsson et al. (Ragnarsson et al. 2013) **(b–e)** Histograms of the methyl order parameters determined for α_1B_-AR-B1D1 bound to the different inverse agonists. Bars are colored according to the amino acid: Ile in green, Leu in blue, and Val in orange. Dashed lines at S^2^_axis_ of 0.32, 0.53, and 0.74 depict borders between motional classes (J’, J, α, ω) as determined using k-means clustering on all data sets combined. The values for α_1B_-AR-B1D1 bound to prazosin are shown for two independently recorded NMR experiments using the same sample in (b) and (c). The total number of obtained methyl order parameters from each experiment is indicated in the top left corner. A more detailed analysis comparing populations within the motional classes did not reveal significant differences (SD1 Section 4.2). **(f)** Average methyl order parameter in the presence of the indicated ligand. Error bars represent standard deviations.

We thus investigated whether the differences between the ligands are reflected in changes in the receptor side-chain dynamics based on methyl order parameters. In addition, we measured the relative side-chain dynamics of the apo and prazosin-bound α_1B_-AR. Further, we assigned most Ile methyl groups by mutagenesis, allowing us to map side-chain dynamics onto the α_1B_-AR crystal structure and compare them with structural properties, such as packing density and surface exposure of the side chains. The remaining methyl resonances were assigned to the amino acid type (Leu or Val), which enabled us to compare the dynamics between different side chains.

## Results

### Receptor construct and NMR experiments

We used the stable α_1B_-AR construct denoted α_1B_-AR-B1D1 (Schuster et al. 2020). Compared to the native human α_1B_-AR, α_1B_-AR-B1D1 harbors 13 amino acid mutations, the deletion of residues Gly240– Phe284 in the third intracellular loop (ICL3), as well as N- and C-terminal truncations at residues Ser35 and Gly369, respectively (see Supplementary Data 1 (SD1) Section 1.1). We determined side-chain dynamics for Ile, Leu, and Val residues by measuring proton triple-quantum coherence relaxation rates on methyl groups, following the methodology developed by Sun, Kay, and Tugarinov (Sun et al. 2011). The δ-methyl of Ile, one of the two δ-methyls of Leu, and one of the two γ-methyls of Val side chains were ^13^C, ^1^H labeled, whereas the rest of the side chain contained ^12^C and ^2^H nuclei. One of the two methyl groups in Leu and Val was exclusively but non-stereospecifically labeled. In addition, the receptor was uniformly ^2^H,^15^N labeled. We obtained methyl order parameters for α_1B_-AR-B1D1 bound to the inverse agonists prazosin, tamsulosin, and ρ-TIA in detergent micelles at 320 K. We have previously shown that another α_1B_-AR construct, which differs from α_1B_-AR-B1D1 by having the full-length ICL3, retains high-affinity binding to prazosin (K_D_ ≈ 0.8 nM) (Schuster et al. 2020). The binding of tamsulosin and ρ-TIA by α_1B_-AR-B1D1 is evident from the changes in the [^13^C,^1^H]-HSQC spectra when the different ligands were added (Figure S2.1 and S2.2). Side-chain dynamics for apo α_1B_-AR-B1D1 were studied at 298 K due to the insufficient stability of the apo receptor at 320 K.

### Influence of ligands on side-chain dynamics

The obtained methyl order parameters for α_1B_-AR-B1D1 bound to the three different inverse agonists cover almost the entire range of possible values (Figure 2b–e). Interestingly and in contrast to α_1B_-AR-B1D1, the other membrane proteins investigated so far had no side chains belonging to the most rigid motional class ω (Kooijman et al. 2020b; O’Brien et al. 2020). The average of the methyl order parameters for α_1B_-AR-B1D1 in micelles was 0.51 (Figure 2f), which is larger than the values reported for bacteriorhodopsin (bR) in nanodiscs (Kooijman et al. 2020b) (mean S^2^_axis_ = 0.41) and for sensory rhodopsin II (pSRII) (O’Brien et al. 2020) in micelles (mean S^2^_axis_ = 0.37) or bicelles (mean S^2^_axis_ = 0.45). This indicates that α_1B_-AR-B1D1 adopts a more rigid structure (on a ps to ns timescale) than the two microbial rhodopsins. At present, it is unclear whether the increased rigidity in α_1B_-AR-B1D1 is a native property of inactive α_1B_-AR, is due to the stabilizing mutations, or to stabilizing effects of the detergent lauryl maltose neopentyl glycol (LMNG) (Lee et al. 2020), or a combination of these factors. In addition, the absence of native trimer contacts in bR and missing contacts between pSRII and its transducer protein may increase the overall flexibility of these two proteins compared to their natural environment.

The methyl order parameters for α_1B_-AR-B1D1 were distributed similarly in the three different inverse agonist complexes (Figure 2b–e). To test whether the three ligands caused receptor-wide changes in side-chain dynamics, the percentage of order parameters within each motional class was compared between the ligands. Only minor fluctuations were present in the percentages per class between the ligands, comparable to the fluctuations between duplicate measurements of α_1B_-AR-B1D1 bound to prazosin (Figure S4.2.1). Thus, no major receptor-wide differences in side-chain dynamics were observed for the investigated ligand complexes of α_1B_-AR-B1D1. However, localized or compensatory (Wankowicz et al. 2022) changes in dynamics might escape detection when comparing the overall distributions of order parameters between ligands. Such changes can, however, be detected by comparing dynamics at the residue level (see below). The relative side-chain dynamics data of apo α_1B_ - AR-B1D1 display decreased values compared to the prazosin-bound receptor, suggesting increased side-chain dynamics (Figure S4.3.1). Unfortunately, due to the inability to determine the overall correlation time at 298 K, it is more likely that the observed differences are due to different rotational correlation times rather than changes in side-chain dynamics (SD1 Section 4.3).

### Influence of ligands on Ile side-chain dynamics

While no differences in global side-chain dynamics were apparent between different liganded forms of α_1B_-AR-B1D1, local changes in dynamics may be resolved by comparing the order parameters of assigned residues (Figure 3a, S4.4.1 and S4.4.2). To this end, we assigned the majority of Ile residues using point mutations to either Leu or Val. Ile residues are located throughout the receptor (Figure 1), and their S^2^_axis_ values span all motional classes (Figure 3a), thereby providing convenient probes. The most rigid Ile side chains belong to I56^1x43^ and I60^1x47^ in TM1 (superscripts denote GPCRdb numbering (Isberg et al. 2015)). The most dynamic Ile side chains are those of I42^1x29^ and I178^4x58^ at the extracellular ends of TM1 and TM4, respectively (Figure 3b). The methyl group of I178^4x58^ gives rise to two peaks in the presence of prazosin and tamsulosin (Figure 3c), indicating either two distinct conformational states of I178^4x58^ or a nearby conformational change with an interconversion rate slower than 50 s^-1^. Methyl order parameters derived from both I178^4x58^ signals indicate that side-chain dynamics are identical in these two conformational states (Figure S4.4.3).

**Figure 3.**
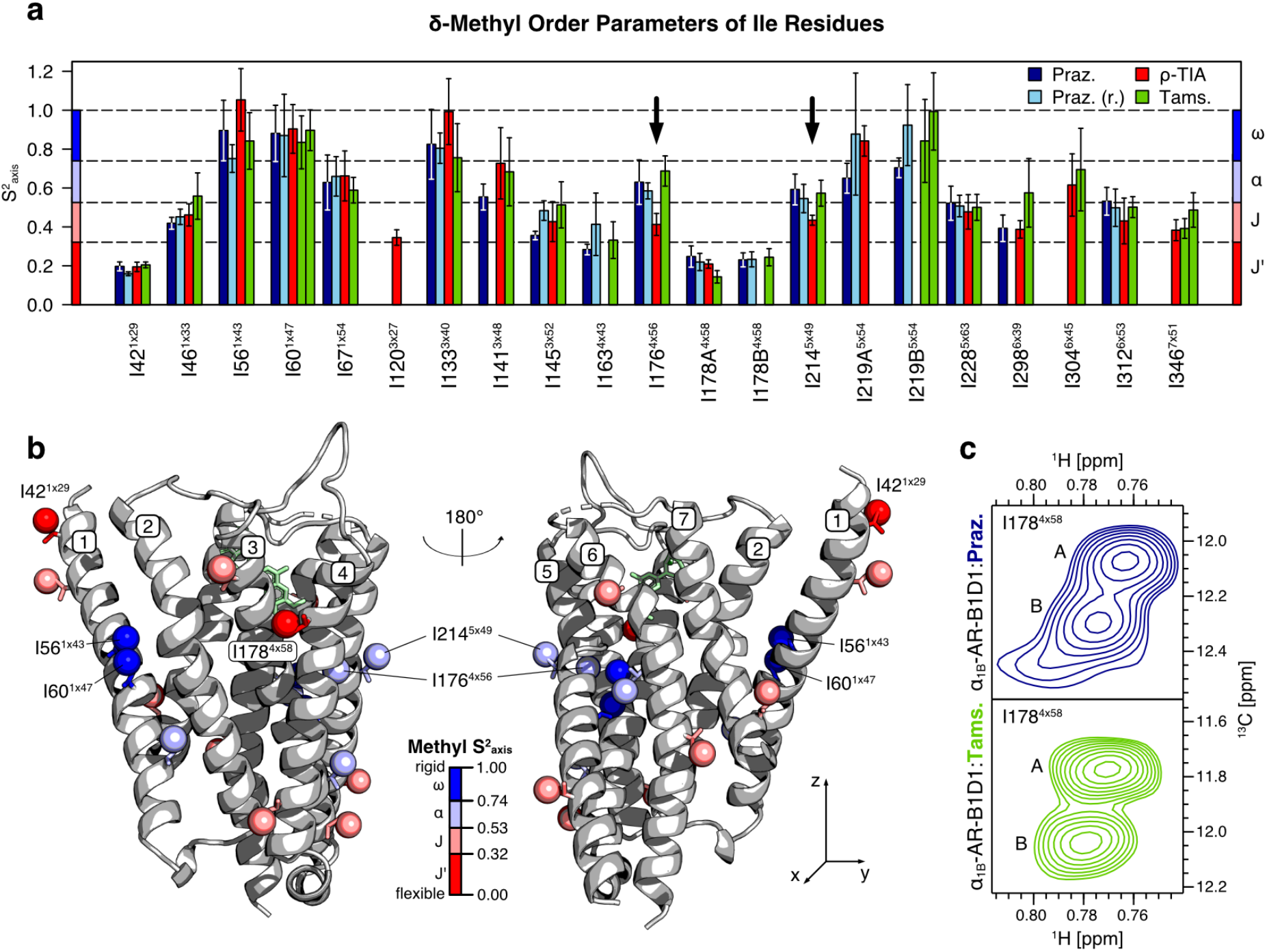
δ-methyl order parameters of assigned Ile residues. **(a)** Bar plots of Ile δ-methyl order parameters as measured with each indicated ligand. Arrows highlight I176^4x56^ and I214^5x49^ S^2^_axis_ values, which changed notably with ρ-TIA compared to prazosin and tamsulosin. Error bars indicate the S^2^_axis_ standard errors from the fit (for 95 % CI see Figure S4.4.1 and S4.4.2). Residues that lead to multiple signals in [^13^C,^1^H]-HSQC spectra for more than one ligand are plotted separately and labeled alphabetically (Figure S2.4). Tamsulosin led to multiple signals for several residues, whose S^2^_axis_ values are shown next to one another in the bar plot (I60^1x47^, I219B^5x54^, I346^7x51^). Missing bars indicate that no reliable values could be obtained. **(b)** Crystal structure of inverse agonist-bound α_1B_-AR (Protein Data Bank (PDB) entry 7B6W) (Deluigi et al. 2022) with Ile δ-methyl groups shown as spheres. Methyl groups are colored according to the motional class of the mean S^2^_axis_ across all ligand-bound samples. The locations of the two most flexible (I42^1x29^ and I178^4x58^), the two most rigid Ile side chains (I56^1x43^ and I60^1x47^) as well as the two residues with the most notable ligand-induced changes in dynamics (I176^4x56^ and I214^5x49^) are indicated on the structure. The side chains of both I176^4x56^ and I214^5x49^ show enhanced motions in the presence of ρ-TIA compared to prazosin and tamsulosin. **(c)** [^13^C,^1^H]-HSQC peaks assigned to the δ-methyl group I178^4x58^ in the prazosin- and tamsulosin-bound receptor.

The two most notable ligand-induced changes in side-chain dynamics were observed for I176^4x56^ and I214^5x49^ (Figure 3a and S4.4.2). Both residues are located on the same “face” of α_1B_-AR, and their methyl carbons are separated by 8.3 Å (Figure 3b). The side chains of both I176^4x56^ and I214^5x49^ have smaller order parameters (i.e., they are more flexible) when the receptor is bound to ρ-TIA compared to prazosin and tamsulosin. The increase in dynamics corresponds to a transition from the α to the J class for both side chains, hinting at a substantial increase in motional freedom. These differences in S^2^_axis_ values between ligands, however, are similar to some of those between duplicate prazosin measurements (see e.g. I145^3x52^) and hence must be viewed with caution. Nonetheless, since two probes in relative proximity (I176^4x56^ and I214^5x49^) show substantially increased dynamics in the presence of ρ-TIA, it is less likely that this is due to measurement errors or uncertainties. Among other potential (albeit not significant) differences between the ligand-bound forms, ρ-TIA seems to rigidify I133^3x40^ compared to prazosin and tamsulosin. I133^3x40^ is interesting as it belongs to the P^5x50^I^3x40^F^6x44^ motif, a highly conserved amino acid triad involved in GPCR activation, and it is part of the transmission switch motif formed by I^3x40^, L^5x51^, F^6x44^, and W^6x48^ (Zhou et al. 2019).

### Ile δ-methyl order parameters and overall structural properties

When Ile δ-methyl order parameters are mapped on the α_1B_-AR crystal structure bound to the inverse agonist (+)-cyclazosin (PDB entry 7B6W), it appears that the extent of motion follows the protein z-axis bidirectionally, perpendicular to the membrane plane (Figure 3b): The most rigid side chains are mainly located close to the receptor center along the z-axis (where the center of the phospholipid bilayer would be), whereas more flexible side chains are primarily located toward the intra- and extracellular ends of the receptor. This could be expected for a receptor that transmits a signal from its extracellular to its intracellular side and thus requires a well-organized central segment to relay the conformational changes between these two faces of the receptor.

To investigate how the receptor structure influences side-chain dynamics, we determined the correlations between the Ile δ-methyl S^2^_axis_ values and the structural properties of those δ-methyl groups in the α_1B_-AR crystal structure (PDB entry 7B6W). The correlations to the following three structural properties were analyzed: (1) the position of the δ-methyl groups on the protein z-axis perpendicular to the membrane plane, (2) the atom density (denoted as “packing” herein), quantified by counting atoms within 5 Å of the δ-methyl group, and (3) the side-chain exposure on the protein surface, quantified by the solvent accessible surface area (Figure 4). These three structural properties are not mutually independent but describe overlapping properties of the structure, i.e., stronger packing is found toward the center of the transmembrane region, whereas side chains on the protein surface are less densely packed. The Spearman’s rank correlation coefficient (ρ) was used to determine the extent of the correlations (ρ = 1 for perfectly positive, ρ = −1 for perfectly negative, and ρ = 0 for no correlation).

**Figure 4.**
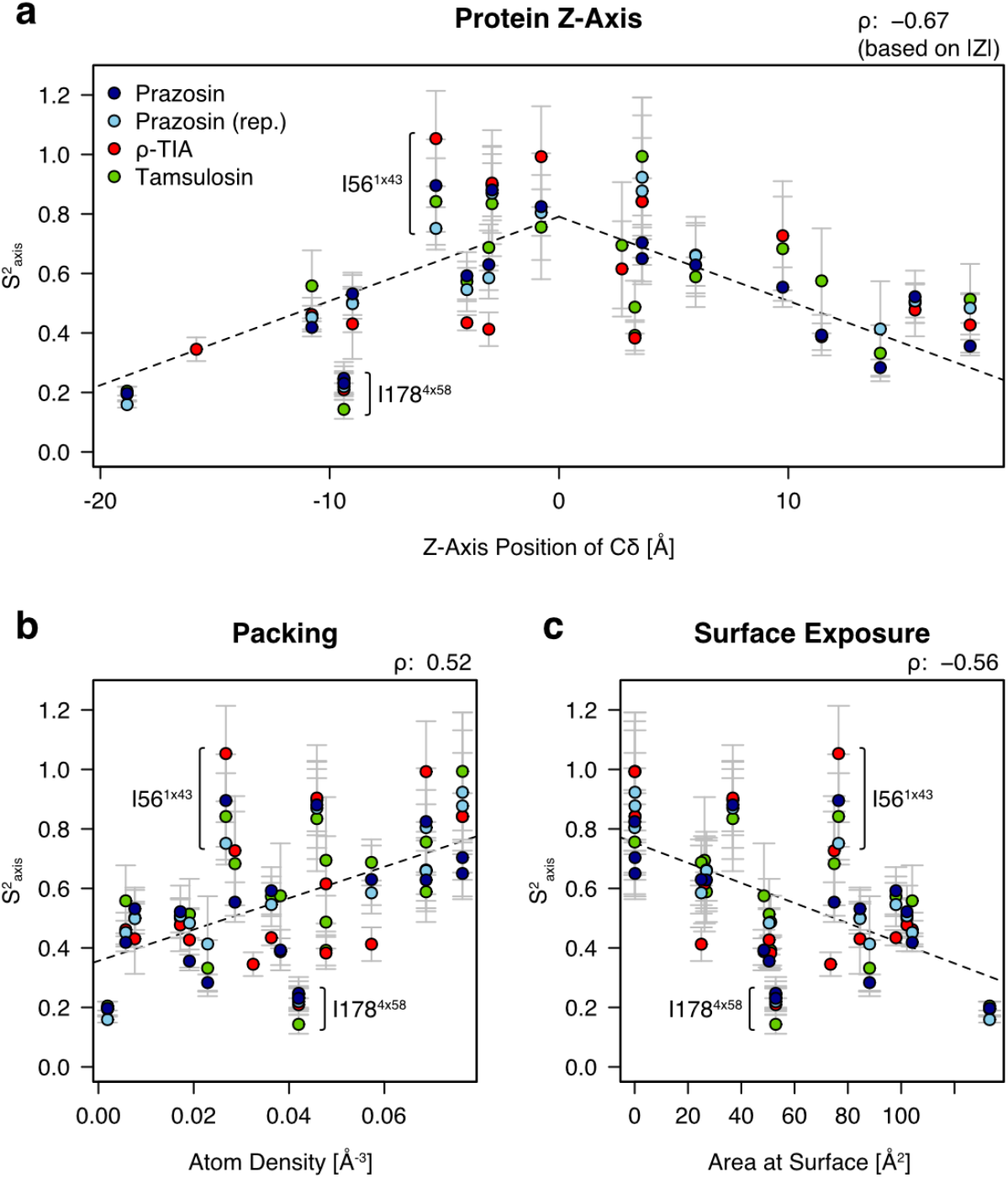
Correlation of Ile δ-methyl order parameters with structural properties of α_1B_-AR. Ile δ-methyl S^2^_axis_ values were correlated to three structural properties of the Ile δ-methyl groups according to the crystal structure of inverse agonist-bound α_1B_-AR (PDB entry 7B6W) (Deluigi et al. 2022). **(a)** Correlation between Ile S^2^_axis_ and the position of δ-methyl groups along the receptor z-axis, as determined by the Cδ position. The receptor center was defined as the average z-coordinate within the α_1B_-AR PDB file. Negative and positive z-coordinates indicate the extracellular and intracellular receptor halves, respectively. **(b)** Correlation between Ile S^2^_axis_ and side-chain packing. **(c)** Correlation between Ile S^2^_axis_ and surface exposed area of the side chain, as quantified using GETAREA (Fraczkiewicz and Braun 1998). Spearman’s rank correlation coefficient (ρ) is given in the upper right corner of each plot. The two residues that deviate the strongest from the general trends (I56^1x43^ and I178^4x58^) are labeled in all plots. Error bars indicate the S^2^_axis_ standard errors. Dashed trendlines are based on linear regression. Correlations for each individual order parameter data set were significant (p-value < 0.05) with the exceptions of S^2^_axis_ to side-chain packing and surface exposure in the ρ-TIA-bound α_1B_-AR-B1D1 (Figure S4.5.2).

The combined Ile δ-methyl order parameters from all data sets correlate moderately with all three above-mentioned structural properties. The strongest correlation of the side-chain dynamics is with the position of the Ile residues along the receptor z-axis (ρ = −0.67). Further, side-chain dynamics correlate with packing (ρ = 0.52) and surface exposure (ρ = −0.56): Side chains that are less densely packed and/or are more solvent- or detergent-exposed tend to undergo larger motions. Correlations of the structural properties of α_1B_-AR bound to (+)-cyclazosin with the individual dynamic data sets of α_1B_-AR-B1D1 bound to prazosin and tamsulosin are stronger than with the dynamics of α_1B_-AR-B1D1 bound to ρ-TIA (Figure S4.5.1). The weaker correlations with the dynamics of α_1B_-AR-B1D1 bound to ρ-TIA indicate subtle receptor-wide changes in side-chain dynamics, which could be due to structural changes caused by the allosteric peptide ρ-TIA compared to the orthosteric small-molecules prazosin and tamsulosin.

The correlations between all combined Ile δ-methyl S^2^_axis_ values and the structural properties of those δ-methyl groups imply that Ile methyl order parameters can be explained to a considerable extent based on the structure alone. For the investigated structural properties, the z-axis position explains 44 %, packing explains 27 %, and surface exposure explains 30 % of the S^2^_axis_ variation (Figure S4.5.3). The extent to which structure contributes to side-chain dynamics seems notably high in this membrane receptor, since side-chain dynamics in soluble proteins have been described as not generally correlated to similar structural properties (Igumenova et al. 2006). Two Ile residues in α_1B_-AR-B1D1 do not follow the above-mentioned general trends: I56^1x43^ appears more rigid, whereas I178^4x58^ is more dynamic than expected based on the currently available α_1B_-AR crystal structure (PDB entry 7B6W). The higher-than-expected rigidity of I56^1x43^ likely reflects the outward tilting of TM1 due to crystal packing in 7B6W, which may result in an overestimation of the side-chain surface exposure and an underestimation of the packing density for I56^1x43^, compared to the structure in solution (Deluigi et al. 2022). In contrast, the higher-than-expected flexibility of I178^4x58^ suggests that high flexibility in this receptor region could be functionally relevant. Notably, a conformational transition is detected by I178^4x58^ in the prazosin- and tamsulosin-bound α_1B_-AR-B1D1 (Figure 3c), the extracellular end of TM4 is important for receptor activation in the closely related α_2C_-AR (Chen et al. 2019), and agonist-induced signaling of α_1B_-AR was abolished by a stabilizing mutation (G183^4x63^V) in this region (Deluigi et al. 2022).

### Differences in methyl order parameters between Ile, Leu, and Val

A comparison of Ile, Leu, and Val S^2^_axis_ values in α_1B_-AR-B1D1 reveals differences in the distributions of their methyl order parameters (Figure 5). Whether a signal belongs to a Leu or a Val methyl group was determined by using reference spectra of α_1B_-AR-B1D1 in which only Val methyl groups were labeled (Figure S2.3) (Mas et al. 2013). Most Val side chains exhibit large order parameters, i.e. rigid behavior (Figure 5c), whereas most Leu side chains exhibit small order parameters, i.e. flexible behavior (Figure 5b). Finally, Ile side chains show a more centered distribution of their S^2^_axis_ values (Figure 5a). These observations raise the question of whether the different distributions of methyl order parameters reflect an intrinsic property of the amino acids or whether the different amino acids are primarily located in dynamically or structurally distinct receptor regions.

**Figure 5.**
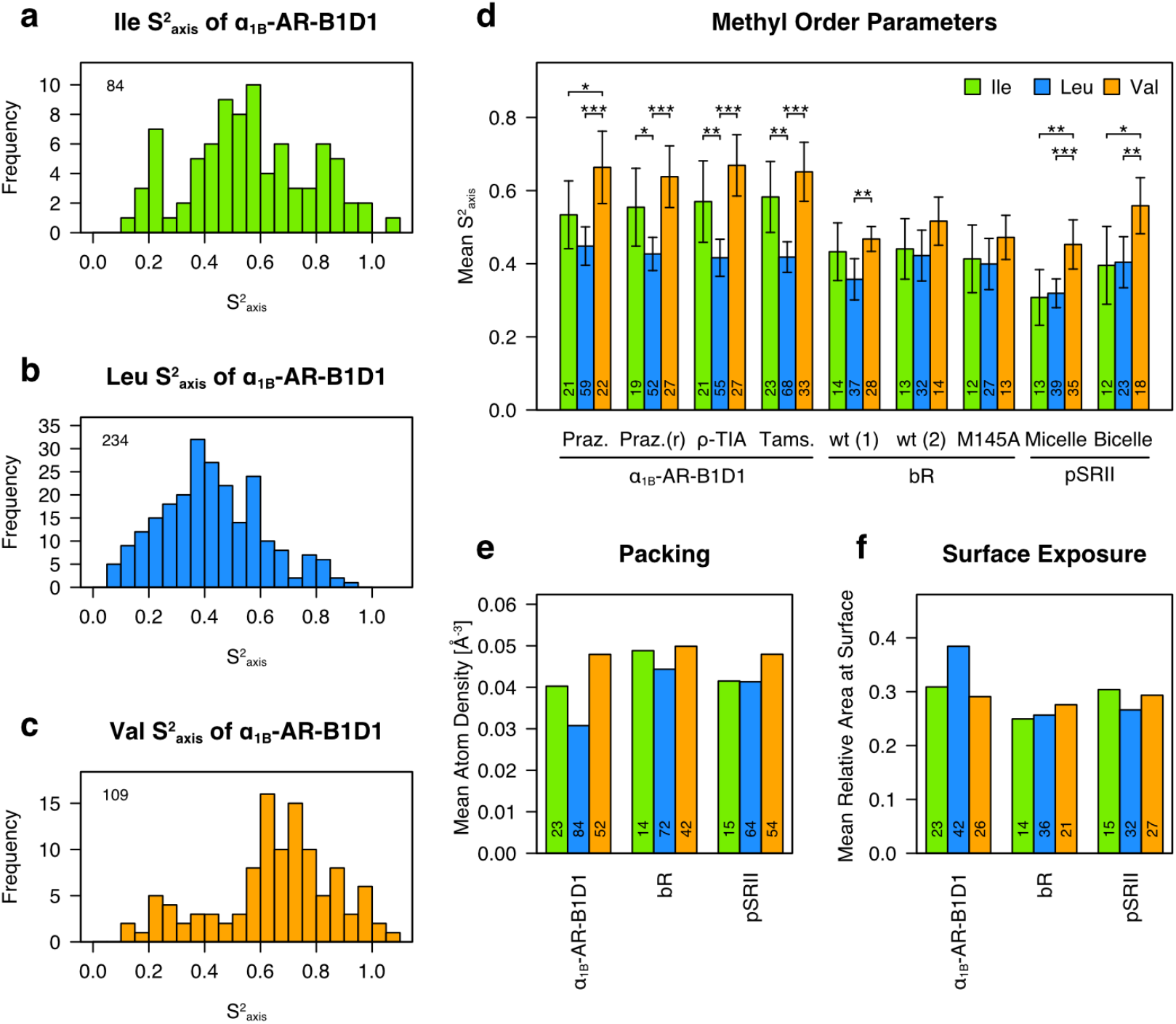
Distribution of methyl order parameters in α_1B_-AR-B1D1 by amino acid type and comparison of dynamics and structural characteristics between α_1B_-AR, bR, and pSRII. **(a–c)** Histograms of all obtained S^2^_axis_ values for α_1B_-AR-B1D1. The total number of S^2^_axis_ values per amino acid is shown in the top left corner. Average Ile, Leu, and Val methyl order parameters including the 95 % confidence interval for the mean. Data for α_1B_-AR-B1D1 bound to the indicated ligands and for the published S^2^_axis_ values of bR (Kooijman et al. 2020b) and pSRII (O’Brien et al. 2020) are included. Significances between order parameters were assessed using the Wilcoxon rank-sum test. Significances are indicated by one (p-value ≤ 0.05), two (p-value ≤ 0.01), and three (p-value ≤ 0.001) stars. The values for bR order parameters were scaled by a factor of 1.17 compared to the published values due to a correction of the rotational correlation time (manuscript in preparation). The numbers indicate the number of order parameters for each amino acid type. **(e)** Side-chain packing based on the structures of α_1B_-AR (PDB entry 7B6W), bR (PDB entry 5ZIM), and pSRII (PDB entry 1H68). Numbers indicate the number of methyl groups for each amino acid type (considering only the δ-group for Ile). **(f)** Protein surface exposure of the side chains relative to the surface of the side chain of the free amino acid. Numbers indicate the number of side chains per amino acid type and protein.

To shed light on this question, we compared the side-chain dynamics of α_1B_-AR-B1D1 with those of the two microbial rhodopsins bacteriorhodopsin (bR) and sensory rhodopsin II (pSRII), whose methyl order parameters were published by Kooijman et al. (Kooijman et al. 2020b) and O’Brien et al. (O’Brien et al. 2020), respectively (Figure 5d). α_1B_-AR, bR, and pSRII share a common architecture of 7 transmembrane helices; however, bR is a light-driven proton pump and pSRII is a photoreceptor for blue light. Val methyl order parameters mostly adopted higher values than Leu or Ile residues in all three proteins, likely related to the shorter Val side chain that limits the extent of possible motions compared to Ile and Leu (Sharp et al. 2014; Wand 2001). The observed difference between the average Ile and Leu S^2^_axis_ for δ-methyl groups in α_1B_-AR-B1D1, however, requires a different explanation as Ile and Leu side chains differ only in the branching position of one of the methyl groups. Further, systematic differences between the δ-methyl order parameters of Ile and Leu side chains are either absent or only very small in bR and pSRII, suggesting that the difference in branching position is unlikely the reason for the difference between mean Ile and Leu side-chain dynamics in α_1B_-AR-B1D1.

Side-chain dynamics correlate with structural properties and are likely partially governed by them, as shown above for Ile side-chain dynamics in α_1B_-AR-B1D1 (Figure 4). The same correlations for packing and surface exposure are present for Ile and Leu in the two microbial rhodopsins as well (Figure S4.6.1 and S4.6.3). A comparison between α_1B_-AR-B1D1 and the two microbial rhodopsins revealed differences in side-chain packing and exposure on the protein surface, which likely account for the observed differences in side-chain dynamics between Ile and Leu in α_1B_-AR-B1D1 (Figure 5e–f): The largest difference is observed for Leu residues, which are packed substantially less tightly in α_1B_-AR than in the two microbial rhodopsins. Further, Leu is substantially less tightly packed than Ile within α_1B_-AR, whereas a similar difference is missing in the microbial rhodopsins. The corresponding trends are reflected in the amount of exposure at the protein surface, with Leu in α_1B_-AR being the most exposed amino acid among all three proteins. These structural observations agree well with the average methyl order parameters of Leu and Ile in the three proteins. Thus, the structural environment in which Leu and Ile side chains are found might explain the observed patterns in their dynamic behavior.

In summary, our analysis indicates that Ile and Leu residues are present in similar structural and dynamical environments in the two microbial rhodopsins, whereas they are distributed in a structurally differentiable manner in α_1B_-AR, suggesting distinct roles for these amino acids. Therein, Ile residues occur more frequently in regions with increased packing and display more rigidity than Leu residues, whose side chains are more protein surface-exposed and undergo increased motions. The difference in side-chain packing and exposure at the protein-surface is common among class A GPCRs and could hint at a role of Leu in adjusting protein hydropathy for correct membrane insertion (Baumann and Zerbe 2023).

## Discussion

Ligand binding and allostery are two crucial facets of GPCR pharmacology and signal transmission that depend heavily on protein dynamics, and those include fast motions within the ps to ns range. Fast dynamics, however, have been so far only poorly characterized for GPCRs due to technical challenges. Mostly conformational changes on slower timescales are studied, for example by NMR using strategically placed ^19^F tags. In one study of the A_2A_R, three different active conformational states were detected at the cytosolic end of TM6 in agonist- and Gα-bound receptor whereas both inactive and active conformational states were present in the inverse agonist-bound receptor (Huang et al. 2021). Similarly, the presence of multiple peaks in the [^13^C,^1^H]-HSQC spectrum of α_1B_-AR-B1D1 for I298^6x39^ imply that this inactive receptor construct also samples distinct conformational states that differ at the intracellular end of TM6 (Figure 6a).

**Figure 6.**
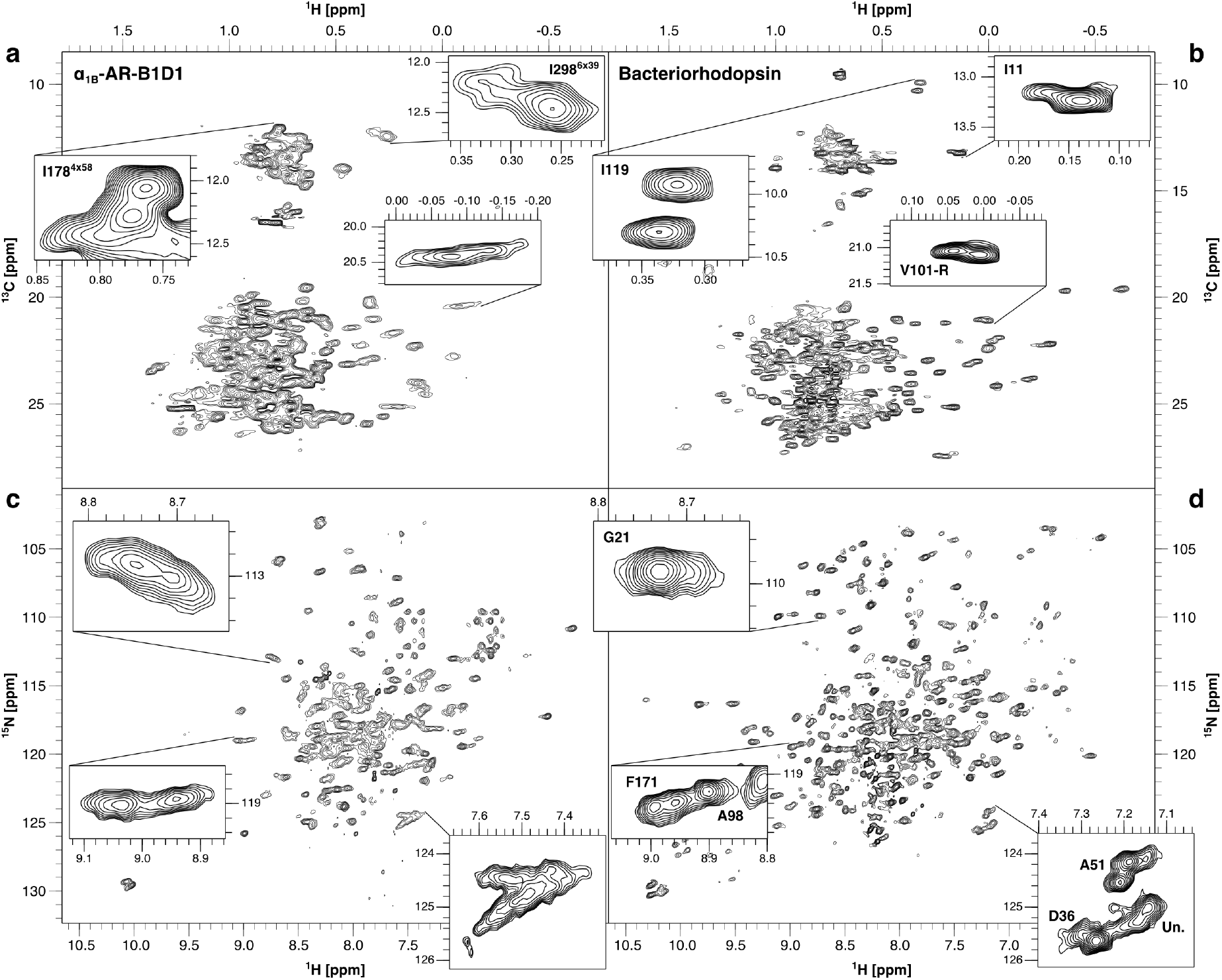
Comparison of spectra of prazosin-bound α_1B_-AR-B1D1 (left) and bR (right). Spectra were recorded at 700 MHz and 320 K. Insets highlight peaks at similar shifts or with similar characteristics between the two proteins. [^13^C,^1^H]-HSQCs of ILV-labeled α_1B_-AR-B1D1 **(a)** and bR **(b)** indicate that methyl groups are generally better resolved in bR. Note that there are fewer ILV residues in bR (Ile: 15, Leu: 38, Val: 21) than in α_1B_-AR-B1D1 (Ile: 24, Leu: 47, Val: 28). ^15^N-TROSYs of uniformly ^15^N-labeled α_1B_-AR-B1D1 **(c)** and bR **(d)** indicate exchange processes in both proteins. Note that signals from the TM portions are largely absent in the spectra of α_1B_-AR-B1D1 due to the missing exchange of deuterons to protons for water-inaccessible amides (Schuster et al. 2020). The overrepresentation of loop amides might make α_1B_-AR-B1D1 appear more dynamic. Rotational correlation times of both proteins are similar (37.07 ns for α_1B_-AR-B1D1 in micelles (138.1 kDa (Schuster et al. 2020)) and 42.58 ns for bR in nanodiscs (127 kDa (Kooijman et al. 2020a))).

In an energy landscape, all these different conformational states (further referred to simply as states) correspond to energy wells separated by substantial energy barriers. Slow dynamics (μs to ms range) typically describe transitions between these deep energy wells and therefore typically correspond to rather large conformational changes, e.g., the rearrangement of a TM helix. In contrast, fast dynamics characterize transitions within a single deep energy well and thus describe transitions between microstates. While ^19^F studies are very successful in characterizing slow dynamics, they provide little information about the dynamics governing transitions between microstates. In contrast, studying the dynamics of methyl groups potentially provides a wealth of information on both slow and fast dynamics. Fast dynamics are especially interesting since they account for most of the conformational entropy of the system, which affects e.g. ligand binding and the coupling of signaling effectors (Hoffmann et al. 2022; Igumenova et al. 2006).

A comparison of methyl spectra of two 7-TM membrane proteins, namely α_1B_-AR-B1D1 and bR, suggests the presence of additional exchange in the ms regime in α_1B_-AR-B1D1, because signals are generally broader than in bR (Figure 6). Dark-adapted bR exists in an equilibrium of two very slowly exchanging states, often resulting in two well-separated methyl signals as also observed for I178^4x58^ in α_1B_-AR-B1D1. Inspection of the ^15^N-TROSY spectrum of α_1B_-AR-B1D1 that reports on the backbone again reveals the presence of a multitude of states that interconvert in the ms regime (single broad peaks) and much more slowly (multiple peaks in close proximity).

Overall, side-chain dynamics are conserved between the different inverse agonist-bound forms of α_1B_ - AR-B1D1 on a global receptor-wide scale. Even though we only tested inverse agonists, these varied in their chemical structures, sizes, receptor subtype selectivities, extent of inverse agonism, and binding modes (Chen et al. 2004; Michel et al. 2020; Proudman et al. 2020). Based on all these differences, one might expect that they would cause at least some receptor-wide differences, impacting overall structural and/or dynamical properties of the receptor. Clark et al. (Clark et al. 2017) demonstrated in the case of agonist- and inverse agonist-bound A_2A_R that it is possible to detect global changes in fast dynamics. Hence, it is likely that no substantial differences in side-chain dynamics exist between the different inverse agonist-bound forms in α_1B_-AR-B1D1. Interestingly, global side-chain dynamics appear to be conserved even in the absence of ligands, implying that inverse agonist binding does not significantly change the conformational landscape of α_1B_-AR-B1D1 in ways that could be detected by side-chain dynamics. Note that, for example, a decrease in the exchange rate between conformational states upon inverse agonist binding would not influence side-chain dynamics.

Even though these measurements were carried out on a stabilized mutant that is signaling-inactive, the same behavior might be expected for wild-type (wt) α_1B_-AR due to the very low basal activity of this receptor (Kjelsberg et al. 1992). The very low basal activity means that wt apo α_1B_-AR mostly populates one or multiple inactive states and rarely transitions to the active state(s). The binding of inverse agonists further stabilizes the inactive state(s), making the active state(s) virtually inaccessible; however, the change in populations is likely small due to the already very low basal activity of the wt receptor.

Similarly to wt α_1B_-AR in its apo and inverse agonist-bound form, the α_1B_-AR-B1D1 construct used in this study is likely locked in its inactive form due to the thermostabilizing mutations (Deluigi et al. 2022). α_1B_-AR-B1D1 might thus be a reasonable model for the study of the inactive state(s), and the similar fast dynamics of apo and inverse agonist-bound α_1B_-AR-B1D1 is therefore consistent with a receptor having a very low basal activity, assuming that only the active state(s) show substantially increased side-chain dynamics. However, it is unknown to which degree the mutations possibly changed the conformational landscape of the inactive α_1B_-AR-B1D1 compared to wt α_1B_-AR. Consequently, the dynamics of a less stabilized, signaling-competent construct will require further investigation in future studies.

Despite the similar fast side-chain dynamics of all investigated α_1B_-AR forms, localized changes were observed for I176^4x56^ and possibly I214^5x49^, which undergo increased dynamics when ρ-TIA is bound compared to when either prazosin or tamsulosin are bound (Figure 7a). The peptide ρ-TIA binds allosterically at the extracellular surface of α_1B_-AR to the extracellular tips of TM6 and TM7, in proximity of the extracellular loop three (Figure 2a) (Ragnarsson et al. 2013). Especially I176^4x56^ is a potentially interesting reporter for relevant allosteric changes, since its side chain points towards TM5 and P215^5x50^ of the PIF motif (Wacker et al. 2013) (Ile-Cδ to Pro-Cγ distance of 6.4 Å). In β_2_ -AR, the residues of the PIF motif (P^5x50^, I^3x40^, F^6x44^) undergo conformational rearrangements upon receptor activation due to the reorientations of TM5 (including P^5x50^) and TM6, which includes a large movement of the F^6x44^ side chain and a rotameric change of I^3x40^ (Rasmussen et al. 2011a; Rasmussen et al. 2011b; Wacker et al. 2013).

**Figure 7.**
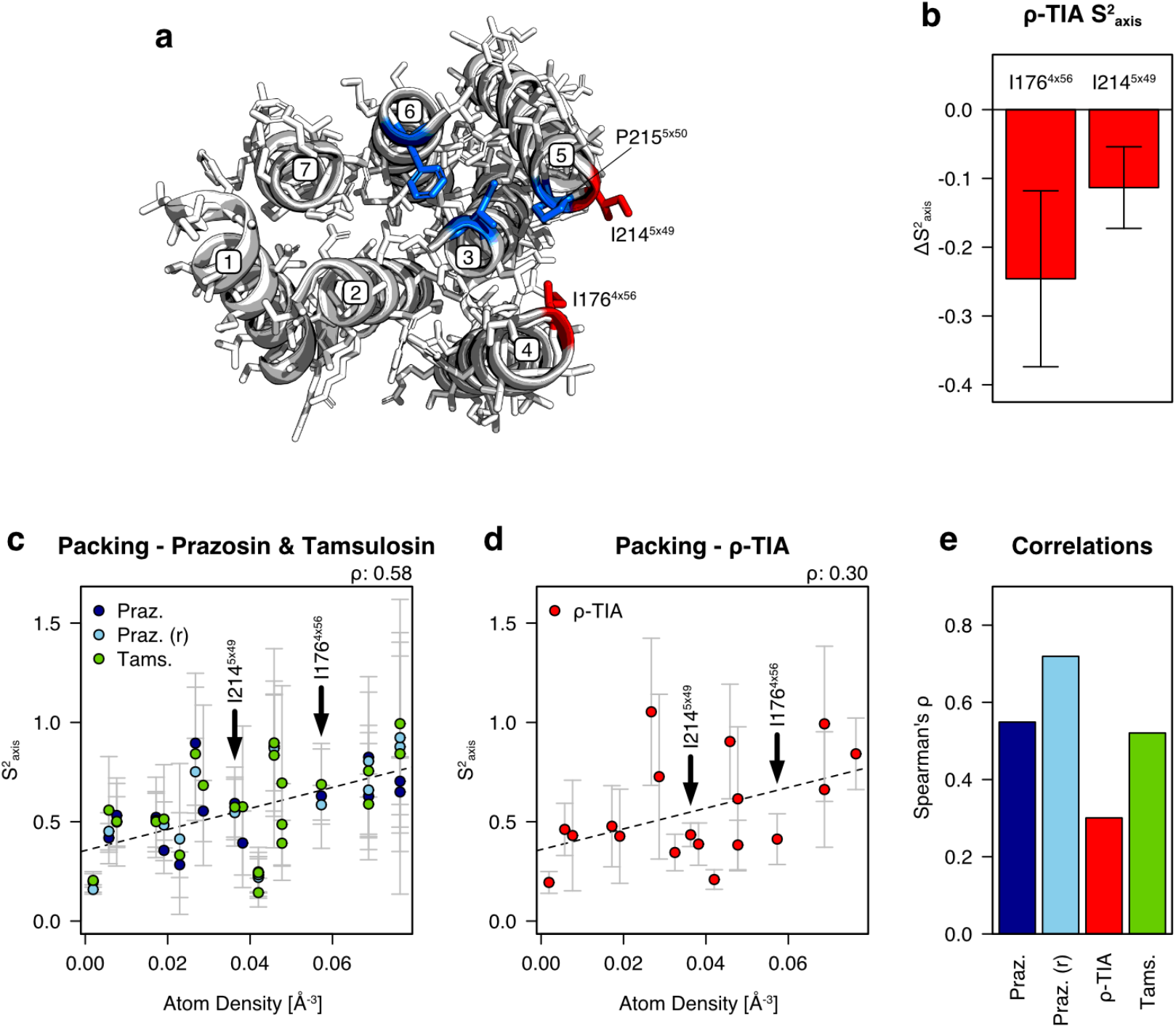
Localized and overall changes in side-chain dynamics in the presence of ρ-TIA. **(a)** α_1B_-AR bound to (+)-cyclazosin (PDB entry 7B6W) viewed from the extracellular side. The extracellular region of the receptor has been omitted for clarity. Ile side chains with increased flexibility in the ρ-TIA-bound α_1B_-AR-B1D1 (I176^4x56^ and I214^5x49^) are highlighted in red, while residues of the PIF motif (P215^5x50^, I133^3x40^, F303^6x44^) are highlighted in blue. **(b)** Difference in experimental and predicted S^2^_axis_ of I176^4x56^ and I214^5x49^ in ρ-TIA-bound α_1B_-AR-B1D1. The predicted values were calculated based on the correlation with packing density in the structure of α_1B_-AR bound to (+)-cyclazosin (PDB entry 7B6W) using a linear model. Negative values indicate a higher flexibility than expected. Error bars indicate the 95 % confidence intervals. **(c)** Correlation between side-chain packing and Ile S^2^_axis_ of prazosin- and tamsulosin-bound α_1B_-AR-B1D1. **(d)** Correlation between side-chain packing and Ile S^2^_axis_ of ρ-TIA -bound α_1B_-AR-B1D1. Error bars indicate the 95 % confidence intervals. **(e)** Spearman’s rank correlation coefficients (ρ) between side-chain packing and individual S^2^_axis_ datasets. Individual correlations for side-chain packing, surface exposure, and protein z-axis are shown in Figure S4.5.1.

Altered side-chain dynamics in this region might thus indicate a potential difference in the inactivation mechanism between the orthosteric and the allosteric ligands. The side chains of both I176^4x56^ and I214^5x49^ switch from the α- to the J-class of motion, implying that they undergo rotameric transitions more frequently in the ρ-TIA-bound receptor (Wand and Sharp 2018). We speculate that the increased dynamics in the presence of ρ-TIA reflect a decrease in packing density between TM4 and TM5. The δ-methyl order parameters of I176^4x56^ and I214^5x49^ in the prazosin- and tamsulosin-bound receptor correspond to the expected values based on the (+)-cyclazosin-bound receptor structure (PDB entry 7B6W) (Figure 7c). In contrast, the δ-methyl order parameters obtained in the presence of ρ-TIA are significantly smaller than expected when using the same structure, suggesting less dense packing in the above-mentioned region when the receptor binds ρ-TIA (Figure 7b and 7d). We want to stress here that I176^4x56^ and I214^5x49^ likely do not capture all changes that occur in the ρ-TIA-bound α_1B_-AR-B1D1 but are merely the side chains that passed the significance threshold. Interestingly, I133^3x40^, which is part of the PIF motif and the transmission switch, shows a large (but due to experimental error non-significant) increase in rigidity in the presence of ρ-TIA (Figure 3a). Further, the poorer correlation of (+)-cyclazosin-bound α_1B_-AR structural properties to the side-chain dynamics of ρ-TIA-bound α_1B_-AR-B1D1 compared to those of either prazosin- or tamsulosin-bound α_1B_-AR-B1D1 implies subtle global changes in dynamics, which might be due to structural changes triggered by the allosteric ligand (Figure 7e). The fact that no changes in global dynamics between the ligands were detected indicates that the changes in dynamics are not substantial enough to alter the overall methyl order parameter distribution or that compensatory changes result in identical overall dynamics.

In conclusion, we have demonstrated the feasibility to determine side-chain dynamics in a GPCR using NMR by methyl relaxation. We characterized fast side-chain dynamics of a stabilized α_1B_-adrenergic receptor construct in presence of different inverse agonists, showing that this methodology is powerful enough to identify localized changes in fast dynamics and also subtle global changes correlated with the structure.

## Methods

### Protein expression and purification

Detailed descriptions of the expression and purification of the α_1B_-AR-B1D1 construct used in this study can be found in Supplementary Data 1 (SD1) in Sections 1.2 and 1.3. α_1B_-AR-B1D1 was optimized for the expression in the inner membrane of *E. coli* cells, and thermostability was enhanced by introducing 13 mutations, truncated N- and C-termini, and a truncation within the intracellular loop 3 (Deluigi et al. 2022; Schuster et al. 2020; Yong et al. 2018) compared to wild-type human α_1B_-AR (UniProt entry P35368). Expression in *E. coli* cells allows for a high degree of deuteration and the use of established labeling schemes for ^13^C,^1^H-labeling of the δ-methyl groups of isoleucine and leucine, and the γ-methyl groups of valine side chains using α-ketoisovaleric and α-ketobutyric acid (Kerfah et al. 2015; Tugarinov et al. 2006; Tugarinov and Kay 2004). We used variants of α-ketoisovaleric and α-ketobutyric acid that only the methyl group carbon, but not other carbons, were labeled to avoid ^13^C,^13^C couplings, resulting in high resolution in the ^13^C dimension. α_1B_-AR-B1D1 was expressed in minimal media using perdeuterated glucose, as previously described (Schuster et al. 2020). After a TALON Superflow purification step, fusion proteins (maltose-binding protein (MBP) and thioredoxin A) were cleaved off from α_1B_-AR-B1D1 by 3C protease treatment. The last step of the purification utilizes a ligand column consisting of an immobilized prazosin derivative to isolate properly folded receptors and remove the cleaved fusion proteins (Deluigi et al. 2022; Egloff et al. 2015). α_1B_-AR-B1D1 was solubilized from the *E. coli* membrane with DDM/CHS (n-dodecyl-β-D-maltoside / cholesteryl hemisuccinate) and purified using LMNG (lauryl maltose neopentyl glycol), leading to micelles in which the receptor construct is stable for several weeks at 47°C, the highest temperature at which experiments were performed. Background deuteration was maximized using LMNG with perdeuterated lipid tails (FB Reagents). Protonated LMNG was exchanged to the deuterated LMNG while the receptor was immobilized on the ligand column using two different protocols (details in SD1 Section 1.3), yielding micelles that contained either 50 % or 95 % deuterated LMNG. The experiments shown here were based on two purifications: the first purification yielded the prazosin-, tamsulosin-, and ρ-TIA-bound receptor samples with micelles containing 50 % deuterated LMNG that were used to record spectra at 320 K. The second, optimized purification yielded the prazosin-bound and apo receptor samples with micelles containing 95 % deuterated LMNG that were used to record spectra at 298 K.

### NMR spectroscopy

Samples were measured in 5 mm Shigemi NMR tubes containing 250 μM α_1B_-AR-B1D1 in 20 mM Na-phosphate, 20 mM NaCl, and 0.01 % (w/v) LMNG at pH 7.0 in 90 % H_2_ O and 10 % D_2_ O. Samples for Ile assignments usually contained 50 to 100 μM α_1B_-AR-B1D1, using the same buffer composition as above. All experiments used gradient-based coherence selection schemes (Keeler et al. 1994). Rotational correlation times (τ_c_) were estimated using a 2D [^15^N,^1^H]-TRACT experiment based on Lee et al. (Lee et al. 2006) with delays of 1, 2, 3, 4, 6, 8, 10, 12, 14, 16, 20, 25, 30, 35, 40, and 50 ms. Data were recorded using a pseudo-3D experiment with 64 scans for prazosin- and 128 scans for ρ-TIA-bound α_1B_-AR-B1D1 at 320 K. TRACT was recorded two times at 298 K, once with the same delays as above and 128 scans, and once with delays of 1, 2, 4, 8, 12, and 24 ms and 512 scans to improve the signal-to-noise ratio for the prazosin-bound receptor at the lower temperature. For measurements of methyl dynamics, we used the proton triple-quantum buildup experiment developed by Tugarinov and coworkers (Sun et al. 2011; Tugarinov et al. 2007). Dynamics at 320 K were recorded for the prazosin-, ρ-TIA-, and tamsulosin-bound α_1B_-AR-B1D1 using 32 scans for the “allowed” and 48 scans for the “forbidden” experiments with delays of 1.04, 2, 4, 8, 16, 24, 32, 40, 60, and 80 ms, with a repeated recording with 8 ms delay to assess variability between measurements. Dynamics at 298 K were measured for the apo and the prazosin-bound α_1B_-AR-B1D1 using delays of 2, 4, 8, 16, 24, 32, and 40 ms with increased scan numbers (48 scans for the “allowed” and 96 scans for the “forbidden” data set) to compensate for the higher loss of signal at the lower temperature. All experiments were recorded on a Bruker AV-Neo 700 MHz spectrometer. Spectra were processed using TopSpin v4.1.1 and analyzed using CcpNmr Analysis v2.5.1 (Vranken et al. 2005).

### Assignments

A total of 18 Ile δ-methyl groups of the prazosin-bound α_1B_-AR-B1D1 were assigned (Figure S2.4) using mutations to either Leu or Val residues (Figure S2.6), with one ambiguity being resolved using a ^13^C-resolved [^1^H,^1^H]-NOESY spectrum (Figure S2.7). Some of these assignments were directly transferred to the tamsulosin- and the ρ-TIA-bound receptor, whereas additional mutants were made to assign additional signals, resulting in 19 assigned Ile residues with tamsulosin bound, and 19 assigned Ile residues with ρ-TIA bound (Figure S2.4, S2.8, S2.9). By supplementing unlabeled Leu, the pathway leading from α-ketoisovaleric acid to Leu can be suppressed, which allowed us to distinguish whether signals originated from Leu or Val methyl groups in [^13^C,^1^H]-HSQC spectra (Figure S2.3) (Mas et al. 2013).

### Correlation time estimation

Rates were determined by fitting exponential decays to the intensity data using R v4.0.3 (Team 2019) with the minpack.lm package (Elzhov et al. 2016). The rotational correlation times (τ_c_) were calculated for each signal in the [^15^N,^1^H]-TRACT spectra based on the algebraic solution presented by Robson et al. (Robson et al. 2021) using an amide S^2^_axis_ of 0.9 to improve the accuracy of τ_c_ as they suggested (equations are shown in the SD1 in Section 1.4). The correlation times of the proteins were calculated by taking the mean of the individually calculated τ_c_ values. Values were excluded from this calculation when they belonged to side chains, were smaller than 20 ns, or had standard errors larger than 12 ns. The resulting τ_c_ values at 320 K were 37.1 ns ± 1.3 ns (95 % CI) for the prazosin- and 36.5 ns ± 1.5 ns (95 % CI) for the ρ-TIA-bound receptor. The two values do not differ significantly (Welch’s t-test p-value: 0.574), although the conopeptide is expected to slightly increase the size and thus the rotational correlation time of the complex. The correlation time of 37.1 ns, as estimated with prazosin-bound α_1B_ - AR-B1D1, was used to calculate the methyl order parameters for all three ligands at 320 K. We attempted to measure the rotational correlation time also at 298 K; however, it was not possible to obtain reliable values. We thus compared the prazosin-bound and the apo receptor at 298 K based on their relative side-chain dynamic values (using η instead of S^2^_axis_ values). An extended discussion of the correlation time estimation and the tables with the obtained values can be found in the SD1 in Section 3 and in the SD2 in Tables S12 and S13. Briefly, τ_c_ values from residue-based TRACT experiments are affected by the presence of additional dynamics and by the differences in chemical shift anisotropy (CSA). Neglecting the CSA dispersion and factoring it all into backbone dynamics carries the risk of overestimating τ_c_ and thereby underestimating order parameters. Unfortunately, these two additional contributions cannot be distinguished in TRACT data. We noticed that the distribution of τ_c_ values above a 20 ns agreed with a distribution that is expected based on differences in CSAs alone. The mean of this distribution was therefore taken as the true τ_c_ of the receptor instead of the upper values as done before in Kooijman et al. (Kooijman et al. 2020b) . We used a BEST version of the TRACT experiment, which possibly led to a systematic underestimation of R_β_ (anti-TROSY R_2_). This in turn might have led to an underestimation of τ_c_ and thus an overestimation of methyl order parameters by about 10 % as based on a ubiquitin reference. Such a systematic error would affect only the absolute size (i.e. scaling) of S^2^_axis_ values. Therefore, the banding of order parameters and all analyses that compare α_1B_-AR-B1D1 S^2^_axis_ values with α_1B_-AR-B1D1 S^2^_axis_ values would remain valid. Further, α_1B_-AR-B1D1 would remain more rigid than bR (Kooijman et al. 2020b) and pSRII (O’Brien et al. 2020) if S^2^_axis_ values were 10 % smaller. However, the differences in mean S^2^_axis_ would become less pronounced, especially when compared to pSRII in bicelles.

### Order parameter calculation

Methyl order parameters for prazosin-, tamsulosin-, and ρ-TIA-bound α_1B_-AR-B1D1 were calculated according to Sun, Kay and Tugarinov (Sun et al. 2011):

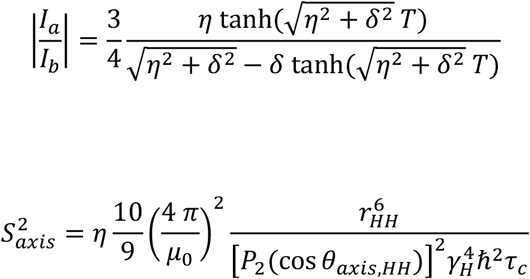

with

μ_0_ = 4 π × 10^−7^ H/m

*ħ* = 6.62607015 × 10^−34^ (2 π)^−1^ J s

r_HH_ = 1.813 × 10^−10^ m

θ_axis,HH_ = 90°

γ_H_ = 267.52218744 × 10^6^ rad s^−1^ T^−1^

where μ_0_ is the vacuum permeability, *ħ* the reduced Planck constant, r_HH_ the distance between methyl protons and γ_H_ the gyromagnetic ratio of protons. θ_axis,HH_ is the angle formed by the vectors between two methyl protons and between the methyl carbon and the adjacent carbon. The values for η and δ were fitted based on the intensity ratios I_a_ /I_b_, where I_a_ is the intensity of signals from the “forbidden” experiment in which triple-quantum proton coherences are formed, whereas I_b_ refers to the “allowed” experiment in which single-quantum proton coherences relax, using R v4.0.3 (Team 2019) with the minpack.lm package (Elzhov et al. 2016). The function P_2_ refers to the second Legendre polynomial. In some cases, the volumes of weaker peaks became strongly influenced by noise, especially at longer delays. To remove outliers in an automated procedure, the fits for the ratios were performed in the absence of one and two data points whenever the residual standard error (RSE) was larger than 0.035 (for measurements at 320 K) or 0.075 (for measurements at 298 K). If the removal of a single datum decreased the RSE to less than 75 % of the original value, then this data point was regarded as an outlier and removed. Two data points were removed whenever this led to a decrease of the RSE to 50 % or less compared to the RSE of the best solution with a single data point removed. Only values obtained from fits that led to standard errors (SEs) for η values below 50 s^−1^ (320 K) or 75 s^−1^ (298 K), to SEs for δ values below 50 s^−1^ (320 K) or 100 s^−1^ (298 K) and with RSEs smaller than 0.065 (320 K) or 0.075 (298 K) were considered for further analysis. Dynamic data with values that correspond to the limits of the fit (0 and 400 s^−1^ for η; 0 and -400 s^−1^ for δ) were removed from the data sets. Some manual adjustments were made to remove additional unreliable S^2^_axis_ values (6 in total) and to include others when they still appeared reliable enough despite not fulfilling one of the criteria (4 in total). The fitted η values were used to compare side-chain dynamics between the apo and the prazosin-bound receptor at 298K. Methyl order parameters were calculated for the data recorded at 320 K using the η values and the obtained τ_c_ of 37.07 ns with a SE of 0.66 ns. S^2^_axis_ standard errors were estimated based on the η SE and the τ_c_ SE using 1 000 Monte Carlo samplings.

About 100 methyl order parameters were obtained per sample, corresponding to 57 % of the 174 methyl groups that were labeled. The highest number of reliable values were obtained for the tamsulosin-bound receptor with a total of 125 methyl order parameters. Most complete data sets were obtained for Ile residues with 19 to 23 out of 24 maximally expected methyl order parameters (79 – 96 %), followed by Leu residues with 52 to 68 out of 94 possible values (55 – 72 %) and by Val residues with 22 to 33 out of 56 possible values (39 – 59 %). Methyl order parameters of the α_1B_-AR-B1D1 indicate the presence of very rigid side chains, including a small number of order parameters larger than one, raising the question of whether the reported values are accurate. Too large S^2^_axis_ values might be best explained by the presence of large errors, e.g. I56^1x43^ of the ρ-TIA-bound receptor (Figure 3a). Due to the slower reorientations of rigid side chains and the resulting less favorable NMR properties, large order parameters are generally obtained at a lower precision. However, we cannot exclude a systematic error due to the TRACT version used (see above).

### Banding of order parameters

The assignment of order parameters into discrete motional classes assumes that individual order parameters represent values that are statistically distributed around a mean value within each class. However, many proteins fail to show clear divisions of order parameters into distinct classes (Best et al. 2004). In the case of the α_1B_-AR-B1D1, k-means clustering with 4 centers was generally the best way to group different combinations of the data, as assessed with the average silhouette method and the likelihood function as used by Sharp et al. (Sharp et al. 2014) in equation 2, which assumes normally distributed order parameters within each band. Four bands were optimal or among the best solutions for all individual data sets (except for tamsulosin), when all the data sets were combined and when all Ile δ-methyl order parameters were combined, with centers being set at very similar positions. Clustering the complete data set led to centers at 0.217, 0.425, 0.625, and 0.853, which is in agreement with previously described band positions for J’-, J-, α-, and ω-bands (Kooijman et al. 2020b; O’Brien et al. 2020; Sharp et al. 2014), corresponding to borders at 0.32 (J’ to J), 0.53 (J to α), and 0.74 (α to ω). The four motional classes contained 21.0, 33.9, 27.6, and 17.5 % of the overall order parameter population, respectively. Interestingly, the deviation from this banding is quite large when only Leu or Val residues are considered, which might imply that individual amino acids needed to be considered separately. Clustering and analyses were done with R v4.0.3 (Team 2019) using the cluster package (Maechler et al.).

### Structural analyses

The published α_1B_-AR structure bound to the inverse agonist (+)-cyclazosin (PDB entry 7B6W) (Deluigi et al. 2022) was used to map Ile δ-methyl order parameters on the structure and to extract structural features. The structures of bR (PDB entry 5ZIM) and pSRII (PDB entry 1H68) were used analogously. Missing atoms in the three structures were modeled (using the mutagenesis tool), and protons were added in PyMol v2.4.2. The distances to the α_1B_-AR central plane perpendicular to the membrane plane were calculated with respect to the Ile Cδ-positions. The α_1B_-AR central plane was determined as the average z-coordinate of all atoms within the α_1B_-AR based on PDB entry 7B6W. Side-chain packing densities were calculated by counting the number of atoms belonging to the protein within a 5 Å radius around the δ-methyl carbons of Ile and Leu and the γ-methyl carbons of Val residues and dividing by the spherical volume. Atoms that belong to the same residue as the probed methyl group were not included in the count. Calculations were carried out in R v4.0.3 (Team 2019) using the bio3d (Grant et al. 2021) package. The area of the side chains at the protein surface was quantified by determining the solvent-accessible surface area using GETAREA (Fraczkiewicz and Braun 1998) using default settings with a water probe radius of 1.4 Å. The relative area of the side chains at the protein surface was calculated using the mean surface values of the free amino acid side chains (174.2 Å^2^ for Ile, 174.0 Å^2^ for Leu, and 147.3 Å^2^ for Val residues). If not mentioned otherwise, all calculations, statistical analyses, and plots were carried out using R v4.0.3 (Team 2019) with RStudio v1.4.1103 (Team 2021) and the package stringr (Wickham 2019). Structures were plotted using PyMol v2.4.2.

## Data availability

Ile assignments, side-chain dynamics and TRACT data of α_1B_-AR-B1D1 are listed in the Supplementary Data 2: Table S1 (Ile assignment with prazosin at 320 K), Table S2 (Ile assignment with ρ-TIA at 320 K), Table S3 (Ile assignment with tamsulosin at 320 K), Table S4 (Ile assignment with prazosin at 298 K), Table S5 (Ile assignment of apo α_1B_-AR-B1D1 at 298 K), Tables S6 and S7 (order parameters with prazosin at 320 K), Table S8 (order parameters with ρ-TIA at 320 K), Table S9 (order parameters with tamsulosin at 320 K), Table S10 (side-chain dynamics with prazosin at 298 K), Table S11 (side-chain dynamics of apo α_1B_-AR-B1D1 at 298 K), Table S12 (TRACT data with prazosin at 320K), Table S13 (TRACT data with ρ-TIA at 320K).

Ile assignments and order parameters of α_1B_-AR-B1D1 are also deposited on the BMRB under the following IDs: 51922 (with prazosin at 320 K), 51923 (with tamsulosin at 320 K), 51924 (with ρ-TIA at 320 K), 51925 (with prazosin at 298 K), and 51926 (apo at 298 K).

## CRediT authorship contribution statement

**Christian Baumann:** Conceptualization, Investigation, Formal analysis, Writing – Original Draft, Visualization, Data Curation, Funding acquisition **Wan-Chin Chiang:** Investigation, Visualization **Renato Valsecchi:** Investigation **Simon Jurt:** Methodology, Investigation **Mattia Deluigi:** Writing – Review & Editing **Matthias Schuster:** Conceptualization **Andreas Plückthun:** Conceptualization, Writing – Review & Editing **Oliver Zerbe:** Conceptualization, Investigation, Writing – Original Draft, Supervision, Funding acquisition

## Acknowledgements

This project was supported by research grants from the Swiss National Science Foundation (grants No. 310030-197679 and 310030-159453) and the Forschungskredit of the University of Zurich. (grant FK-18-083). We thank D. Scott and P. Gooley for providing samples of ρ-TIA.

