## Supplementary Data 1 for "Side-Chain Dynamics of the α1B-Adrenergic Receptor determined by NMR via Methyl Relaxation"

**Side-Chain Dynamics of the $\alpha_{1B}$-Adrenergic Receptor determined by NMR via Methyl Relaxation**

Christian Baumann, Wan-Chin Chiang, Renato Valsecchi, Simon Jurt, Mattia Deluigi, Matthias Schuster, Andreas Plückthun, Oliver Zerbe

**Index:**

1. Detailed Methods 2
    1. Thermostabilized $\alpha_{1B}$-AR Construct 2
    2. Expression of $\alpha_{1B}$-AR-B1D1 3
    3. Purification of $\alpha_{1B}$-AR-B1D1 4
    4. TRACT 7
2. Spectra and Assignments 8
3. Correlation Time Estimation using TRACT 13
4. Side-Chain Dynamics 16
    1. $\alpha_{1B}$-AR-B1D1 Methyl Dynamics Data 16
    2. Comparisons Between $S^2_{axis}$ Distributions 17
    3. Side-Chain Dynamics at 298 K 18
    4. Ligand-Dependent Differences in Ile $S^2_{axis}$ 20
    5. Correlations with $\alpha_{1B}$-AR-B1D1 Ile $S^2_{axis}$ 23
    6. Correlations with microbial rhodopsin Ile and Leu $S^2_{axis}$ 26

### 1. Detailed Methods

#### 1.1 Thermostabilized $\alpha_{1B}$-AR Construct

The thermostabilized $\alpha_{1B}$-AR variant $\alpha_{1B}$-AR-B1D1 and the protocols for expression and purification were established in previous studies (Deluigi et al. 2022; Schuster et al. 2020; Yong et al. 2018). The stability of the wild-type $\alpha_{1B}$-AR was improved using CHESS (Scott and Plückthun 2013) in the presence of a fluorescently labeled prazosin derivative and by truncating the termini and the intracellular loop 3 (ICL3), resulting in a construct with 13 mutations:

**$\alpha_{1B}$-AR-B1D1:**

```
             40          50          60          70
 GPGSSTLPQ  LDITRAISVG  LVLGAFILFA  IVGNILVILS  VACNRHLRTP

80          90          100         110         120
TNYFIVNLAM  ADLLLCFTVL  PFSAALEVLG  YWVLGRTFCD  IWAAMDVLCC
                 1                        2            3

130         140         150         160         170
TASILSLCAI  SIDRYIGVRY  YLQYPTLVTR  RKAILALLCV  WVLSTVISIG
                        4                    5

180         190         200         210         220
PLLVWKEPAP  NYDKVCGVTE  EPFYALFSSL  GSFYIPLAVI  LVMYCRVYIV
   6         7  8

230        |285         295         305         315
AKRTTKNLEA|KFSREKKAAK  MLGIVVGMFI  LCWLPFFIAL  PLGSLFSTLK
           |            9

325         335         345         355         365         375
PPDAVFKVLL  WLGYFNSCLN  PIIYLCFSKE  FKRAFVRILG  CQCRGTRELE  VLFQ
       10,11                12  13
```

```
Mutations:
M1:     S95C   2x54
M2:     I116T  3x23
M3:     V124M  3x31
M4:     S150Y  ICL2
M5:     S168C  4x48
M6:     G183V  4x63
M7:     D191Y  ECL2
M8:     E194V  ECL2
M9:     T295M  6x36
M10:    V333L  7x37
M11:    F334L  7x38
M12:    P349L  7x54
M13:    S351F  7x66
```

The sequence numbering in $\alpha_{1B}$-AR-B1D1 is according to the wild-type $\alpha_{1B}$-AR sequence. Mutations are numbered from 1 to 13 based on their occurrence in the amino acid sequence. Helices are highlighted in orange, and mutations are marked with red boxes. Residues in grey boxes were introduced into this construct and are not present in the wild-type receptor. Due to the truncation of the intracellular loop 3, A239 is followed by K285. Positions of mutations are given by GPCRdb numbering (Isberg et al. 2015).

#### 1.2 Expression of $\alpha_{1B}$-AR-B1D1

The stabilized variant $\alpha_{1B}$-AR-B1D1 was expressed as a fusion to an N-terminal maltose-binding protein (MBP) and a C-terminal thioredoxin A (TrxA) in *E. coli* BL21 cells (C2530H, New England Biolabs) using the pRG (a pBR322-derived) vector (Egloff et al. 2015; Egloff et al. 2014; Schuster et al. 2020). Transformed *E. coli* cells were plated on LB-agar plates that contained 1% D-(+)-glucose and 100 mg/l ampicillin. Single colonies were selected to grow precultures in 5 ml M9 $H_2O$ media (3 g/l $KH_2PO_4$, 7.5 g/l $Na_2HPO_4 \cdot 2\ H_2O$, 0.5 g/l NaCl, 2 mM $MgSO_4$, 100 mg/l ampicillin, 1 ml/l 1000x trace metals (ingredients are shown below), 1.5 g/l $NH_4Cl$, 5 g/l D-(+)-glucose). The bacteria were adapted to $D_2O$ using 2 – 3 additional precultures with increasing content of M9 $D_2O$ media (99.8% $D_2O$, 3 g/l $KH_2PO_4$, 6 g/l $Na_2HPO_4$, 0.5 g/l NaCl, 2 mM $MgSO_4$, 100 mg/l ampicillin, 1 ml/l 1000x trace metals, 1.2 g/l $^{15}NH_4Cl$, 3.5 g/l D-(+)-glucose). M9 $D_2O$ media composition for precultures and expression cultures were identical. The expression cultures grew in baffled 2 liter Erlenmeyer flasks. Expression was induced at an $OD_{600}$ of 0.8 – 1.0 with 0.5 mM IPTG and continued for 14 – 20 h at 30°C.

**Trace Metals (1000x) in $H_2O$:**

| | |
|---|---|
| HCl 32% | 50 m/l |
| $FeSO_4 \cdot 7\ H_2O$ | 7 g/l |
| $CaCl_2 \cdot 2\ H_2O$ | 184 mg/l |
| $H_3BO_3$ | 64 mg/l |
| $CoCl_2 \cdot 6\ H_2O$ | 18 mg/l |
| $CuCl_2 \cdot 2\ H_2O$ | 4 mg/l |
| $ZnCl_2$ | 340 mg/l |
| $Na_2MoO_4 \cdot 2\ H_2O$ | 604 mg/l |
| $MnCl_2 \cdot 4\ H_2O$ | 40 mg/l |

In $\alpha_{1B}$-AR-B1D1 produced for dynamics experiments, methyl groups of Ile, Leu, and Val residues were labeled using 70 mg/l α-ketobutyric acid (methyl-$^{13}C$, 99%; 3,3-$D_2$, 98%; Cambridge Isotope Laboratories (CIL)) and 160 mg/l α-ketoisovaleric acid (3-methyl-$^{13}C$, 99%; 3,4,4,4-$D_4$, 98%; CIL). Precursors were added 1 h before induction (at an $OD_{600}$ of 0.6). The M9 $D_2O$ media for methyl group labeling contained 3.5 g/l D-(+)-glucose-1,2,3,4,5,6,6-$d_7$ with an otherwise identical composition as mentioned above. The first batch (yielding $\alpha_{1B}$-AR-B1D1 with prazosin, tamsulosin, and ρ-TIA for dynamics experiments at 320 K) was expressed in 5.25 liters of M9 $D_2O$ media. The second batch (yielding $\alpha_{1B}$-AR-B1D1 with prazosin and in the apo form for dynamics experiments at 298 K) was expressed in 4.2 liters of M9 $D_2O$ media. The second batch was expressed with deuterated $^{15}NH_4Cl$ to further reduce protonation. $^{15}ND_4Cl$ was prepared by dissolving $^{15}NH_4Cl$ in $D_2O$ (10 ml/g) and subsequent lyophilization.

$\alpha_{1B}$-AR-B1D1 constructs produced to assign Ile δ-methyl groups were labeled using 70 mg/l α-ketobutyric acid, which was added 1 h before induction. The M9 $D_2O$ media were prepared with >94% $D_2O$ (recycled in-house) and contained 3.5 g/l of protonated D-(+)-glucose. Typically, expression cultures with 350 ml M9 medium were used to produce individual samples. Ile methyl groups were assigned by mutating each of the 24 Ile residues to either Leu or Val: I42$^{1x29}$L, I46$^{1x33}$L, I56$^{1x43}$V, I60$^{1x47}$L, I64$^{1x51}$L, I67$^{1x54}$V, I84$^{2x43}$V, I120$^{3x27}$L, I133$^{3x40}$V, I139$^{3x46}$L, I141$^{3x48}$V, I145$^{3x52}$V, I163$^{4x43}$V, I176$^{4x56}$L, I178$^{4x58}$L, I214$^{5x49}$L, I219$^{5x54}$L, I228$^{5x63}$V, I298$^{6x39}$V, I304$^{6x45}$L, I312$^{6x53}$L, I346$^{7x51}$L, I347$^{7x52}$V, and I362$^{8x57}$L. Spectra are shown in Figure S2.6, S2.8 and S2.9.

Val methyl groups were selectively labeled based on the protocol of Mas et al. (Mas et al. 2013) to determine whether signals belong to Leu or Val methyl groups. Selective Val methyl labeling was achieved by adding unlabeled Leu (80 mg/l) together with α-ketoisovaleric acid (160 mg/l) to the expression culture 1 h before induction. Samples were produced using expression volumes of 600 and 700 ml. Spectra are shown in Figure S2.3.

##### 1.3 Purification of $\alpha_{1B}$-AR-B1D1

Purifications were carried out at 4°C with precooled buffers. *E. coli* cells were resuspended in resuspension buffer (100 mM HEPES pH 8 at 4°C, 20% glycerol, 400 mM NaCl) using 1.8 to 2.2 ml buffer per gram of cells. Lysozyme (Roth; 10 mg/g of cells) and DNase I (Roche; 0.2 mg/g of cells) were dissolved in 1.8 – 2.2 ml lysis buffer (50 mM HEPES pH 8 at 4°C, 10% glycerol, 200 mM NaCl, 15 mM $MgCl_2$) per gram of cells and added to the cell suspension. The receptor was solubilized using n-dodecyl-β-D-maltopyranoside (DDM, Anatrace (D310)) and cholesteryl hemisuccinate (CHS, Sigma-Aldrich). Solubilized DDM/CHS (10% / 1% (w/v)) was added to yield a final concentration of 2% / 0.2% (w/v) in the cell suspension. Cell lysis and receptor solubilization took place over 2.5 h under constant gentle stirring or rolling. The lysate was centrifuged at 18 000 rpm in an SS34 rotor for 30 min. PD-10 columns (1 column for approximately 3 g of cells) were packed with 2.5 ml of TALON Superflow resin (GE Healthcare) and equilibrated with TALON wash buffer 1 (TWB1; 25 mM HEPES pH 8 at 4°C, 10% glycerol, 600 mM NaCl, 0.075% lauryl maltose neopentyl glycol (LMNG, Anatrace (NG310)), 15 mM imidazole). The supernatant of the lyzed cells was added to the TALON Superflow resin in 50 ml centrifuge tubes with additional imidazole (corresponding to 15 mM imidazole in the supernatant volume) and incubated for 2.5 h while rolling. The TALON Superflow resin suspension was transferred back to the PD-10 columns and washed 4x with 10 ml TWB1 and 4x with 10 ml TWB2 (25 mM HEPES pH 7 at 4°C, 10% glycerol, 150 mM NaCl, 0.05% LMNG). 2.5 ml TALON elution buffer (TEB; 25 mM HEPES pH 8 at 4°C, 10% glycerol, 150 mM NaCl, 0.05% LMNG, 200 mM EDTA pH 8) was added to each of the closed PD-10 columns. The TALON Superflow resin was resuspended, and the suspension was incubated for 5 min before elution. The receptor construct was further eluted using 2x 2.5 ml TEB. The receptor construct was eluted with EDTA instead of imidazole, since imidazole binds weakly to $\alpha_{1B}$-AR-B1D1, which would interfere with the subsequent ligand-affinity purification step. The fusion proteins MBP and TrxA were cleaved off using 3C protease (1 mg 3C protease with 0.075% LMNG (w/v) per 15 ml of TALON Superflow elution; 3C protease was produced in-house (Schuster et al. 2020)) during an incubation period of 1 h.

$\alpha_{1B}$-AR-B1D1 was separated from misfolded receptor molecules and from other contaminants using a resin carrying an immobilized prazosin derivative (Deluigi et al. 2022). This prazosin column (PC) resin (about 1 ml thereof per 20 ml of the TALON Superflow elution) was equilibrated with TEB and added to the TALON Superflow elution. Receptor binding took place overnight while rolling. The PC resin was transferred to one or multiple PD-10 columns and washed 4x with 3 column volumes (CVs) of PC wash buffer (PCWB; 25 mM HEPES pH 8 at 4°C, 10% glycerol, 600 mM NaCl, 0.025% LMNG, 200 mM EDTA pH 8), followed by washing 4x with 3 CVs of PC buffer (PCB; 20 mM Na-phosphate pH 7 at room temperature (RT), 20 mM NaCl, 0.01% LMNG). The receptor was eluted either directly with prazosin or, to obtain the apo receptor, with imidazole. All samples for the side-chain dynamics experiments were purified via the apo state, including the prazosin samples. For the elution with prazosin, 6 CVs of PC prazosin elution buffer (PCPEB; 20 mM Na-phosphate pH 7 at RT, 20 mM NaCl, 0.01% LMNG, 85 µM prazosin) were added to the PC resin and incubated for 2 h while rolling before eluting. The receptor was eluted further using 2x 2 CVs PCPEB. For the elution with imidazole, 5 CVs of PC imidazole elution buffer (PCIEB; 20 mM Na-phosphate pH 7 at RT, 20 mM NaCl, 0.01% LMNG, 350 mM imidazole) were added to the PC resin and incubated for 1 h while rolling before eluting. This step was repeated once, followed by further elution with 3x 1 CV of PCIEB. The PC elution was concentrated using an Amicon Ultra-15 with a 50 kDa cut-off and the buffer was exchanged to PCB with a PD-10 desalting column (GE Healthcare). Residual imidazole was removed by overnight dialysis against 500 ml PCB using a 5 ml Float-A-Lyzer device (Spectrum) with a cut-off of 8 – 10 kDa (or alternatively by a second PD-10 desalting column step).

The protein concentration of the apo receptor sample was determined using a Nanodrop spectrometer at a wavelength of 280 nm using an extinction coefficient of 52 370 $M^{-1}$ $cm^{-1}$. If the protein was eluted with prazosin directly, a 0.5 ml Zeba Spin desalting column with a 7 kDa cut-off (Thermo Fisher Scientific) equilibrated with Zeba buffer (10 mM Na-phosphate pH 7 at RT, 100 mM NaCl, 0.01% LMNG, 2.5 µM prazosin) was used to remove excess prazosin. Measured protein concentrations were multiplied by an empirically established correction factor of 0.7 to correct for the absorption of receptor-bound prazosin. Protein yields were typically in the range of 1.0 – 1.5 mg/l of cell culture, with purifications via imidazole elution from PC generally resulting in lower yields than purifications via prazosin elution. Samples of apo $\alpha_{1B}$-AR-B1D1 were concentrated using an Amicon Ultra-4 with a 50 kDa cut-off before ligands and 10% $D_2O$ were added. The concentrations of added ligands corresponded to two times the receptor concentrations. To adjust for the dilution of the buffer components (20 mM Na-phosphate pH 7 at RT, 20 mM NaCl, 0.01% LMNG) due to the added ligands and $D_2O$, corresponding amounts of 10x PCB were added to the samples. The following ligands were used for the samples that were measured: prazosin hydrochloride (Sigma; 1 mM stock in $H_2O$), tamsulosin hydrochloride (Sigma; 10 mM stock in $H_2O$), and ρ-TIA (10 mM stock in $H_2O$). 5 mm Shigemi NMR tubes with sample volumes of about 220 to 250 µl were used for NMR experiments. Protein concentrations were about 250 µM for dynamics experiments and approximately 50 to 100 µM for methyl assignments.

The background protonation of the samples intended for measuring side-chain dynamics was reduced by using deuterated LMNG ($^2$H-LMNG) (FB Reagents) that has 98% of its 42 protons in the aliphatic tail replaced by deuterons with the residual 2% protons found at the enolizable position. Deuterated instead of protonated LMNG was used in PCWB, PCB, and PCIEB to enable the exchange of the protonated to the deuterated detergent in the micelles. Two steps of the purification were adjusted to improve detergent exchange and to reduce the amount of $^2$H-LMNG used, respectively. During the first PC resin purification step, the column was washed 6 instead of 4 times with 3 CVs of PCWB. The dialysis step to remove imidazole was carried out with 250 ml PCB instead of 500 ml. This exchange protocol was used for the first purification batch ($\alpha_{1B}$-AR-B1D1 with prazosin, tamsulosin, and ρ-TIA for dynamics experiments at 320 K) and yielded an exchange of $^1$H-LMNG to $^2$H-LMNG on the order of 50%, based on the detergent signals in the [$^{13}$C,$^1$H]-HSQC spectrum. The exchange was further improved by introducing additional exchange steps while the receptor was bound on the PC resin. After the overnight binding of the receptor to the PC resin, the column was washed 4 times with 3 CVs of PCWB (containing protonated LMNG) and then once with exchange buffer (EXB; 20 mM Na-phosphate pH 7 at RT, 20 mM NaCl, 0.05% $^2$H-LMNG). The detergent was exchanged during 6 incubation periods with 5 ml EXB per mg of $\alpha_{1B}$-AR-B1D1. The PC resin was rolled for 2 h during each incubation step, except for the last one, which was done overnight. The next day, the PC resin was washed 2 times with 3 CVs of PCB before continuing with the established imidazole (PCIEB) protocol. The imidazole was removed using two PD-10 desalting column steps instead of using just one PD-10 desalting step that is followed by a dialysis step. This modified purification protocol was used to produce the second purification batch ($\alpha_{1B}$-AR-B1D1 with prazosin and in the apo form for dynamics experiments at 298 K) and yielded an exchange of $^1$H-LMNG to $^2$H-LMNG of 95%, based on integration of the residual detergent signals in the [$^{13}$C,$^1$H]-HSQC spectrum.

#### 1.4 TRACT

Equations and parameters used to calculate $\tau_c$ values based on Robson et al. (Robson et al. 2021) are shown below. The assumed $S^2_{axis}$ for backbone amides of 0.9 improves the accuracy of the correlation times since also amides within rigid regions of proteins possess order parameters that are smaller than one (0.85 – 0.95). The $\tau_c$ values obtained from individual amides of prazosin- and ρ-TIA-bound $\alpha_{1B}$-AR-B1D1 are listed in the Supplementary Data 2 in Table S12 and S13. The deposited tables include only values that passed all quality filters and were used to calculate global correlation times. The tables include the values for $\tau_c$ (ns) and their associated standard errors (SE) as estimated using 1 000 Monte Carlo samplings based on the SEs of the α- and β-rates. The values and SEs of the α- and β-rates are given in the tables as well. Some considerations regarding the calculation of $\tau_c$ are discussed in more detail in Section 3.

$$\tau_c = \frac{\sqrt[3]{125\, c^3\omega_N^6 + 24\sqrt{3}\sqrt{625\,(S^2_{axis})^2\, c^4\omega_N^{10} - 3025\,(S^2_{axis})^4\, c^2\omega_N^8 + 21952\,(S^2_{axis})^6\,\omega_N^6} + 1800\,(S^2_{axis})^2\, c\,\omega_N^4}}{24\,S^2_{axis}\,\omega_N^2} - \frac{336\,(S^2_{axis})^2\,\omega_N^2 - 25\,c^2\omega_N^4}{24\,S^2_{axis}\,\omega_N^2\sqrt[3]{125\, c^3\omega_N^6 + 24\sqrt{3}\sqrt{625\,(S^2_{axis})^2\, c^4\omega_N^{10} - 3025\,(S^2_{axis})^4\, c^2\omega_N^8 + 21952\,(S^2_{axis})^6\,\omega_N^6} + 1800\,(S^2_{axis})^2\, c\,\omega_N^4}} + \frac{5\,c}{24\,S^2_{axis}}$$

$$c = \frac{R_\beta - R_\alpha}{2p\delta_N(3\,cos^2\theta - 1)}$$

$$p = \frac{\mu_0\,\gamma_H\,\gamma_N\,h}{16\,\pi^2\,\sqrt{2}\,r_{HN}^3}$$

$$\delta_N = \frac{\gamma_N\,B_0\,\Delta\delta_N}{3\sqrt{2}}$$

$\mu_0 = 4\,\pi \times 10^{-7}$ H/m
$h = 6.62607004 \times 10^{-34}$ J Hz$^{-1}$
$\theta = 17°$
$\gamma_H = 267.52218744 \times 10^6$ rad s$^{-1}$ T$^{-1}$
$\gamma_N = -27.116 \times 10^6$ rad s$^{-1}$ T$^{-1}$
$r_{HN} = 1.02 \times 10^{-10}$ m
$\Delta\delta_N = 160 \times 10^{-6}$
$B_0 = 16.4438966$ T
$\omega_N = -\gamma_N B_0$
$S^2_{axis} = 0.9$

#### 2. Spectra and Assignments

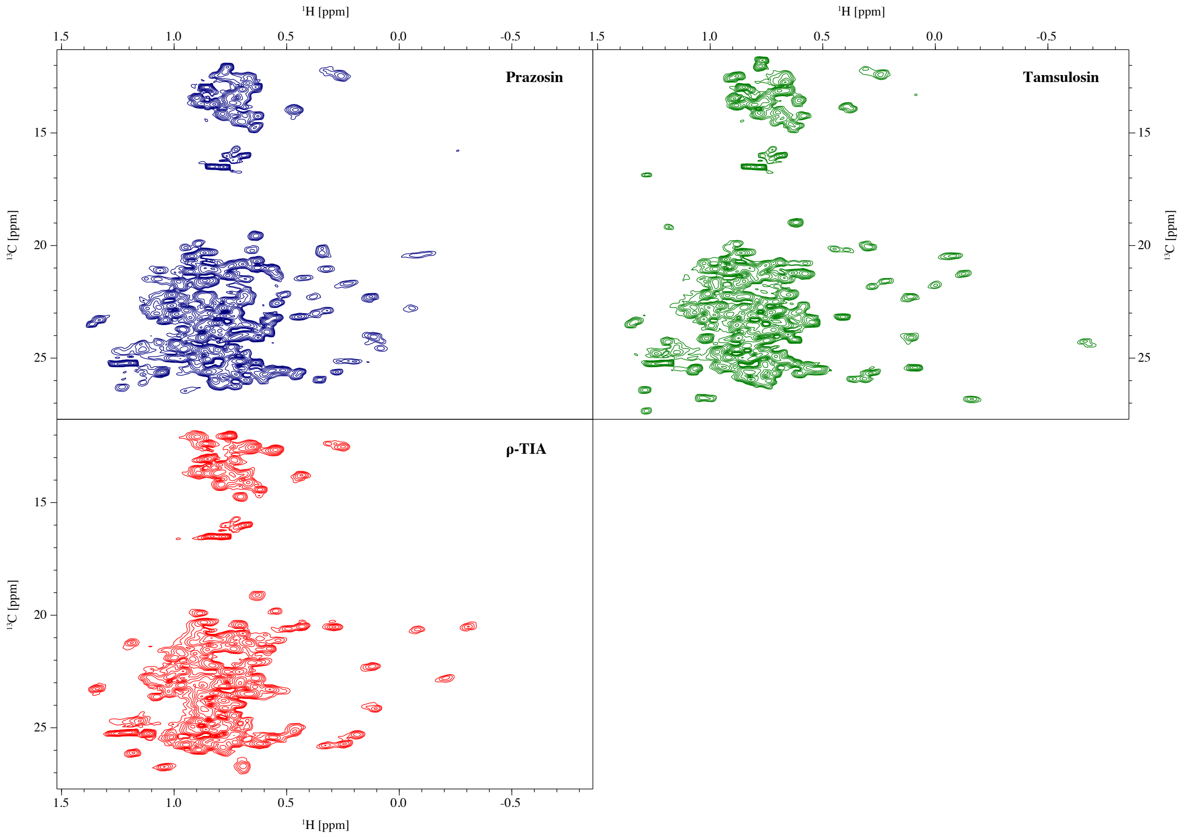


**Figure S2.1.** [$^{13}$C,$^{1}$H]-HSQC spectra of α$_{1B}$-AR-B1D1 binding to each of the three inverse agonists at 320 K with labeled δ-methyl groups in Ile and Leu side chains and labeled γ-methyl groups in Val side chains. Spectra were recorded at 700 MHz.

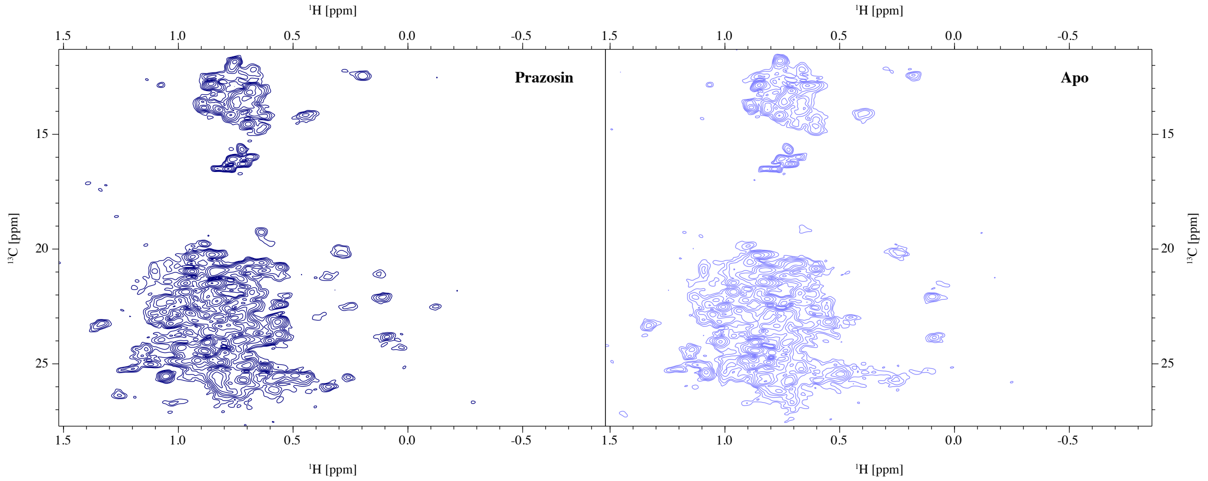


**Figure S2.2.** [$^{13}$C,$^{1}$H]-HSQC spectra of α$_{1B}$-AR-B1D1 binding prazosin and in the apo state at 298 K with labeled δ-methyl groups in Ile and Leu side chains and labeled γ-methyl groups in Val side chains. Spectra were recorded at 700 MHz.

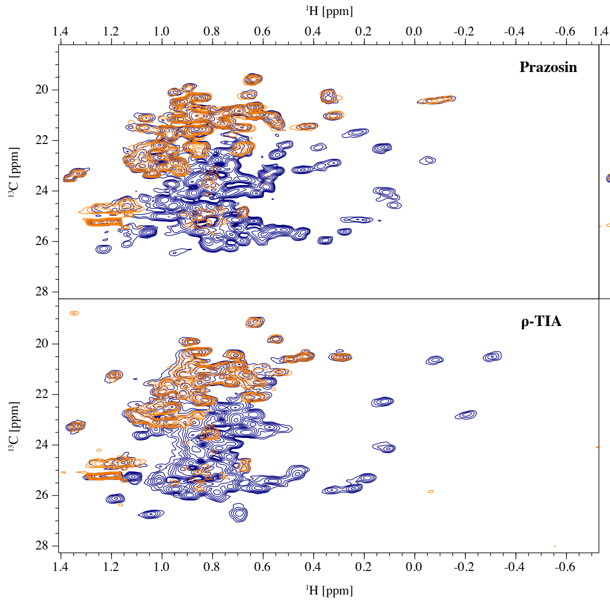


**Figure S2.3.** [$^{13}$C,$^{1}$H]-HSQC spectra of α1B-AR-B1D1 binding to each of the three inverse agonists at 320 K using different labeling schemes to distinguish between Leu and Val methyl groups. Spectra in blue show labeled Leu and Val methyl groups by supplementation of α-ketoisovaleric acid to the expression. Spectra in orange are based on selective labeling of Val methyl groups based on additional supplementation of unlabeled Leu to the expression with α-ketoisovaleric acid (Mas et al. 2013).

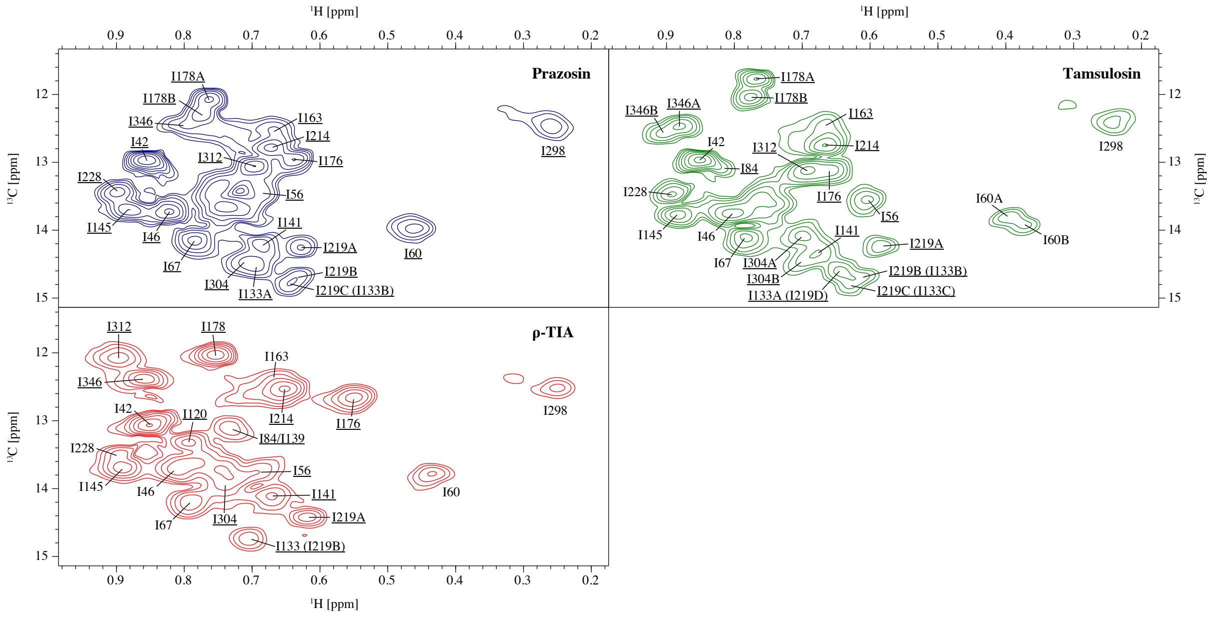


**Figure S2.4.** Assigned α1B-AR-B1D1 Ile δ-methyl groups with the three inverse agonists at 320 K. Assignments that were confirmed by point mutation to either Leu or Val are underlined. Non-underlined assignments were inferred from the assignment in presence of prazosin and were not further confirmed using the corresponding point mutation. Ambiguities are indicated by stating all possible assignments, with less likely assignments in parentheses. This mostly concerns the assignments for I133 and I219 that show considerable shift changes when the other residue is mutated due to their spatial proximity. Multiple peaks for the same residue are labeled alphabetically.

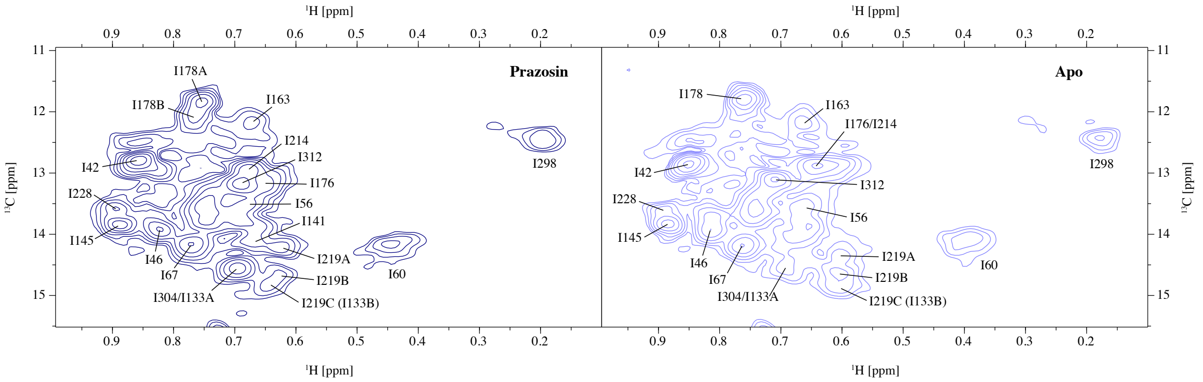


**Figure S2.5.** Assigned α1B-AR-B1D1 Ile δ-methyl groups at 298 K. Assignments were adapted from the assignment of prazosin-bound receptor at 320 K via an additional spectrum at 310 K.

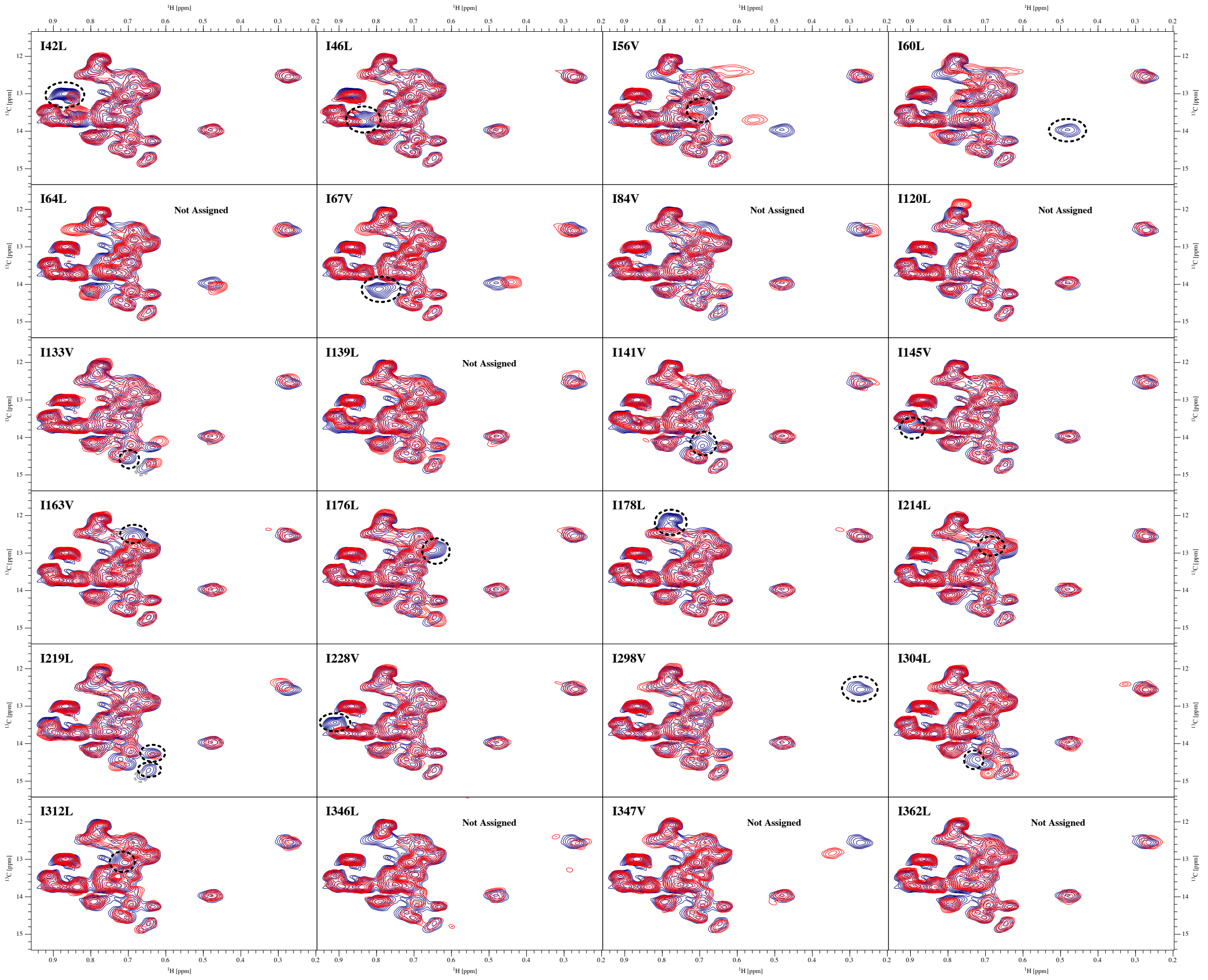


**Figure S2.6.** [$^{13}$C,$^{1}$H]-HSQC spectra of all α1B-AR-B1D1 Ile to Leu or Val mutations with prazosin. Spectra were recorded at 320 K and at 700 MHz. Assignments are highlighted by dashed circles. The ambiguous assignment for the peak that belongs to either I133 or I219 is highlighted in grey. Correlations in the 3D $^{13}$C-resolved [$^{1}$H,$^{1}$H]-NOESY confirmed that the two peaks assigned to I219 belong to the same residue (Figure S2.7).

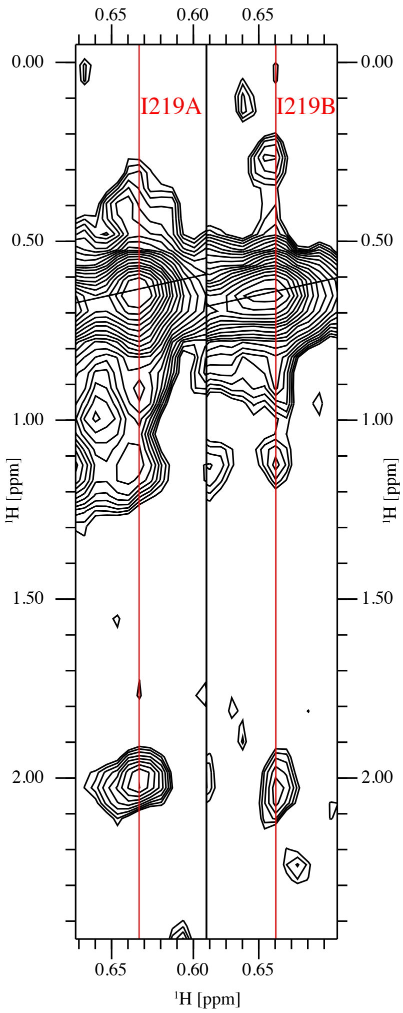


**Figure S2.7.** Strips from the 3D $^{13}$C-resolved [$^{1}$H,$^{1}$H]-NOESY of ILV labeled, prazosin-bound $\alpha_{1B}$-AR-B1D1 recorded at 320 K and at 700 MHz. The strips belong to the two peaks from I219, with I219A being the one with the smaller carbon chemical shift (16.89 ppm vs 17.41 ppm for I219B). The cross peak at 2 ppm was not present for other Ile residues, indicating that both peaks stem from the same residue and that the residual intensity at the position of I219A with the I219L-mutant stems from another Ile methyl group.

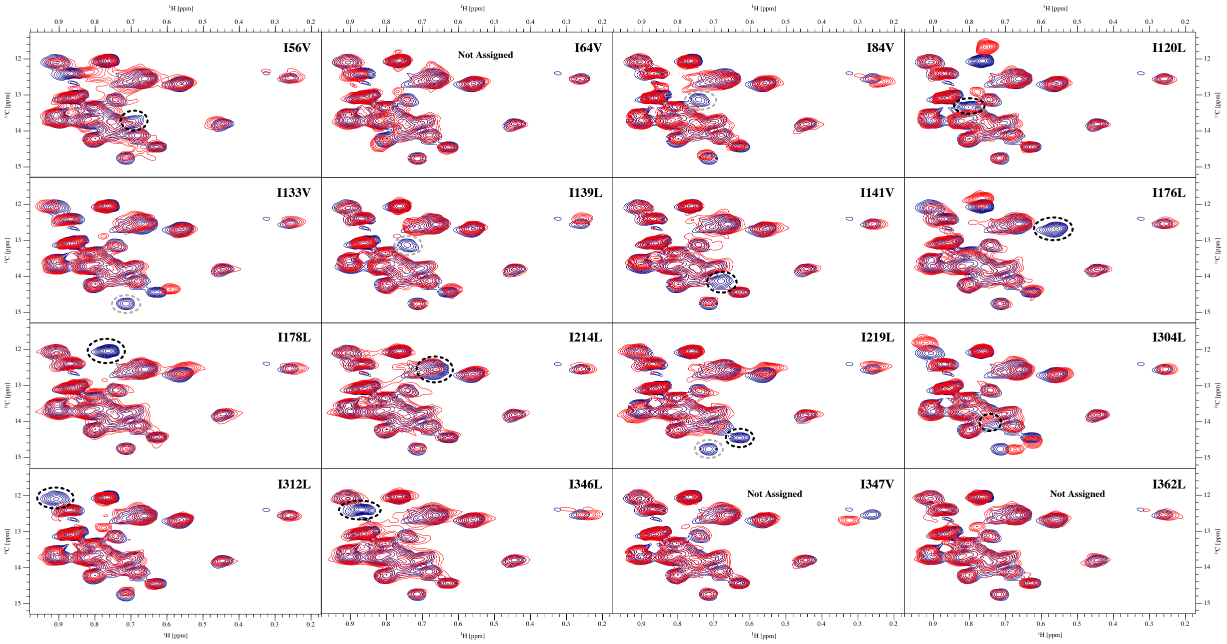


**Figure S2.8.** [$^{13}$C,$^{1}$H]-HSQC spectra of all measured α$_{1B}$-AR-B1D1 Ile to Leu or Val mutations with ρ-TIA. Spectra were recorded at 320 K and at 700 MHz. Assignments are highlighted by dashed circles with ambiguous ones highlighted in grey.

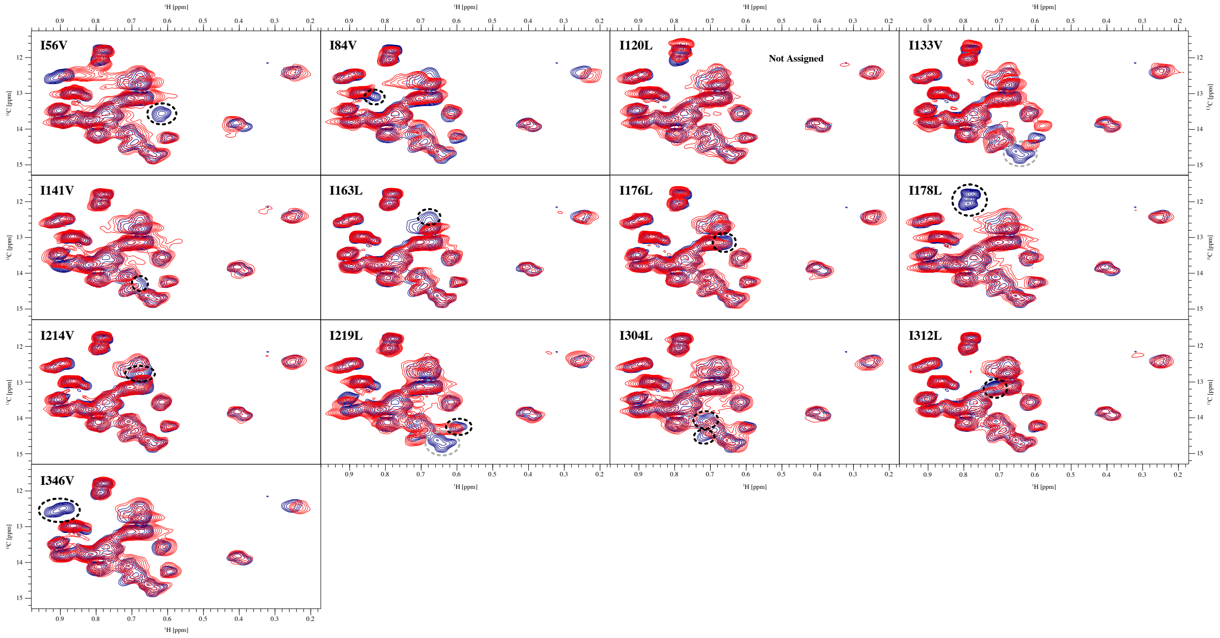


**Figure S2.9.** [$^{13}$C,$^{1}$H]-HSQC spectra of all measured α$_{1B}$-AR-B1D1 Ile to Leu or Val mutations with tamsulosin. Spectra were recorded at 320 K and at 700 MHz. Assignments are highlighted by dashed circles with ambiguous ones highlighted in grey.

### 3. Correlation Time Estimation using TRACT

Rotational correlation times were determined for the prazosin- and ρ-TIA-bound $\alpha_{1B}$-AR-B1D1 at 320 K using a 2D version of the TRACT experiment. Using a 2D TRACT allows the removal of side-chain amides and flexible backbone amides from the calculation, thereby increasing the accuracy of the $\tau_c$ estimation. A total of 113 and 88 backbone amides gave reasonable $\tau_c$ values (SE below 12 ns, $\tau_c$ values < 20 ns) for the measurement with prazosin and ρ-TIA, respectively. The values were distributed around mean values, which were regarded as the actual rotational correlation times of $\alpha_{1B}$-AR-B1D1 binding prazosin (37.07 ns) and ρ-TIA (36.49 ns) (Figure S3.1). The means did not differ significantly in a Welch's t-test (p-value: 0.574). We also attempted to measure the correlation time of prazosin-bound $\alpha_{1B}$-AR-B1D1 at 298 K using two different setups that differed in the delays and in the number of scans used. With both setups, $\tau_c$ values of up to roughly 120 ns were obtained. They appeared almost uniformly distributed and it was therefore not possible to obtain any reliable value for the correlation time of the receptor. Hence, only the correlation times that were obtained at 320 K were used to calculate order parameters and are discussed in this Section.

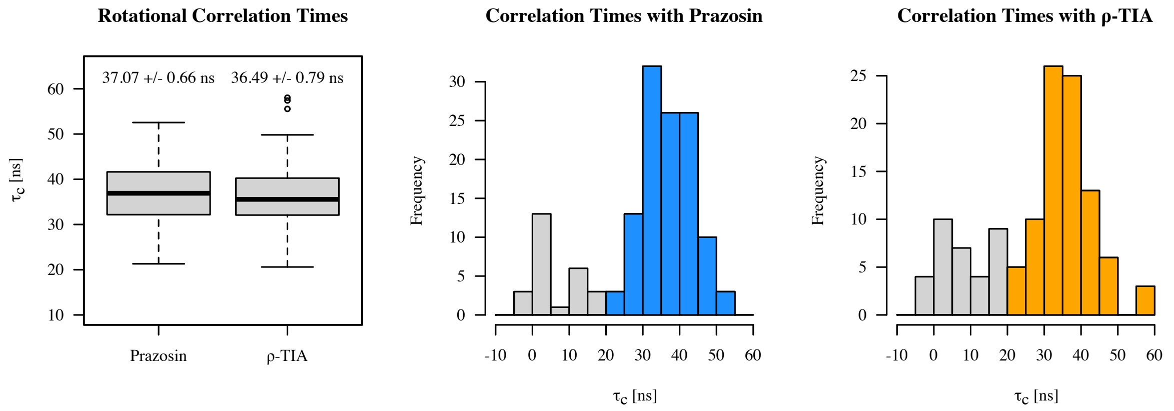


**Figure S3.1.** Rotational correlation times for $\alpha_{1B}$-AR-B1D1 binding prazosin and ρ-TIA at 320 K. Apparent $\tau_c$ values were determined from every amide peak that was visible in the spectra. The boxplot on the left shows the distribution of backbone amides with apparent $\tau_c$ values > 20 ns, which have standard errors smaller than 12 ns. The values at the top in the boxplot indicate the means and their standard errors. The difference in the mean between the ligands is not significant (t-test p-value: 0.574). The histogram on the left shows the apparent $\tau_c$ values as measured with the $\alpha_{1B}$-AR-B1D1 prazosin complex and the one on the right the values as measured with ρ-TIA. All apparent $\tau_c$ values with standard errors smaller than 12 ns are shown. Apparent $\tau_c$ values larger than 20 ns were used to calculate the global correlation time and are highlighted in blue and orange.

The use of a 2D version of the TRACT experiment enabled $\tau_c$ values for individual amides to be calculated, which appeared to be distributed around a mean value for the $\alpha_{1B}$-AR-B1D1 bound to either prazosin or ρ-TIA (Figure S3.1). There might be only two main reasons for amide $\tau_c$ values to be different from actual $\tau_c$ values that report on the rotational correlation time of a protein. For conciseness, the values of individual amide groups are referred to as apparent $\tau_c$ values and the value for the actual rotational correlation time of a protein as global $\tau_c$ value. The first possible cause for the distribution is the presence of residual dynamics. Movements of the backbone on a similar timescale as the rotational correlation time of the protein might lead to apparent $\tau_c$ values of the backbone that are smaller than the global $\tau_c$ value. Local differences in these residual backbone dynamics might cause a distribution of apparent $\tau_c$ values as could have been observed in this case. If this were the underlying cause of the distribution, then, using the mean apparent $\tau_c$ value as global correlation time would lead to an underestimation of the correlation time. Because $\alpha_{1B}$-AR-B1D1 is solubilized directly from the *E. coli* membrane without any refolding after expression in $D_2O$, only the water-accessible amide groups can be observed by NMR. This includes all the loops and ends of helices, whereas large fractions of the transmembrane regions remain deuterated and hence invisible for NMR. Especially loop regions might show additional backbone flexibility and thus contain amides with apparent $\tau_c$ values that are smaller than the global $\tau_c$ value. If this were the case, then the largest apparent $\tau_c$ values would be the ones closest to the global $\tau_c$ value and they should be occurring at some elevated frequency as they would represent the rigid core of the protein. A distribution as with the $\alpha_{1B}$-AR-B1D1, in which the largest apparent $\tau_c$ values appear only in the tail of the distribution, would imply that the entire protein experiences residual dynamics and thus no apparent $\tau_c$ value would necessarily correspond to the global $\tau_c$. Hence, none or almost none of obtained ~100 apparent $\tau_c$ values would stem from rigid regions within $\alpha_{1B}$-AR-B1D1, which appears to be unlikely. However, residual dynamics appear to be the most likely explanation for very small apparent $\tau_c$ values below 20 ns. Further, local variations in order parameters ($S^2_{axis}$) of rigid amides as mentioned by Robson et al. (Robson et al. 2021) are insufficient to explain the observed variations since they would only lead to apparent $\tau_c$ values between 35.00 ns ($S^2_{axis}$ = 0.85) and 39.14 ns ($S^2_{axis}$ = 0.95) if the global $\tau_c$ is 37.07 ns.

The most likely source for the observed variations in apparent $\tau_c$ values stems from dispersion in magnitude ($\Delta\delta_N$) and orientation ($\theta$) of the $^{15}N$ chemical shift anisotropy (CSA) tensors. Both of these values are required to calculate rotational correlation times but are generally not determined. The calculation of the apparent $\tau_c$ values for each amide is thus based on the average values for these two parameters ($\Delta\delta_N$ = 160 ppm; $\theta$ = 17°) (Lee et al. 2006; Robson et al. 2021). However, these parameters are known to vary within a range of 125 to 216 ppm for $\Delta\delta_N$ and of 6 to 26° for $\theta$ (Saito et al. 2010). Especially variations in $\Delta\delta_N$ might lead to errors of up to 15% for $\tau_c$ values (Robson et al. 2021). In the classical 1D TRACT experiment, these variations average out because only the ensemble of all amides is considered for the $\tau_c$ calculation, which is not the case for 2D TRACT data. In this case, the apparent $\tau_c$ values contain contributions from local CSA. To test whether such CSA dispersion could explain the observed distributions of apparent $\tau_c$ values, we used simulations based on the experimental global $\tau_c$ values and normally distributed $\Delta\delta_N$ and $\theta$ values (Figure S3.2). The resulting distributions of simulated apparent $\tau_c$ values resemble the distributions of the experimental apparent $\tau_c$ values. Simulated values range from roughly 20 to 55 ns with average values slightly smaller than expected. Only very few simulated apparent $\tau_c$ values were smaller than 20 ns, suggesting that experimentally obtained values in that range more likely reflect residual dynamics than uncommon $\Delta\delta_N$ and $\theta$ values. Thus, apparent $\tau_c$ values below this threshold were attributed to residual dynamics. Apparent $\tau_c$ values below 20 ns appear to be more frequent for the $\alpha_{1B}$-AR-B1D1 ρ-TIA than the prazosin complex, indicating the presence of increased backbone dynamics when the allosteric peptide is bound. All in all, the simulations suggest that the observed distributions of experimental apparent $\tau_c$ values are likely based on the variation in CSA.

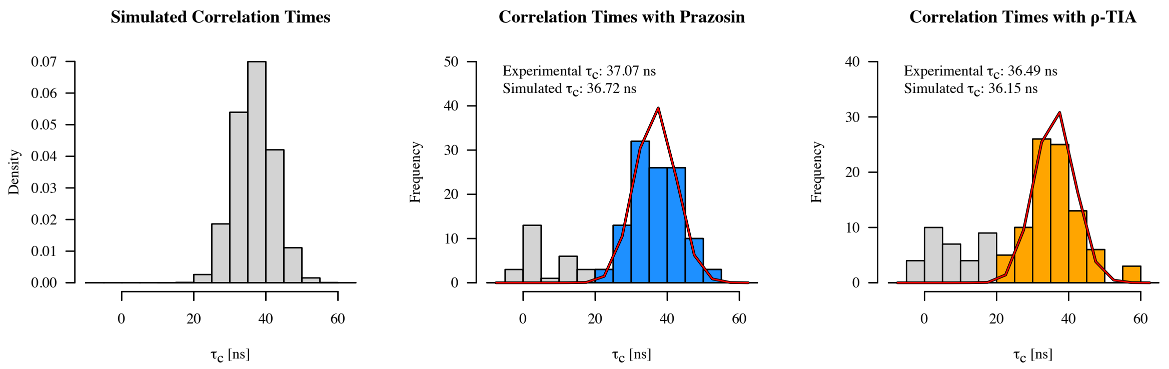


**Figure S3.2.** Histograms of simulated (left) and experimental (middle & right) apparent $\tau_c$ values. The left histogram shows a simulated distribution of apparent $\tau_c$ values based on 10 000 samplings of normally distributed $\Delta\delta_N$ and $\theta$ values with a global $\tau_c$ of 37.07 ns, corresponding to the one that was measured for prazosin-bound $\alpha_{1B}$-AR-B1D1. The distribution of these parameters does not necessarily follow a Gaussian one, but this was assumed for simplicity. The mean CSA values ($\Delta\delta_N$ = 160 ppm; $\theta$ = 17°) are based on Robson et al. (Robson et al. 2021) and their standard deviations ($\Delta\delta_N$ = 19 ppm; $\theta$ = 5°) on Saitô et al. (Saito et al. 2010). The experimental $\tau_c$ values as obtained for prazosin- and ρ-TIA-bound $\alpha_{1B}$-AR-B1D1 are depicted in the middle and on the right, respectively. Values used to calculate the global rotational correlation time are indicated in blue and orange. Values in grey highlight apparent $\tau_c$ values that were excluded from the estimation because they likely stem from amides that undergo residual dynamics. The curves in red show simulated apparent $\tau_c$ values based on the experimental mean values for prazosin- and ρ-TIA-bound $\alpha_{1B}$-AR-B1D1. The simulated global $\tau_c$ is given together with the experimental one.

The main problem caused by the variations in CSA is that it is not possible to distinguish amides that show low apparent $\tau_c$ values because of locally increased dynamics from amides that have uncommon CSA values, resulting in either under- or overestimates of the global $\tau_c$. For the calculation of the rotational correlation time of $\alpha_{1B}$-AR-B1D1, we assumed that the variation in CSA causes the distribution of apparent $\tau_c$ values around the global $\tau_c$ value of the receptor, and that residual backbone dynamics can be neglected for these apparent $\tau_c$ values. We further assumed that apparent $\tau_c$ values below 20 ns belong to residues with residual dynamics, and we thus excluded them from the calculation of the global $\tau_c$. We note that the individual removal of such amides is a clear advantage of the 2D experiment over the 1D version, for which $\tau_c$ values represent a lower limit for the correlation time due to the contribution of amides within flexible regions. However, it is not necessarily clear whether a particular apparent $\tau_c$ value is small due to residual dynamics or because of uncommon $\Delta\delta_N$ and $\theta$ values. The cutoff of 20 ns for apparent $\tau_c$ values seems to agree with the simulations, but the estimation of the global $\tau_c$ values of course depends on where exactly this cutoff is set. If values down to 10 ns would be included, then the global $\tau_c$ values would change from 37.07 to 35.38 ns for prazosin-bound and from 36.49 to 34.50 ns for ρ-TIA-bound $\alpha_{1B}$-AR-B1D1.

### 4. Side-Chain Dynamics

##### 4.1 $\alpha_{1B}$-AR-B1D1 Methyl Dynamics Data

The ratio of the intensities obtained from the forbidden and the allowed experiments often gave reasonable curves that allowed to fit $\eta$ and $\delta$ values reliably. Multiple cut-offs were used to determine the quality of the fits and to remove unreliable values. More generous cut-offs were chosen in order to not introduce any undesired biases, especially towards more dynamic side chains. Since NMR properties become more unfavorable in rigid side chains, the signal-to-noise ratio decreases for rigid side chains, leading to worse fits. Therefore, the use of too strict cut-offs would eliminate many of those rigid methyl groups such that the protein would appear more dynamic than it is. The decreasing quality of the NMR properties are also displayed by the correlation between $\delta$ values, a measure for other relaxation sources, and the side-chain dynamics, which indicates that rigid methyl groups are more prone to undergo additional relaxation. The decreased quality of the fits for rigid side chains led to larger standard errors for larger order parameters. To generally improve the quality of the fits, an automated removal of outliers was implemented into the fitting procedure. If the removal of one or two data points improved the fit beyond a defined threshold, then the corresponding data points were not considered for the final fit. Thresholds were set to agree with a visual identification of outliers. Outliers occurred generally at longer delays, where ratios were affected more by noise because the intensities decreased with increasing delays in the spectra of both experiments. The number of data points that were removed to obtain the presented side-chain dynamics are indicated by the degree of freedom (df) of the remaining data points in the tables in the Supplementary Data 2. This outlier selection might risk overfitting in some cases. In cases where outliers were removed $\eta$ values did not necessarily change much. However, the associated standard errors could decrease considerably, leading to order parameters that might appear to be better defined than they actually are.

#### 4.2 Comparisons Between $S^2_{axis}$ Distributions

To test whether the inverse agonists altered the side-chain dynamics of the receptor, the fractions of order parameters within each motional class were compared between the data sets recorded with the different ligands (Figure S4.2.1). Some differences between the ligands are present; however, they are of a similar size as the ones detected between the two data sets with prazosin-bound $\alpha_{1B}$-AR-B1D1, which were recorded on the same sample. Thus, the differences are best explained by experimental variability.

It is likely that we obtained order parameters of a somewhat different set of residues in each data set. Therefore, we need to consider sampling bias for the comparison of the fractions. We used bootstrapping to test the extent to which the $S^2_{axis}$ fractions vary within motional classes depending on which data are used exactly. Bootstrapping randomly resamples the data. For each of the resampling, the fraction of $S^2_{axis}$ within each motional class was determined. This gives a range within which the fractions vary depending on which values are included. This approach was used to mimic the sampling bias. Overall, bootstrapping indicates that the differences between the ligands are not due to changes in side-chain dynamics but rather due to the experimental variability.

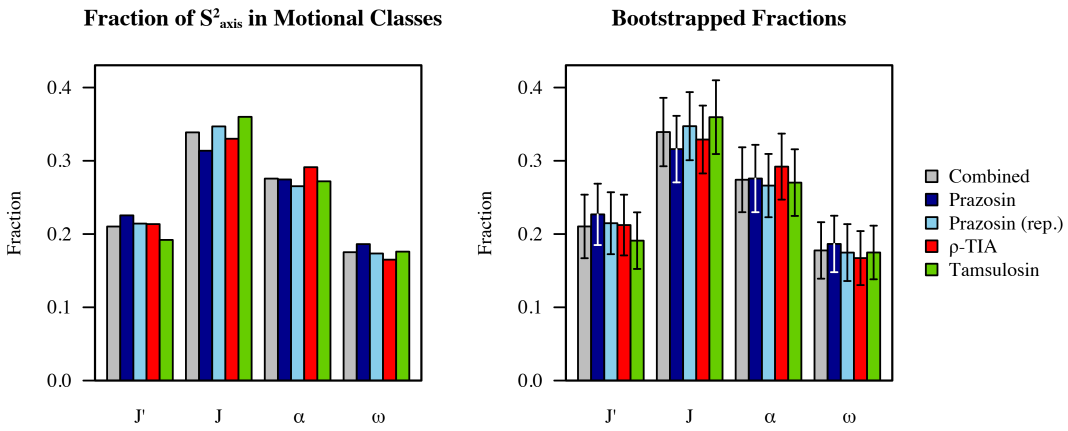


**Figure S4.2.1.** Fraction of order parameters found within the given motional classes based on the obtained data directly (left) or based on 1 000 bootstrap repeats using sample sizes of 100 (right). Error bars indicate the standard deviation of the bootstrapped fractions.

#### 4.3 Side-Chain Dynamics at 298 K

A comparison of the relative side-chain dynamics between apo and prazosin-bound $\alpha_{1B}$-AR-B1D1 shows an overall decrease in η values for the apo receptor (Figure S4.3.1). The observed difference could indicate that the apo receptor has either generally increased side-chain dynamics or a shorter $\tau_c$ compared to the prazosin-bound form. Since it was not possible to obtain reliable $\tau_c$ values for the correlation times at 298 K, it is not possible to distinguish between these two possible causes. We hypothesized that if both η data sets, when scaled to show identical means, still reveal differences in how individual motional classes were populated, then that would indicate true differences in side-chain dynamics. If, however, the motional classes become similarly populated in the scaled data sets, then the differences are rather due to a change in $\tau_c$.

If order parameters are assumed to be temperature-independent, then it is possible to estimate the correlation time at 298 K by comparing the η values with the $S^2_{axis}$ values obtained at 320 K (Figure S4.3.1). Linear regression with y-intercept set to zero was used to correlate the Ile dynamics of prazosin-bound receptors between both temperatures. A $\tau_c$ of 77.8 ns was obtained for prazosin-bound $\alpha_{1B}$-AR-B1D1 at 298 K, resulting in an $S^2_{axis}$ mean of 0.53, which is close to 0.51 as was obtained with all three ligands at 320 K. The correlation time of the apo receptor was then scaled to yield an identical mean $S^2_{axis}$ value. The resulting $\tau_c$ of the apo receptor was 70.8 ns, and therefore 7.0 ns smaller than the one estimated for the prazosin-bound receptor. With this $S^2_{axis}$ scaling of $\tau_c$, differences in dynamics between liganded and apo $\alpha_{1B}$-AR-B1D1 essentially disappear when amino acid types are compared (Figure S4.3.1).

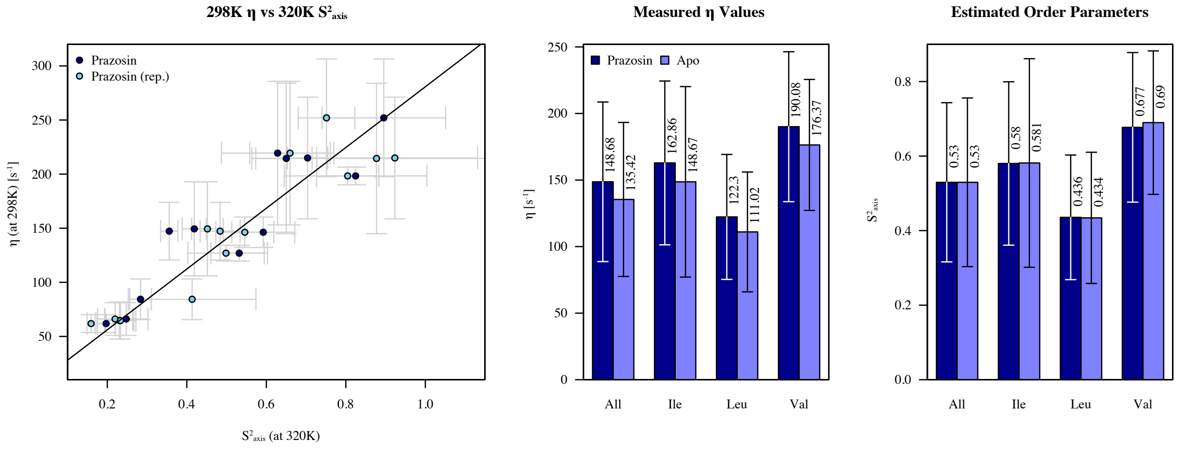


**Figure S4.3.1.** The rotational correlation time $\tau_c$ at 298 K of prazosin-bound $\alpha_{1B}$-AR-B1D1 was estimated by correlating 298 K Ile η values with $S^2_{axis}$ values that were obtained at 320 K (left). Error bars indicate the standard errors. Bar graphs compare the mean side-chain dynamics between η values (middle) and estimated order parameters (right). Estimated order parameters are based on different $\tau_c$ values for the prazosin-bound and the apo $\alpha_{1B}$-AR-B1D1.

A comparison of the number of order parameters per motional class indicates that only minor receptor-wide differences between the apo and the liganded receptor might be present, which appear negligible once the populations are resampled using bootstrapping (Figure S4.3.2). Overall, the fractions of $S^2_{axis}$ within the various motional classes appear to fluctuate within a similar range as with the different inverse agonists at 320 K (Figure S4.2.1). Based on these findings, we attribute the differences in $\eta$ values to a difference in $\tau_c$ between apo and liganded receptor. The difference in $\tau_c$ may suggest differences in the hydrodynamic radius of liganded and apo $\alpha_{1B}$-AR-B1D1, or the presence of additional dynamics on timescales similar to $\tau_c$ of the apo receptor. Additionally, it may indicate that the correlation time is highly sensitive to sample conditions, e.g. to the amount of free micelles or the viscosity of the sample.

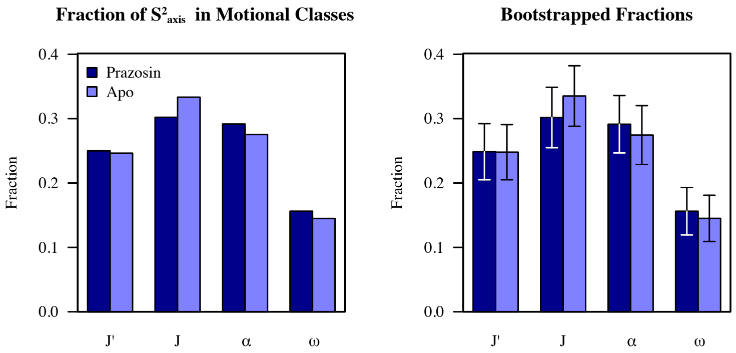


**Figure S4.3.2.** Fractions of order parameters found within given motional classes based on the estimated order parameters (left) and with 1 000 bootstrap repeats using a sample size of 100 (right). Error bars indicate the standard deviation of the bootstrapped fractions. Motional classes were determined using k-means clustering on the estimated order paramters at 298 K.

#### 4.4 Ligand-Dependent Differences in Ile $S^2_{axis}$

The significances of the differences between Ile order parameters of the differently liganded states were determined by a Monte Carlo approach. Order parameters were resampled 5 million times based on Student's t-distributions centered around the experimental value and by using their standard errors as standard deviations (Figure S4.4.1 & S4.4.2). The frequency by which the absolute difference between sampled order parameters were larger than the absolute experimental difference was taken as the p-value. This corresponds to a two-tailed test. We generally used a cut-off of 5% as indicator of significance but considered p-values up to 10% as being potentially interesting. The standard error of each $S^2_{axis}$ contains the uncertainty from the $\tau_c$ determination, which has no impact on the comparison between $S^2_{axis}$ values. This stems from the fact that an error in $\tau_c$ would affect all $S^2_{axis}$ values in the same way. Therefore, a small but systematic overestimation of the $S^2_{axis}$ error is present when significances for differences were calculated. Further, we acknowledge that different approaches could be used to determine significances between order parameters, and that these different methods might lead to different assessments on which differences are significant.

I178$^{4x58}$ gave rise to two signals in the [$^{13}$C,$^{1}$H]-HSQC spectra for the two tested orthosteric ligands, indicating the presence of two different conformational states for I178$^{4x58}$ itself or in its proximity. The second conformation was absent or not detectable with ρ-TIA-bound $\alpha_{1B}$-AR-B1D1. The side-chain dynamics of I178$^{4x58}$ appeared to be identical across the different ligands and the two different conformations (Figure S4.4.3). The $S^2_{axis}$ of I178A$^{4x58}$ in tamsulosin-bound $\alpha_{1B}$-AR-B1D1 is not significantly smaller than any other I178$^{4x58}$ $S^2_{axis}$ value.

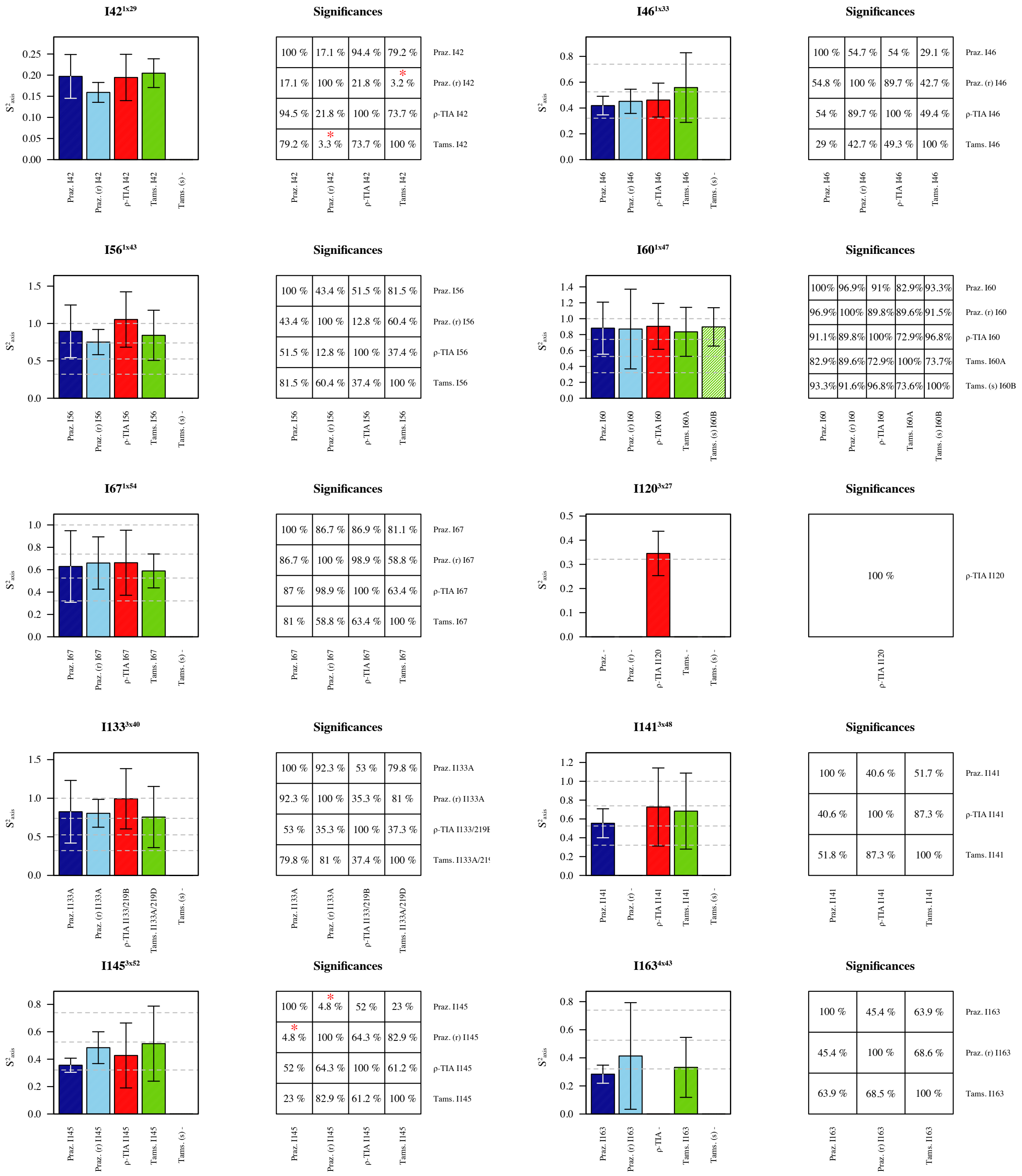


**Significances – I42**

| | Praz. I42 | Praz. (r) I42 | ρ-TIA I42 | Tams. I42 |
|---|---|---|---|---|
| Praz. I42 | 100 % | 17.1 % | 94.4 % | 79.2 % |
| Praz. (r) I42 | 17.1 % | 100 % | 21.8 % | * 3.2 % |
| ρ-TIA I42 | 94.5 % | 21.8 % | 100 % | 73.7 % |
| Tams. I42 | 79.2 % | * 3.3 % | 73.7 % | 100 % |

**Significances – I46**

| | Praz. I46 | Praz. (r) I46 | ρ-TIA I46 | Tams. I46 |
|---|---|---|---|---|
| Praz. I46 | 100 % | 54.7 % | 54 % | 29.1 % |
| Praz. (r) I46 | 54.8 % | 100 % | 89.7 % | 42.7 % |
| ρ-TIA I46 | 54 % | 89.7 % | 100 % | 49.4 % |
| Tams. I46 | 29 % | 42.7 % | 49.3 % | 100 % |

**Significances – I56**

| | Praz. I56 | Praz. (r) I56 | ρ-TIA I56 | Tams. I56 |
|---|---|---|---|---|
| Praz. I56 | 100 % | 43.4 % | 51.5 % | 81.5 % |
| Praz. (r) I56 | 43.4 % | 100 % | 12.8 % | 60.4 % |
| ρ-TIA I56 | 51.5 % | 12.8 % | 100 % | 37.4 % |
| Tams. I56 | 81.5 % | 60.4 % | 37.4 % | 100 % |

**Significances – I60**

| | Praz. I60 | Praz. (r) I60 | ρ-TIA I60 | Tams. I60A | Tams. (s) I60B |
|---|---|---|---|---|---|
| Praz. I60 | 100% | 96.9% | 91% | 82.9% | 93.3% |
| Praz. (r) I60 | 96.9% | 100% | 89.8% | 89.6% | 91.5% |
| ρ-TIA I60 | 91.1% | 89.8% | 100% | 72.9% | 96.8% |
| Tams. I60A | 82.9% | 89.6% | 72.9% | 100% | 73.7% |
| Tams. (s) I60B | 93.3% | 91.6% | 96.8% | 73.6% | 100% |

**Significances – I67**

| | Praz. I67 | Praz. (r) I67 | ρ-TIA I67 | Tams. I67 |
|---|---|---|---|---|
| Praz. I67 | 100 % | 86.7 % | 86.9 % | 81.1 % |
| Praz. (r) I67 | 86.7 % | 100 % | 98.9 % | 58.8 % |
| ρ-TIA I67 | 87 % | 98.9 % | 100 % | 63.4 % |
| Tams. I67 | 81 % | 58.8 % | 63.4 % | 100 % |

**Significances – I120**

| | ρ-TIA I120 |
|---|---|
| ρ-TIA I120 | 100 % |

**Significances – I133**

| | Praz. I133A | Praz. (r) I133A | ρ-TIA I133/219B | Tams. I133A/219D |
|---|---|---|---|---|
| Praz. I133A | 100 % | 92.3 % | 53 % | 79.8 % |
| Praz. (r) I133A | 92.3 % | 100 % | 35.3 % | 81 % |
| ρ-TIA I133/219I | 53 % | 35.3 % | 100 % | 37.3 % |
| Tams. I133A/21( | 79.8 % | 81 % | 37.4 % | 100 % |

**Significances – I141**

| | Praz. I141 | ρ-TIA I141 | Tams. I141 |
|---|---|---|---|
| Praz. I141 | 100 % | 40.6 % | 51.7 % |
| ρ-TIA I141 | 40.6 % | 100 % | 87.3 % |
| Tams. I141 | 51.8 % | 87.3 % | 100 % |

**Significances – I145**

| | Praz. I145 | Praz. (r) I145 | ρ-TIA I145 | Tams. I145 |
|---|---|---|---|---|
| Praz. I145 | 100 % | * 4.8 % | 52 % | 23 % |
| Praz. (r) I145 | * 4.8 % | 100 % | 64.3 % | 82.9 % |
| ρ-TIA I145 | 52 % | 64.3 % | 100 % | 61.2 % |
| Tams. I145 | 23 % | 82.9 % | 61.2 % | 100 % |

**Significances – I163**

| | Praz. I163 | Praz. (r) I163 | Tams. I163 |
|---|---|---|---|
| Praz. I163 | 100 % | 45.4 % | 63.9 % |
| Praz. (r) I163 | 45.4 % | 100 % | 68.6 % |
| Tams. I163 | 63.9 % | 68.5 % | 100 % |

**Figure S4.4.1.** Comparison of Ile δ-methyl order parameters obtained with inverse agonists at 320 K. $S^2_{axis}$ values obtained from additional signals occurring only with the tamsulosin-bound $\alpha_{1B}$-AR-B1D1 are labeled with (s) and highlighted as hatched bars. Assignments correspond to the ones shown in Figure S2.4. Error bars in bar graphs indicate the 95% confidence interval (based on t-statistics) and dotted grey lines indicate the borders between the different motional classes and the theoretical maximum of one. Significances of the differences were assessed using a Monte Carlo approach in which the order parameters were resampled based on Student's t-distributions. Significances of 5% and below are highlighted by a red star. Significances of 10% and below are marked by a red star in parentheses.

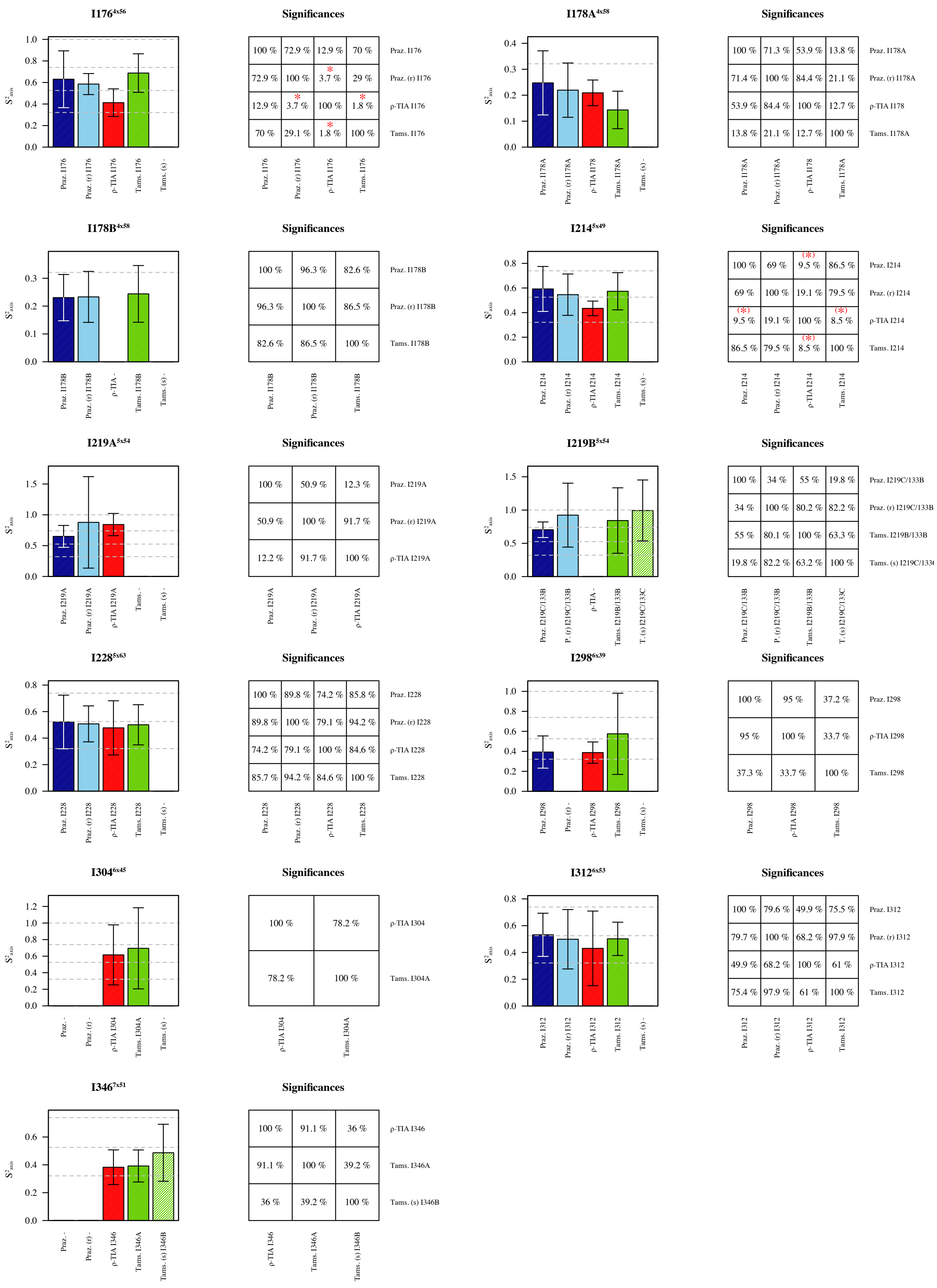


**Figure S4.4.2.** Comparison of Ile δ-methyl order parameters obtained with inverse agonists at 320 K (continuation of Figure S4.4.1). For details and explanation of symbols see Fig S4.4.1.

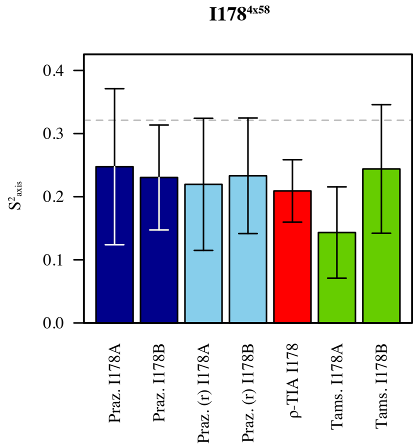


**Significances**

| | Praz. I178A | Praz. I178B | Praz. (r) I178A | Praz. (r) I178B | ρ-TIA I178 | Tams. I178A | Tams. I178B |
|---|---|---|---|---|---|---|---|
| Praz. I178A | 100 % | 80.8 % | 71.4 % | 84.2 % | 53.8 % | 13.8 % | 96.2 % |
| Praz. I178B | 80.8 % | 100 % | 86 % | 96.3 % | 63.8 % | 10.9 % | 82.6 % |
| Praz. (r) I178A | 71.3 % | 86 % | 100 % | 83.3 % | 84.4 % | 21.1 % | 72 % |
| Praz. (r) I178B | 84.2 % | 96.3 % | 83.3 % | 100 % | 61.7 % | 11.7 % | 86.6 % |
| ρ-TIA I178 | 53.9 % | 63.8 % | 84.4 % | 61.7 % | 100 % | 12.6 % | 51 % |
| Tams. I178A | 13.7 % | 10.9 % | 21.1 % | 11.7 % | 12.7 % | 100 % | 10.3 % |
| Tams. I178B | 96.2 % | 82.6 % | 72 % | 86.5 % | 51 % | 10.3 % | 100 % |

**Figure S4.4.3.** Comparison of I178$^{4x58}$ δ-methyl order parameters obtained with inverse agonists at 320 K. I178$^{4x58}$ is present in [$^{13}$C,$^{1}$H]-HSQC spectra with two peaks, except in the spectra recorded with ρ-TIA. Order parameters obtained from the signal with the smaller carbon shift are labeled with A and values obtained from the second peak with B. Error bars indicate the 95% confidence interval (based on t-statistics) and dotted grey lines indicate the borders between the J'- and the J-class. Significances of the differences were assessed using a Monte Carlo approach in which the order parameters were resampled based on a Student's t-distribution.

#### 4.5 Correlations with $\alpha_{1B}$-AR-B1D1 Ile $S^2_{axis}$

The strength of correlations does not report directly on how significant a given correlation is. However, it is less plausible to obtain a strong correlation than a weak one by random chance given the same sample size. The probability of obtaining a correlation of a certain strength or stronger by chance can be described by p-values. All correlations with $\alpha_{1B}$-AR-B1D1 are significant, except the correlations of Ile $S^2_{axis}$ of ρ-TIA-bound $\alpha_{1B}$-AR-B1D1 to side-chain packing and exposure at the protein surface (Figure S4.5.1 and S4.5.2). Ile $S^2_{axis}$ of ρ-TIA-bound $\alpha_{1B}$-AR-B1D1 showed generally weaker correlations to structural properties than the orthosteric ligands (Figure S4.5.1 and S4.5.2). The decrease in correlations might hint at larger differences in structure between ρ-TIA-bound $\alpha_{1B}$-AR-B1D1 and (+)-cyclazosin-bound $\alpha_{1B}$-AR (PDB entry 7B6W), than between either prazosin- or tamsulosin-bound $\alpha_{1B}$-AR-B1D1 and (+)-cyclazosin-bound $\alpha_{1B}$-AR. The reduced correlation of Ile $S^2_{axis}$ of ρ-TIA-bound $\alpha_{1B}$-AR-B1D1 to the protein z-axis most likely reflects changes in side-chain packing and surface exposure. These three structural properties are not independent of one another and show correlations of −0.75 (Spearman's ρ) between z-axis position and packing, of 0.65 between z-axis position and surface exposure, and of −0.89 between packing and surface exposure. Therefore, a decrease in the correlation with one property is likely to affect the strength of the correlation with another property.

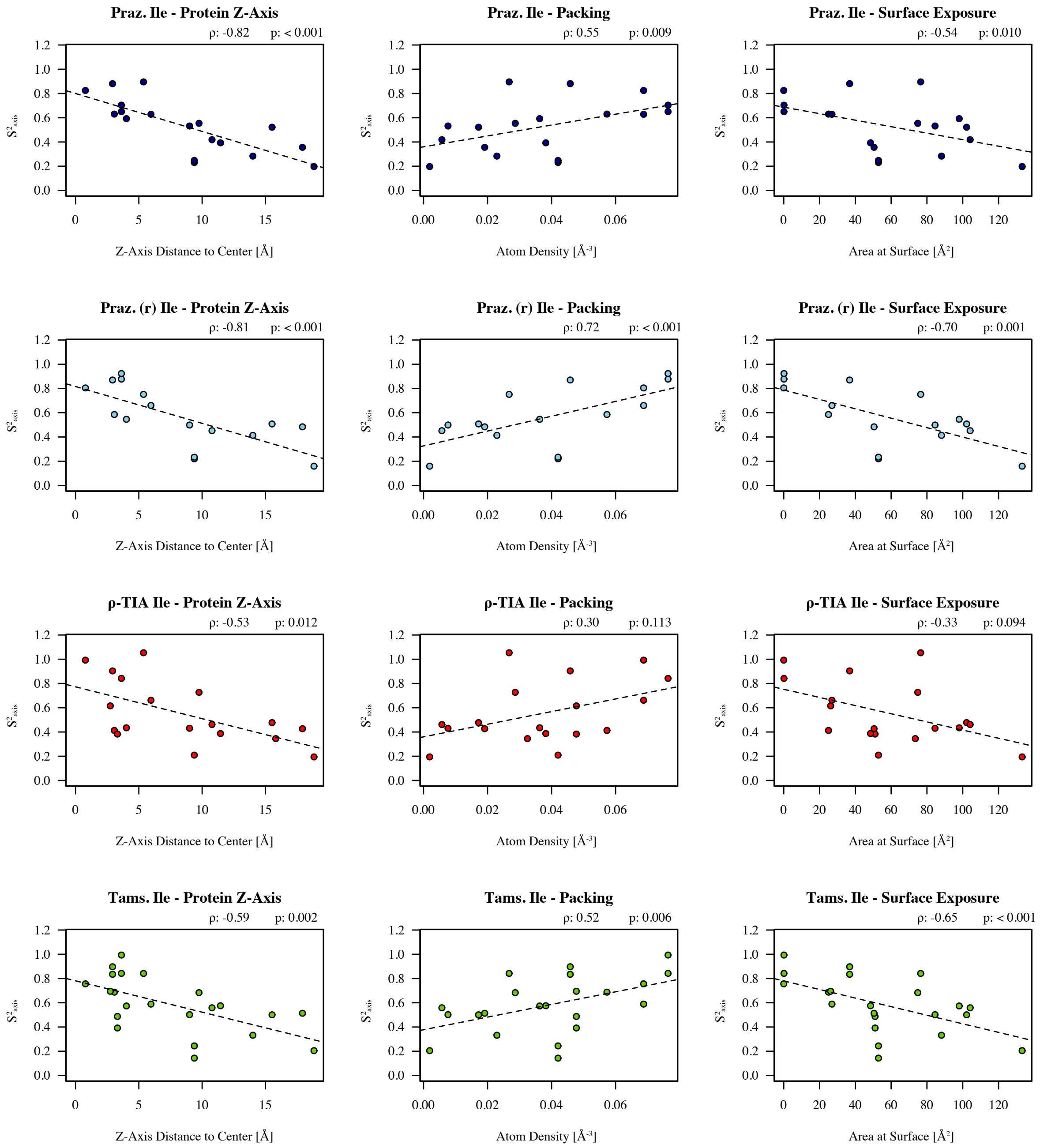


**Figure S4.5.1.** Correlations of Ile methyl order parameters of $\alpha_{1B}$-AR-B1D1 to the structural properties of $\alpha_{1B}$-AR (PDB entry 7B6W). Methyl order parameters are given separately for every ligand dataset. Spearman's rank correlation coefficients (ρ) and the associated p-value are given at the upper right corner of each plot. P-values are based on t-statistics and one-tailed distributions. Dashed trendlines are based on linear regression.

**Significances - Ile**

| | Z-Position | Packing | Surface Exp. |
|---|---|---|---|
| Praz. | *** -0.82 | ** 0.55 | ** -0.54 |
| Praz. (r) | *** -0.81 | *** 0.72 | ** -0.7 |
| ρ-TIA | * -0.53 | 0.30 | -0.33 |
| Tams. | ** -0.59 | ** 0.52 | *** -0.65 |

**Figure S4.5.2.** Spearman's ρ for the correlations between Ile methyl order parameters and the three structural properties (taken from Figure S4.5.1). Significances are based on t-statistics and one-tailed distributions. Probabilities below 5%, 1% and 0.1% are reported by one, two and three stars, respectively.

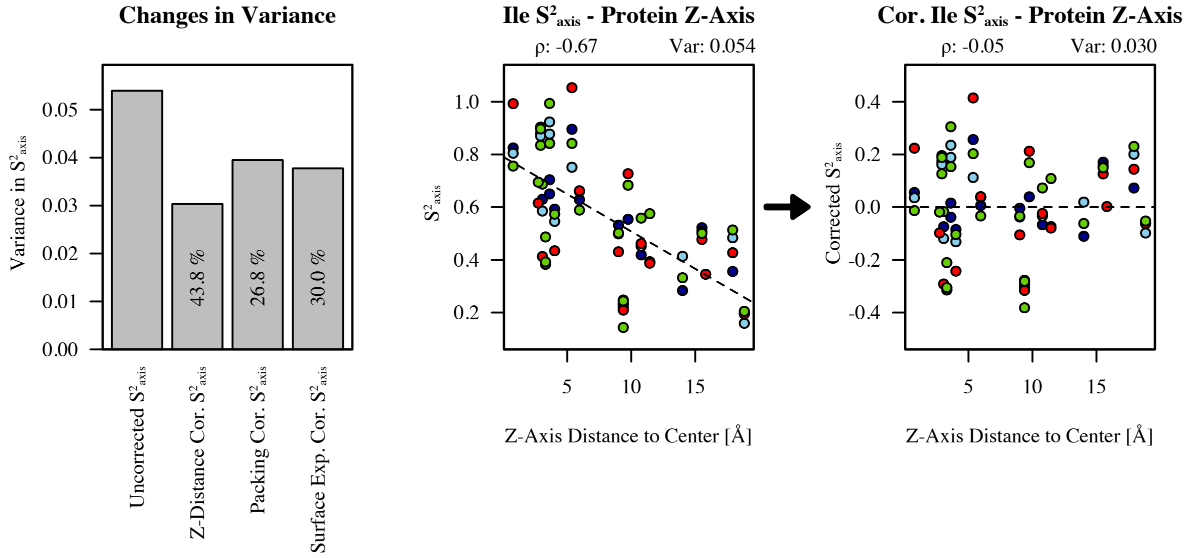


**Figure S4.5.3.** Amount of variation that can be explained by linear models. If we assume a linear relationship between order parameters and the structural properties, then it is possible to subtract the linear model to remove the "influence" of e.g. the protein z-axis (shown in the scatter plots; left: true $S^2_{axis}$, right: corrected $S^2_{axis}$). If a correlation is present, this will reduce the variation between the data points, since one factor that contributes to variations is now removed. The amount of variation that can be explained by this factor, is measured as the change in variance (labeled in percent in the bar graph). This percentage corresponds to $r^2$ (squared Pearson correlation coefficient r). Scatter plots contain all ligand data sets. Spearman's ρ values and variances (Var) of order parameters are displayed above the figures.

##### 4.6 Correlations with microbial rhodopsin Ile and Leu $S^2_{axis}$

Correlations between methyl order parameters and structural properties of the microbial rhodopsins (mRs) were calculated based on PDB entry 5ZIM for bacteriorhodopsin (bR) and PDB entry 1H68 for sensory rhodopsin (pSRII). Correlations for Ile and Leu methyl order parameters are shown in Figure S4.6.1 and S4.6.3 respectively. In terms of side-chain packing and surface exposure, bR shows correlations for Ile and Leu residues that are similar to the correlations with Ile in $\alpha_{1B}$-AR-B1D1. The same correlations are present for pSRII as well but tend to be weaker. Due to the small number of Ile residues, already a single residue can have a large impact on the overall statistics. If I177 and I197 are removed from the pSRII-micelle data set and I197 is removed from the pSRII-bicelle data set (which lacks I177 data), then the correlations resemble the ones found for bR and $\alpha_{1B}$-AR-B1D1 (Figure S4.6.2). I177 and I197 are buried residues in close proximity of one another in the extracellular half of pSRII. Correlations of Val order parameters to side-chain packing and surface exposure are generally absent or too weak to be detected for both mRs.

In bR, correlations between side-chain dynamics and protein z-axis distance to the center plane were mostly absent or too weak to be detected. Interestingly, strong correlations between Ile methyl dynamics and methyl position along the protein z-axis (i.e. distance to the extracellular side), were, however, present.

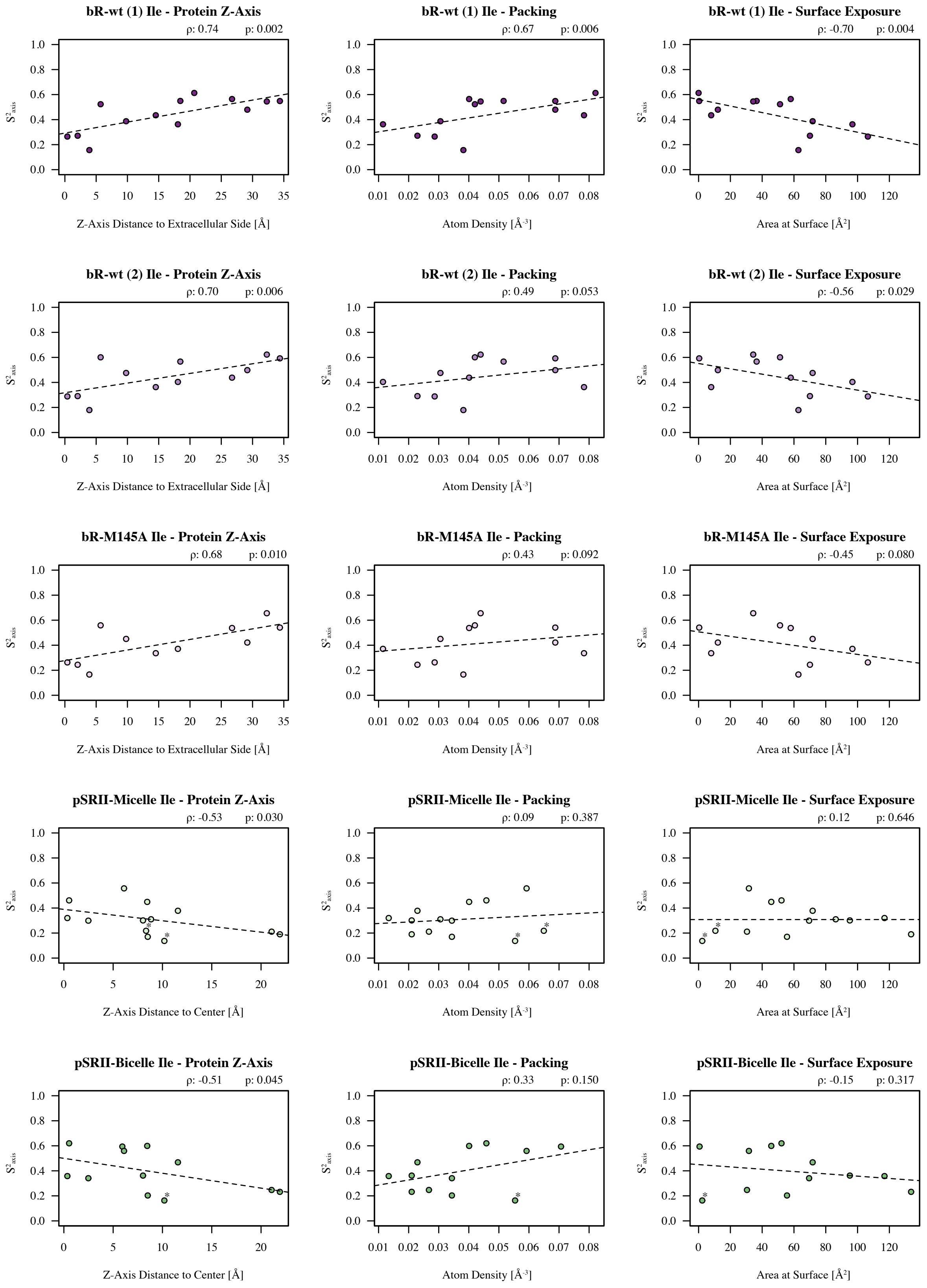


**Figure S4.6.1.** Correlations of Ile methyl order parameters to structural properties of the side chains in bR and pSRII. Order parameters are given separately for every data set. Spearman's rank correlation coefficients (ρ) and the associated p-value are given at the upper right corner of each plot. P-values are based on t-statistics and one-tailed distributions. Dashed trendlines are based on linear regression. I177 and I197 in pSRII are marked by stars.

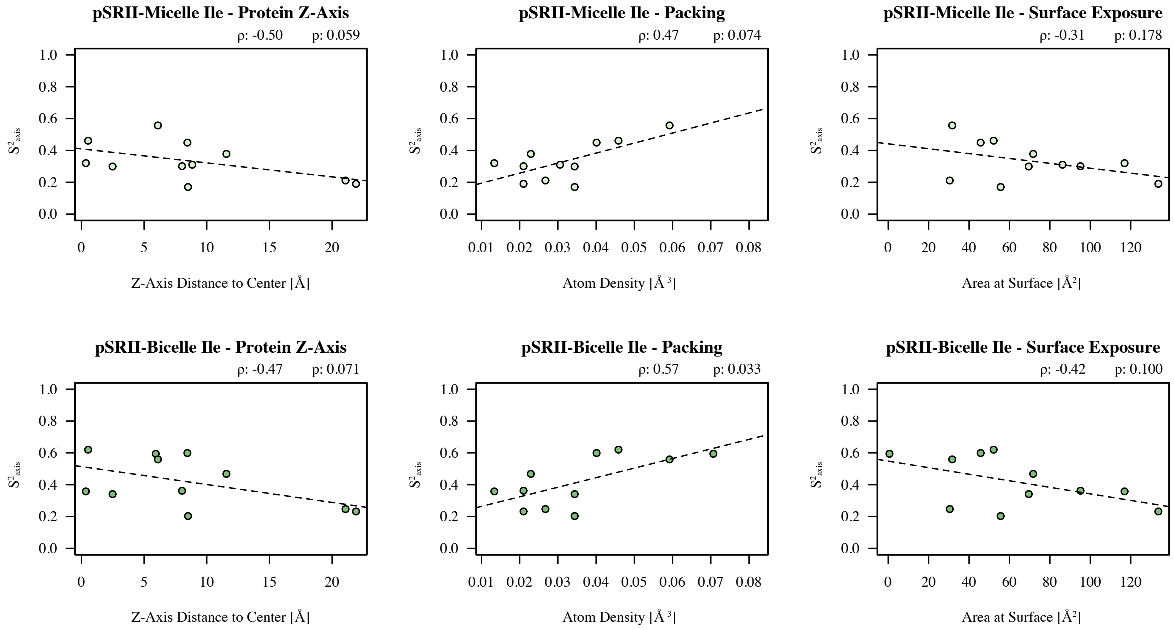


**Figure S4.6.2.** Correlations of pSRII Ile methyl order parameters to structural properties. Plots correspond to Figure S4.6.1 but exclude I177 and I197 data.

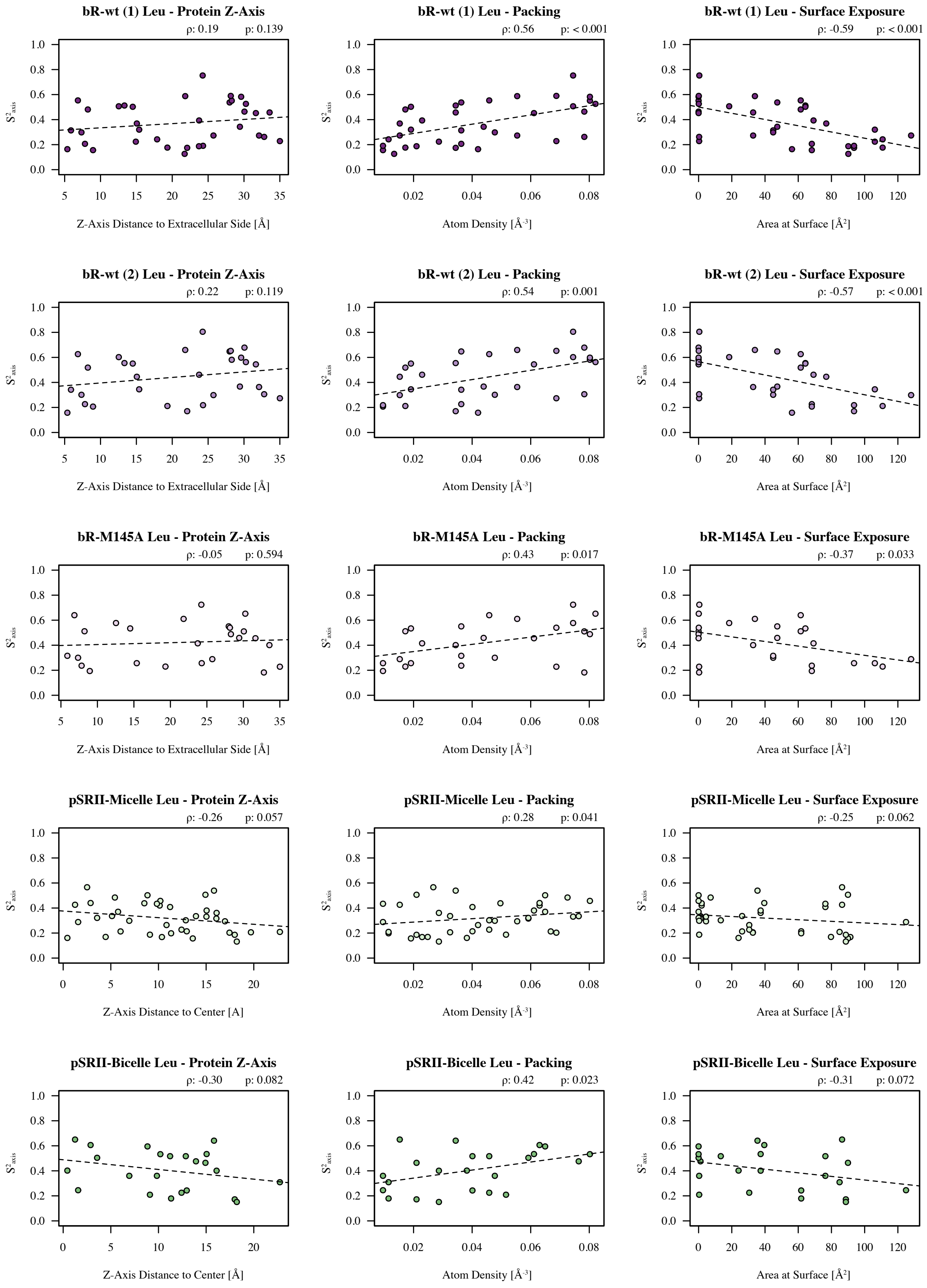


**Figure S4.6.3.** Correlations of Leu methyl order parameters to structural properties of the side chains in bR and pSRII. Methyl order parameters are given separately for every data set. Spearman's rank correlation coefficients (ρ) and the associated p-value are given at the upper right corner of each plot. P-values are based on t-statistics and one-tailed distributions. Dashed trendlines are based on linear regression.
