## Supplementary Data 2 for "Side-Chain Dynamics of the α1B-Adrenergic Receptor determined by NMR via Methyl Relaxation"

**Side-Chain Dynamics of the $\alpha_{1B}$-Adrenergic Receptor determined by NMR via Methyl Relaxation**

Christian Baumann, Wan-Chin Chiang, Renato Valsecchi, Simon Jurt, Mattia Deluigi, Matthias Schuster, Andreas Plückthun, Oliver Zerbe

**This appendix contains Ile assignments and dynamic data with relevant parameters:**


| | | | |
|---|---|---|---|
| 1. Ile assignments at 320 K: | Prazosin | Table S1 | 2 |
| | ρ-TIA | Table S2 | 3 |
| | Tamsulosin | Table S3 | 4 |
| 2. Ile assignments at 298 K: | Prazosin | Table S4 | 5 |
| | Apo | Table S5 | 6 |
| 3. Order parameters ($S^2_{axis}$): | Prazosin | Table S6 | 7 |
| | Prazosin (rep.) | Table S7 | 9 |
| | ρ-TIA | Table S8 | 11 |
| | Tamsulosin | Table S9 | 13 |
| 4. Relative Dynamics (η): | Prazosin | Table S10 | 15 |
| | Apo | Table S11 | 17 |
| 5. TRACT data: | Prazosin | Table S12 | 19 |
| | ρ-TIA | Table S13 | 21 |

**Table S1. Assignments** of Ile δ-methyl groups of the $\alpha_{1B}$-AR-B1D1 binding **prazosin** at **320 K**. Ile assignments in parentheses indicate less likely alternative assignments. GPCRdb numbering for the assigned residues is given in the second column.

| **Residue** | **GPCRdb** | **HD1 [ppm]** | **CD1 [ppm]** |
|---|---|---|---|
| I42 | 1x29 | 0.856 | 12.967 |
| I46 | 1x33 | 0.822 | 13.727 |
| I56 | 1x43 | 0.686 | 13.425 |
| I60 | 1x47 | 0.461 | 13.978 |
| I67 | 1x54 | 0.784 | 14.153 |
| I133A | 3x40 | 0.695 | 14.509 |
| I141 | 3x48 | 0.688 | 14.206 |
| I145 | 3x52 | 0.881 | 13.697 |
| I163 | 4x43 | 0.667 | 12.533 |
| I176 | 4x56 | 0.638 | 12.960 |
| I178A | 4x58 | 0.764 | 12.074 |
| I178B | 4x58 | 0.774 | 12.297 |
| I214 | 5x49 | 0.672 | 12.783 |
| I219A | 5x54 | 0.628 | 14.256 |
| I219B | 5x54 | 0.635 | 14.707 |
| I219C (I133B) | 5x54 (3x40) | 0.646 | 14.798 |
| I228 | 5x63 | 0.898 | 13.435 |
| I298 | 6x39 | 0.259 | 12.477 |
| I304 | 6x45 | 0.705 | 14.482 |
| I312 | 6x53 | 0.697 | 13.060 |
| I346 | 7x51 | 0.800 | 12.451 |

**Table S2. Assignments** of Ile δ-methyl groups of the $\alpha_{1B}$-AR-B1D1 binding **ρ-TIA** at **320 K**. Ile assignments in parentheses indicate less likely alternative assignments. Ile assignments separated by slashes are equally likely. GPCRdb numbering for the assigned residues is given in the second column.

| Residue | GPCRdb | HD1 [ppm] | CD1 [ppm] |
|---|---|---|---|
| I42 | 1x29 | 0.851 | 13.066 |
| I46 | 1x33 | 0.81 | 13.700 |
| I56 | 1x43 | 0.696 | 13.765 |
| I60 | 1x47 | 0.435 | 13.781 |
| I67 | 1x54 | 0.790 | 14.191 |
| I84/I139 | 2x43/3x46 | 0.732 | 13.104 |
| I120 | 3x27 | 0.793 | 13.321 |
| I133 (I219B) | 3x40 (5x54) | 0.705 | 14.736 |
| I141 | 3x48 | 0.671 | 14.111 |
| I145 | 3x52 | 0.890 | 13.694 |
| I163 | 4x43 | 0.671 | 12.368 |
| I176 | 4x56 | 0.550 | 12.659 |
| I178 | 4x58 | 0.754 | 12.029 |
| I214 | 5x49 | 0.652 | 12.531 |
| I219A | 5x54 | 0.619 | 14.422 |
| I228 | 5x63 | 0.896 | 13.519 |
| I298 | 6x39 | 0.250 | 12.516 |
| I304 | 6x45 | 0.739 | 14.050 |
| I312 | 6x53 | 0.898 | 12.078 |
| I346 | 7x51 | 0.858 | 12.381 |

**Table S3. Assignments** of Ile δ-methyl groups of the $\alpha_{1B}$-AR-B1D1 binding **tamsulosin** at **320 K**. Ile assignments in parentheses indicate less likely alternative assignments. GPCRdb numbering for the assigned residues is given in the second column.

| Residue | GPCRdb | HD1 [ppm] | CD1 [ppm] |
|---|---|---|---|
| I42 | 1x29 | 0.851 | 12.965 |
| I46 | 1x33 | 0.803 | 13.756 |
| I56 | 1x43 | 0.604 | 13.551 |
| I60A | 1x47 | 0.394 | 13.812 |
| I60B | 1x47 | 0.374 | 13.909 |
| I67 | 1x54 | 0.782 | 14.104 |
| I84 | 2x43 | 0.824 | 13.065 |
| I133A (I219D) | 3x27 (5x54) | 0.642 | 14.608 |
| I141 | 3x40 | 0.678 | 14.340 |
| I145 | 3x52 | 0.883 | 13.772 |
| I163 | 4x43 | 0.667 | 12.525 |
| I176 | 4x56 | 0.662 | 13.118 |
| I178A | 4x58 | 0.768 | 11.772 |
| I178B | 4x58 | 0.777 | 12.038 |
| I214 | 5x49 | 0.666 | 12.749 |
| I219A | 5x54 | 0.583 | 14.237 |
| I219B (I133B) | 5x54 (3x27) | 0.618 | 14.702 |
| I219C (I133C) | 5x54 (3x27) | 0.629 | 14.764 |
| I228 | 5x63 | 0.892 | 13.476 |
| I298 | 6x39 | 0.244 | 12.415 |
| I304A | 6x45 | 0.699 | 14.084 |
| I304B | 6x45 | 0.69 | 14.407 |
| I312 | 6x53 | 0.693 | 13.122 |
| I346A | 7x51 | 0.881 | 12.471 |
| I346B | 7x51 | 0.902 | 12.563 |

**Table S4. Assignments** of Ile δ-methyl groups of the $\alpha_{1B}$-AR-B1D1 binding **prazosin** at **298 K**. Ile assignments in parentheses indicate less likely alternative assignments. Ile assignments separated by slashes are equally likely. GPCRdb numbering for the assigned residues is given in the second column.

| Residue | GPCRdb | HD1 [ppm] | CD1 [ppm] |
|---|---|---|---|
| I42 | 1x29 | 0.858 | 12.802 |
| I46 | 1x33 | 0.823 | 13.916 |
| I56 | 1x43 | 0.686 | 13.542 |
| I60 | 1x47 | 0.445 | 14.163 |
| I67 | 1x54 | 0.772 | 14.163 |
| I141 | 3x48 | 0.670 | 14.104 |
| I145 | 3x52 | 0.888 | 13.844 |
| I163 | 4x43 | 0.673 | 12.178 |
| I176 | 4x56 | 0.653 | 13.206 |
| I178A | 4x58 | 0.754 | 11.844 |
| I178B | 4x58 | 0.764 | 12.024 |
| I214 | 5x49 | 0.679 | 12.970 |
| I219A | 5x54 | 0.626 | 14.219 |
| I219B | 5x54 | 0.628 | 14.668 |
| I219C (I133B) | 5x54 (3x40) | 0.644 | 14.828 |
| I228 | 5x63 | 0.894 | 13.584 |
| I298 | 6x39 | 0.196 | 12.503 |
| I304/I133A | 6x45/3x40 | 0.696 | 14.558 |
| I312 | 6x53 | 0.690 | 13.177 |

**Table S5. Assignments** of Ile δ-methyl groups of the **apo** $\alpha_{1B}$-AR-B1D1 at **298 K**. Ile assignments in parentheses indicate less likely alternative assignments. Ile assignments separated by slashes are equally likely. GPCRdb numbering for the assigned residues is given in the second column.

| Residue | GPCRdb | HD1 [ppm] | CD1 [ppm] |
|---|---|---|---|
| I42 | 1x29 | 0.851 | 12.860 |
| I46 | 1x33 | 0.813 | 13.924 |
| I56 | 1x43 | 0.668 | 13.552 |
| I60 | 1x47 | 0.395 | 14.083 |
| I67 | 1x54 | 0.763 | 14.187 |
| I145 | 3x52 | 0.885 | 13.832 |
| I163 | 4x43 | 0.663 | 12.190 |
| I176/I214 | 4x56/5x49 | 0.641 | 12.881 |
| I178 | 4x58 | 0.761 | 11.806 |
| I219A | 5x54 | 0.602 | 14.364 |
| I219B | 5x54 | 0.605 | 14.637 |
| I219C (I133B) | 5x54 (3x40) | 0.605 | 14.824 |
| I228 | 5x63 | 0.892 | 13.629 |
| I298 | 6x39 | 0.179 | 12.427 |
| I304/I133A | 6x45/3x40 | 0.694 | 14.555 |
| I312 | 6x53 | 0.710 | 13.104 |

**Table S6. Order parameters** of the $\alpha_{1B}$-AR-B1D1 binding **prazosin**. Data was recorded at **320 K** at 700 MHz. Standard errors of order parameters ($S^2_{axis}$) were estimated using 1 000 Monte Carlo samplings based on the $\eta$ and the $\tau_c$ standard errors. Amino acid assignments that are not fully unambiguous are marked with a star. Ile assignments in parentheses indicate less likely alternative assignments. The degree of freedom (df) indicates if data points were excluded to fit $\eta$ and $\delta$. A df of 9 means that all data points (11) were included, 8 means that one point and 7 means that two data points were excluded from the fit.

| Peak Nr. | Residue | Amino Acid | $S^2_{axis}$ | $S^2_{axis}$ SE | $\eta$ [$s^{-1}$] | $\eta$ SE [$s^{-1}$] | $\delta$ [$s^{-1}$] | $\delta$ SE [$s^{-1}$] | RSE | df |
|---|---|---|---|---|---|---|---|---|---|---|
| 1 | 42 | Ile | 0.197 | 0.023 | 26.350 | 3.026 | -23.100 | 3.937 | 0.026 | 9 |
| 2 | 46 | Ile | 0.419 | 0.030 | 56.016 | 4.163 | -35.678 | 4.165 | 0.020 | 7 |
| 3 | 56 | Ile | 0.895 | 0.156 | 119.794 | 21.008 | -82.333 | 17.711 | 0.043 | 9 |
| 4 | 60 | Ile | 0.881 | 0.145 | 117.823 | 18.739 | -66.113 | 13.578 | 0.045 | 9 |
| 5 | 67 | Ile | 0.628 | 0.141 | 84.071 | 18.619 | -60.800 | 16.947 | 0.057 | 9 |
| 6 | 133A | Ile | 0.825 | 0.179 | 110.317 | 23.520 | -63.793 | 17.560 | 0.060 | 9 |
| 7 | 141 | Ile | 0.554 | 0.067 | 74.090 | 8.813 | -68.418 | 10.176 | 0.025 | 8 |
| 8 | 145 | Ile | 0.356 | 0.022 | 47.595 | 2.897 | -52.207 | 4.306 | 0.011 | 7 |
| 9 | 163 | Ile | 0.284 | 0.027 | 37.928 | 3.625 | -9.461 | 2.788 | 0.036 | 7 |
| 10 | 176 | Ile | 0.630 | 0.114 | 84.267 | 15.775 | -72.878 | 17.039 | 0.041 | 8 |
| 11 | 178A | Ile | 0.248 | 0.055 | 33.113 | 7.399 | -29.994 | 9.230 | 0.050 | 9 |
| 12 | 178B | Ile | 0.230 | 0.036 | 30.830 | 4.730 | -17.262 | 4.938 | 0.044 | 8 |
| 13 | 214 | Ile | 0.592 | 0.079 | 79.235 | 10.717 | -64.878 | 11.049 | 0.032 | 8 |
| 14 | 219A | Ile | 0.650 | 0.077 | 86.971 | 10.477 | -52.963 | 8.688 | 0.034 | 8 |
| 15 | 219C (133B) | Ile | 0.704 | 0.051 | 94.156 | 6.667 | -54.486 | 4.804 | 0.018 | 8 |
| 16 | 228 | Ile | 0.522 | 0.088 | 69.803 | 11.749 | -66.694 | 14.037 | 0.035 | 8 |
| 17 | 298 | Ile | 0.393 | 0.068 | 52.560 | 9.268 | -41.095 | 10.695 | 0.041 | 7 |
| 18 | 312 | Ile | 0.532 | 0.071 | 71.133 | 9.217 | -57.548 | 9.317 | 0.031 | 9 |
| 19 | | Ile | 0.582 | 0.110 | 77.920 | 14.823 | -50.297 | 13.010 | 0.052 | 8 |
| 20 | | Ile | 0.471 | 0.072 | 63.038 | 9.408 | -47.109 | 9.076 | 0.038 | 9 |
| 21 | | Ile | 0.617 | 0.066 | 82.602 | 8.575 | -71.866 | 9.055 | 0.023 | 9 |
| 22 | | Leu | 0.533 | 0.078 | 71.256 | 10.474 | -62.012 | 11.600 | 0.033 | 8 |
| 23 | | Leu | 0.429 | 0.021 | 57.354 | 2.723 | -44.377 | 3.048 | 0.011 | 7 |
| 24 | | Leu | 0.583 | 0.077 | 77.928 | 10.299 | -57.897 | 9.959 | 0.033 | 8 |
| 25 | | Leu | 0.385 | 0.108 | 51.453 | 14.157 | -62.078 | 20.515 | 0.050 | 9 |
| 26 | | Leu | 0.524 | 0.068 | 70.067 | 8.924 | -46.031 | 7.720 | 0.035 | 9 |
| 27 | | Leu | 0.813 | 0.169 | 108.727 | 22.020 | -84.237 | 21.180 | 0.046 | 8 |
| 28 | | Leu | 0.415 | 0.033 | 55.493 | 4.239 | -32.698 | 3.530 | 0.023 | 9 |
| 29 | | Leu | 0.669 | 0.155 | 89.535 | 20.879 | -78.392 | 23.480 | 0.050 | 7 |
| 30 | | Leu | 0.416 | 0.053 | 55.611 | 6.991 | -22.368 | 5.010 | 0.043 | 8 |
| 31 | | Leu | 0.842 | 0.118 | 112.706 | 15.257 | -92.377 | 15.275 | 0.030 | 8 |
| 32 | | Leu | 0.415 | 0.045 | 55.470 | 6.234 | -52.785 | 7.372 | 0.024 | 9 |
| 33 | | Leu | 0.886 | 0.200 | 118.558 | 26.963 | -90.613 | 24.767 | 0.052 | 9 |
| 34 | | Leu | 0.439 | 0.045 | 58.671 | 6.005 | -40.130 | 5.485 | 0.028 | 9 |
| 35 | | Leu | 0.379 | 0.034 | 50.700 | 4.443 | -44.528 | 4.994 | 0.020 | 9 |
| 36 | | Leu | 0.567 | 0.064 | 75.913 | 8.380 | -56.946 | 8.225 | 0.027 | 8 |
| 37 | | Leu* | 0.646 | 0.075 | 86.399 | 10.344 | -75.527 | 11.235 | 0.026 | 8 |
| 38 | | Leu | 0.388 | 0.055 | 51.940 | 7.290 | -24.286 | 5.352 | 0.046 | 9 |
| 39 | | Leu | 0.479 | 0.044 | 64.019 | 5.777 | -45.984 | 5.641 | 0.023 | 8 |
| 40 | | Leu | 0.203 | 0.008 | 27.127 | 0.917 | -15.794 | 1.071 | 0.009 | 8 |
| 41 | | Leu | 0.248 | 0.033 | 33.192 | 4.461 | -21.426 | 4.487 | 0.037 | 9 |
| 42 | | Leu | 0.145 | 0.009 | 19.350 | 1.154 | -12.295 | 1.705 | 0.013 | 8 |
| 43 | | Leu | 0.398 | 0.042 | 53.201 | 5.420 | -33.866 | 5.069 | 0.028 | 8 |
| 44 | | Leu | 0.438 | 0.054 | 58.599 | 7.206 | -39.682 | 6.537 | 0.034 | 9 |
| 45 | | Leu | 0.629 | 0.093 | 84.198 | 11.765 | -93.521 | 15.234 | 0.025 | 9 |
| 46 | | Leu | 0.436 | 0.040 | 58.295 | 5.160 | -44.408 | 5.104 | 0.022 | 9 |
| 47 | | Leu | 0.198 | 0.009 | 26.516 | 1.056 | -18.354 | 1.194 | 0.010 | 9 |
| 48 | | Leu* | 0.791 | 0.188 | 105.768 | 23.946 | -100.564 | 27.243 | 0.044 | 8 |
| 49 | | Leu | 0.145 | 0.007 | 19.352 | 0.928 | -19.819 | 1.475 | 0.010 | 9 |
| 50 | | Leu | 0.095 | 0.005 | 12.775 | 0.574 | -9.853 | 0.979 | 0.009 | 9 |
| 51 | | Leu | 0.259 | 0.026 | 34.616 | 3.472 | -16.450 | 2.928 | 0.032 | 9 |
| 52 | | Leu | 0.454 | 0.099 | 60.748 | 13.483 | -42.182 | 12.488 | 0.060 | 8 |
| 53 | | Leu | 0.777 | 0.212 | 103.980 | 27.842 | -83.308 | 26.911 | 0.061 | 9 |
| 54 | | Leu | 0.506 | 0.049 | 67.648 | 6.476 | -23.875 | 3.842 | 0.036 | 9 |
| 55 | | Leu | 0.426 | 0.030 | 57.051 | 3.806 | -38.514 | 3.460 | 0.018 | 9 |
| 56 | | Leu | 0.227 | 0.014 | 30.361 | 1.785 | -26.124 | 2.201 | 0.014 | 9 |
| 57 | | Leu | 0.833 | 0.245 | 111.463 | 33.416 | -97.800 | 34.589 | 0.063 | 9 |
| 58 | | Leu | 0.144 | 0.010 | 19.219 | 1.340 | -12.906 | 1.718 | 0.018 | 9 |

| | | | | | | | | | |
|---|---|---|---|---|---|---|---|---|---|
| 59 | Leu | 0.396 | 0.041 | 52.945 | 5.440 | -28.724 | 4.637 | 0.031 | 8 |
| 60 | Leu | 0.239 | 0.019 | 31.928 | 2.575 | -24.320 | 2.895 | 0.020 | 9 |
| 61 | Leu | 0.186 | 0.014 | 24.944 | 1.836 | -14.953 | 1.978 | 0.021 | 9 |
| 62 | Leu | 0.200 | 0.009 | 26.778 | 1.117 | -12.084 | 1.029 | 0.013 | 9 |
| 63 | Leu | 0.440 | 0.055 | 58.836 | 7.382 | -66.420 | 10.239 | 0.024 | 8 |
| 64 | Leu | 0.531 | 0.062 | 71.009 | 8.618 | -55.770 | 8.833 | 0.029 | 8 |
| 65 | Leu | 0.593 | 0.070 | 79.363 | 9.326 | -63.967 | 9.295 | 0.028 | 9 |
| 66 | Leu | 0.335 | 0.049 | 44.801 | 6.459 | -43.650 | 8.010 | 0.031 | 9 |
| 67 | Leu | 0.354 | 0.043 | 47.423 | 5.652 | -42.193 | 6.481 | 0.027 | 9 |
| 68 | Leu | 0.502 | 0.085 | 67.213 | 11.046 | -52.721 | 10.970 | 0.041 | 9 |
| 69 | Leu | 0.197 | 0.013 | 26.299 | 1.654 | -11.287 | 1.506 | 0.020 | 9 |
| 70 | Leu | 0.229 | 0.035 | 30.641 | 4.513 | -13.881 | 4.314 | 0.046 | 8 |
| 71 | Leu | 0.422 | 0.043 | 56.512 | 5.847 | -38.181 | 5.327 | 0.028 | 9 |
| 72 | Leu | 0.493 | 0.059 | 65.970 | 7.882 | -76.627 | 11.531 | 0.021 | 7 |
| 73 | Leu | 0.275 | 0.029 | 36.822 | 3.861 | -29.262 | 4.295 | 0.026 | 9 |
| 74 | Leu | 0.837 | 0.105 | 112.018 | 13.726 | -96.237 | 14.285 | 0.026 | 8 |
| 75 | Leu | 0.560 | 0.041 | 74.883 | 5.369 | -33.337 | 3.550 | 0.024 | 9 |
| 76 | Leu | 0.685 | 0.243 | 91.611 | 32.676 | -113.630 | 47.208 | 0.057 | 8 |
| 77 | Leu | 0.368 | 0.036 | 49.192 | 4.696 | -41.062 | 5.102 | 0.023 | 9 |
| 78 | Leu | 0.487 | 0.040 | 65.158 | 5.162 | -45.037 | 4.892 | 0.021 | 8 |
| 79 | Leu | 0.320 | 0.011 | 42.772 | 1.349 | -31.158 | 1.603 | 0.007 | 7 |
| 80 | Leu | 0.643 | 0.160 | 86.024 | 21.741 | -73.753 | 23.267 | 0.056 | 8 |
| 81 | Val | 0.278 | 0.031 | 37.172 | 4.038 | -28.607 | 4.385 | 0.027 | 9 |
| 82 | Val | 0.951 | 0.228 | 127.243 | 29.714 | -105.824 | 29.058 | 0.050 | 9 |
| 83 | Val | 0.679 | 0.053 | 90.856 | 6.841 | -54.145 | 5.553 | 0.021 | 8 |
| 84 | Val | 0.740 | 0.080 | 99.046 | 10.669 | -70.343 | 9.414 | 0.027 | 9 |
| 85 | Val | 0.442 | 0.067 | 59.094 | 9.095 | -57.061 | 10.804 | 0.033 | 9 |
| 86 | Val | 0.683 | 0.145 | 91.388 | 19.732 | -49.494 | 15.871 | 0.063 | 7 |
| 87 | Val | 0.750 | 0.122 | 100.314 | 16.596 | -87.646 | 17.723 | 0.035 | 8 |
| 88 | Val | 0.782 | 0.050 | 104.676 | 6.656 | -89.531 | 6.954 | 0.014 | 8 |
| 89 | Val | 0.345 | 0.035 | 46.136 | 4.468 | -37.412 | 4.804 | 0.024 | 9 |
| 90 | Val | 1.044 | 0.325 | 139.644 | 43.462 | -119.791 | 43.254 | 0.063 | 9 |
| 91 | Val | 0.665 | 0.095 | 88.926 | 12.558 | -54.779 | 10.070 | 0.040 | 9 |
| 92 | Val | 0.847 | 0.257 | 113.333 | 35.054 | -94.732 | 34.797 | 0.067 | 9 |
| 93 | Val | 0.728 | 0.120 | 97.420 | 15.881 | -84.341 | 16.455 | 0.036 | 9 |
| 94 | Val | 0.278 | 0.045 | 37.211 | 6.112 | -29.754 | 7.255 | 0.039 | 8 |
| 95 | Val | 0.635 | 0.085 | 84.910 | 10.841 | -61.510 | 10.215 | 0.032 | 8 |
| 96 | Val | 0.768 | 0.126 | 102.676 | 16.683 | -71.777 | 14.937 | 0.041 | 8 |
| 97 | Val | 0.658 | 0.080 | 88.090 | 10.926 | -73.523 | 11.417 | 0.028 | 8 |
| 98 | Val | 0.227 | 0.020 | 30.427 | 2.536 | -18.030 | 2.810 | 0.022 | 8 |
| 99 | Val | 0.840 | 0.110 | 112.340 | 14.614 | -74.477 | 12.063 | 0.033 | 9 |
| 100 | Val | 0.637 | 0.113 | 85.220 | 14.626 | -68.260 | 14.393 | 0.041 | 8 |
| 101 | Val | 0.690 | 0.128 | 92.337 | 17.067 | -97.843 | 22.205 | 0.033 | 7 |
| 102 | Val | 0.929 | 0.124 | 124.311 | 17.053 | -108.138 | 17.726 | 0.028 | 8 |

**Table S7. Order parameters** of the $\alpha_{1B}$-AR-B1D1 binding **prazosin** at **320 K**, based on a second measurement of the same sample for which data is shown in table S1. Data was recorded at 320 K at 700 MHz without duplicate recording of the experiments with 8 ms delays. Standard errors of order parameters ($S^2_{axis}$) were estimated using 1 000 Monte Carlo samplings based on the η and the $\tau_c$ standard errors. Amino acid assignments that are not fully unambiguous are marked with a star. Ile assignments in parentheses indicate less likely alternative assignments. The degree of freedom (df) indicates if data points were excluded to fit η and δ. A df of 8 means that all data points (10) were included, 7 means that one point and 6 means that two data points were excluded from the fit.

| Peak Nr. | Residue | Amino Acid | $S^2_{axis}$ | $S^2_{axis}$ SE | η [s$^{-1}$] | η SE [s$^{-1}$] | δ [s$^{-1}$] | δ SE [s$^{-1}$] | RSE | df |
|---|---|---|---|---|---|---|---|---|---|---|
| 1 | 42 | Ile | 0.159 | 0.010 | 21.286 | 1.324 | -15.383 | 1.649 | 0.014 | 8 |
| 2 | 46 | Ile | 0.452 | 0.040 | 60.439 | 5.272 | -45.588 | 5.116 | 0.019 | 7 |
| 3 | 56 | Ile | 0.751 | 0.071 | 100.523 | 9.284 | -54.609 | 6.565 | 0.024 | 7 |
| 4 | 60 | Ile | 0.870 | 0.212 | 116.380 | 27.488 | -66.384 | 19.881 | 0.060 | 7 |
| 5 | 67 | Ile | 0.660 | 0.101 | 88.246 | 13.391 | -57.731 | 10.681 | 0.036 | 8 |
| 6 | 133A | Ile | 0.805 | 0.078 | 107.643 | 10.080 | -68.680 | 7.764 | 0.023 | 8 |
| 7 | 145 | Ile | 0.484 | 0.050 | 64.719 | 6.656 | -70.205 | 8.315 | 0.017 | 8 |
| 8 | 163 | Ile | 0.413 | 0.161 | 55.293 | 22.010 | -100.167 | 44.929 | 0.044 | 7 |
| 9 | 176 | Ile | 0.585 | 0.041 | 78.289 | 5.329 | -53.183 | 4.585 | 0.016 | 7 |
| 10 | 178A | Ile | 0.220 | 0.044 | 29.372 | 6.040 | -29.911 | 8.701 | 0.036 | 7 |
| 11 | 178B | Ile | 0.233 | 0.039 | 31.190 | 5.303 | -23.573 | 5.566 | 0.035 | 7 |
| 12 | 214 | Ile | 0.546 | 0.073 | 73.015 | 9.703 | -51.570 | 8.385 | 0.030 | 8 |
| 13 | 219A | Ile | 0.877 | 0.314 | 117.364 | 40.482 | -109.822 | 43.877 | 0.064 | 7 |
| 14 | 219C (133) | Ile | 0.923 | 0.208 | 123.519 | 28.577 | -72.163 | 20.351 | 0.059 | 8 |
| 15 | 228 | Ile | 0.507 | 0.055 | 67.882 | 7.312 | -63.503 | 8.579 | 0.020 | 6 |
| 16 | 312 | Ile | 0.499 | 0.096 | 66.696 | 13.310 | -51.677 | 12.516 | 0.043 | 8 |
| 17 | | Ile | 0.492 | 0.051 | 65.813 | 6.635 | -37.227 | 5.171 | 0.025 | 7 |
| 18 | | Ile | 0.474 | 0.096 | 63.374 | 12.354 | -26.878 | 8.159 | 0.056 | 7 |
| 19 | | Ile | 0.586 | 0.052 | 78.452 | 6.751 | -62.534 | 6.576 | 0.018 | 7 |
| 20 | | Leu | 0.363 | 0.031 | 48.503 | 4.082 | -42.280 | 4.762 | 0.016 | 6 |
| 21 | | Leu | 0.239 | 0.025 | 31.922 | 3.143 | -18.979 | 2.977 | 0.026 | 8 |
| 22 | | Leu | 0.341 | 0.027 | 45.571 | 3.480 | -31.010 | 3.387 | 0.017 | 7 |
| 23 | | Leu | 0.483 | 0.087 | 64.660 | 11.822 | -48.552 | 11.291 | 0.039 | 7 |
| 24 | | Leu | 0.587 | 0.189 | 78.539 | 25.203 | -68.680 | 26.462 | 0.063 | 7 |
| 25 | | Leu | 0.311 | 0.041 | 41.565 | 5.366 | -36.901 | 6.072 | 0.025 | 8 |
| 26 | | Leu | 0.268 | 0.038 | 35.821 | 4.966 | -33.947 | 6.096 | 0.026 | 8 |
| 27 | | Leu | 0.515 | 0.112 | 68.842 | 14.597 | -49.570 | 13.371 | 0.047 | 7 |
| 28 | | Leu | 0.563 | 0.096 | 75.333 | 12.504 | -32.166 | 7.908 | 0.048 | 7 |
| 29 | | Leu | 0.472 | 0.089 | 63.149 | 11.871 | -48.953 | 11.247 | 0.040 | 8 |
| 30 | | Leu | 0.586 | 0.081 | 78.443 | 10.661 | -62.113 | 10.328 | 0.028 | 7 |
| 31 | | Leu | 0.340 | 0.047 | 45.547 | 6.082 | -38.028 | 6.451 | 0.027 | 8 |
| 32 | | Leu | 0.237 | 0.029 | 31.707 | 4.004 | -19.753 | 4.317 | 0.029 | 7 |
| 33 | | Leu | 0.221 | 0.011 | 29.547 | 1.435 | -21.195 | 1.555 | 0.011 | 8 |
| 34 | | Leu | 0.381 | 0.050 | 50.938 | 6.836 | -37.951 | 6.834 | 0.029 | 7 |
| 35 | | Leu | 0.509 | 0.043 | 68.071 | 5.603 | -32.615 | 3.697 | 0.023 | 8 |
| 36 | | Leu | 0.210 | 0.010 | 28.149 | 1.335 | -12.339 | 1.159 | 0.014 | 8 |
| 37 | | Leu | 0.404 | 0.056 | 54.084 | 7.259 | -43.878 | 7.295 | 0.028 | 8 |
| 38 | | Leu | 0.615 | 0.077 | 82.246 | 9.977 | -63.033 | 9.370 | 0.026 | 7 |
| 39 | | Leu | 0.555 | 0.116 | 74.220 | 15.345 | -60.869 | 15.145 | 0.043 | 7 |
| 40 | | Leu | 0.480 | 0.067 | 64.211 | 8.849 | -53.161 | 9.116 | 0.028 | 7 |
| 41 | | Leu | 0.303 | 0.014 | 40.546 | 1.761 | -12.119 | 1.312 | 0.014 | 6 |
| 42 | | Leu | 0.469 | 0.090 | 62.681 | 12.726 | -44.983 | 11.660 | 0.045 | 7 |
| 43 | | Leu | 0.395 | 0.053 | 52.862 | 7.294 | -42.187 | 7.267 | 0.029 | 8 |
| 44 | | Leu | 0.514 | 0.052 | 68.811 | 6.773 | -21.712 | 3.747 | 0.031 | 7 |
| 45 | | Leu | 0.255 | 0.040 | 34.074 | 5.501 | -29.828 | 6.468 | 0.032 | 8 |
| 46 | | Leu* | 0.686 | 0.125 | 91.774 | 16.162 | -80.430 | 16.785 | 0.035 | 7 |
| 47 | | Leu | 0.188 | 0.016 | 25.127 | 2.063 | -9.709 | 1.815 | 0.026 | 8 |
| 48 | | Leu | 0.398 | 0.037 | 53.264 | 4.960 | -34.116 | 4.202 | 0.022 | 8 |
| 49 | | Leu | 0.629 | 0.060 | 84.114 | 8.013 | -57.965 | 6.691 | 0.022 | 8 |
| 50 | | Leu | 0.637 | 0.136 | 85.170 | 18.614 | -67.570 | 17.911 | 0.046 | 7 |
| 51 | | Leu | 0.140 | 0.011 | 18.764 | 1.413 | -17.778 | 2.141 | 0.015 | 8 |
| 52 | | Leu | 0.452 | 0.116 | 60.424 | 15.921 | -35.368 | 12.958 | 0.065 | 7 |
| 53 | | Leu | 0.259 | 0.030 | 34.640 | 4.034 | -25.407 | 4.303 | 0.026 | 7 |
| 54 | | Leu | 0.430 | 0.041 | 57.583 | 5.347 | -30.410 | 3.905 | 0.025 | 8 |
| 55 | | Leu | 0.095 | 0.005 | 12.670 | 0.613 | -7.457 | 0.955 | 0.011 | 8 |
| 56 | | Leu | 0.450 | 0.070 | 60.163 | 9.341 | -26.055 | 6.345 | 0.044 | 7 |

| | | | | | | | | | |
|---|---|---|---|---|---|---|---|---|---|
| 57 | Leu | 0.349 | 0.049 | 46.726 | 6.326 | -27.931 | 5.645 | 0.033 | 7 |
| 58 | Leu | 0.396 | 0.050 | 52.994 | 6.666 | -41.562 | 6.552 | 0.027 | 8 |
| 59 | Leu | 0.409 | 0.122 | 54.672 | 16.415 | -58.569 | 21.249 | 0.051 | 7 |
| 60 | Leu | 0.259 | 0.022 | 34.673 | 2.942 | -13.637 | 2.127 | 0.024 | 7 |
| 61 | Leu | 0.510 | 0.113 | 68.270 | 15.458 | -50.949 | 14.568 | 0.049 | 7 |
| 62 | Leu | 0.594 | 0.081 | 79.476 | 10.957 | -44.164 | 7.593 | 0.035 | 7 |
| 63 | Leu | 0.436 | 0.054 | 58.370 | 6.945 | -36.634 | 5.971 | 0.028 | 7 |
| 64 | Leu* | 0.579 | 0.094 | 77.490 | 12.361 | -53.290 | 10.747 | 0.036 | 7 |
| 65 | Leu | 0.437 | 0.040 | 58.406 | 5.326 | -35.155 | 4.453 | 0.022 | 7 |
| 66 | Leu | 0.501 | 0.064 | 66.986 | 8.502 | -38.590 | 6.384 | 0.032 | 8 |
| 67 | Leu | 0.852 | 0.201 | 114.026 | 26.618 | -98.538 | 26.397 | 0.047 | 8 |
| 68 | Leu | 0.606 | 0.091 | 81.065 | 11.997 | -63.073 | 11.421 | 0.031 | 7 |
| 69 | Leu | 0.395 | 0.062 | 52.863 | 8.057 | -35.946 | 7.132 | 0.035 | 8 |
| 70 | Leu | 0.137 | 0.006 | 18.288 | 0.770 | -11.839 | 0.984 | 0.010 | 8 |
| 71 | Leu | 0.753 | 0.183 | 100.743 | 24.040 | -107.593 | 29.476 | 0.041 | 7 |
| 72 | Val | 0.556 | 0.043 | 74.409 | 5.570 | -38.887 | 4.011 | 0.020 | 7 |
| 73 | Val | 0.748 | 0.091 | 100.033 | 12.336 | -72.206 | 11.262 | 0.027 | 6 |
| 74 | Val | 0.846 | 0.080 | 113.219 | 10.513 | -64.911 | 7.650 | 0.024 | 7 |
| 75 | Val | 0.772 | 0.218 | 103.262 | 29.756 | -85.363 | 28.516 | 0.059 | 8 |
| 76 | Val | 0.631 | 0.078 | 84.455 | 10.485 | -67.134 | 10.108 | 0.026 | 7 |
| 77 | Val | 0.232 | 0.020 | 30.976 | 2.616 | -18.054 | 2.482 | 0.022 | 8 |
| 78 | Val | 0.596 | 0.094 | 79.743 | 12.388 | -67.422 | 12.296 | 0.032 | 8 |
| 79 | Val | 0.616 | 0.094 | 82.353 | 12.359 | -60.707 | 10.778 | 0.033 | 7 |
| 80 | Val | 0.308 | 0.034 | 41.219 | 4.521 | -25.309 | 3.989 | 0.027 | 8 |
| 81 | Val | 0.236 | 0.024 | 31.530 | 3.228 | -23.545 | 3.504 | 0.023 | 8 |
| 82 | Val | 0.787 | 0.156 | 105.242 | 20.674 | -74.488 | 17.357 | 0.045 | 8 |
| 83 | Val | 0.721 | 0.230 | 96.503 | 30.442 | -83.532 | 30.459 | 0.063 | 8 |
| 84 | Val | 0.641 | 0.067 | 85.793 | 8.744 | -64.988 | 8.100 | 0.022 | 7 |
| 85 | Val | 0.424 | 0.038 | 56.663 | 4.782 | -43.011 | 4.752 | 0.018 | 6 |
| 86 | Val | 0.789 | 0.098 | 105.602 | 13.378 | -73.803 | 11.416 | 0.029 | 7 |
| 87 | Val | 0.939 | 0.154 | 125.628 | 20.159 | -93.996 | 17.995 | 0.035 | 7 |
| 88 | Val | 0.701 | 0.130 | 93.754 | 17.421 | -98.214 | 20.655 | 0.032 | 8 |
| 89 | Val | 0.716 | 0.238 | 95.806 | 31.865 | -98.404 | 37.840 | 0.058 | 7 |
| 90 | Val | 0.917 | 0.178 | 122.718 | 23.815 | -103.902 | 23.134 | 0.039 | 8 |
| 91 | Val | 0.716 | 0.080 | 95.804 | 10.819 | -67.468 | 8.804 | 0.022 | 6 |
| 92 | Val | 0.659 | 0.149 | 88.113 | 19.418 | -79.354 | 20.207 | 0.043 | 8 |
| 93 | Val | 0.397 | 0.039 | 53.044 | 5.195 | -33.000 | 4.932 | 0.022 | 6 |
| 94 | Val | 0.864 | 0.184 | 115.545 | 24.285 | -73.767 | 19.165 | 0.050 | 7 |
| 95 | Val | 0.680 | 0.086 | 90.974 | 11.232 | -61.098 | 9.125 | 0.029 | 8 |
| 96 | Val | 0.757 | 0.118 | 101.236 | 15.924 | -60.720 | 12.648 | 0.038 | 6 |
| 97 | Val | 0.813 | 0.082 | 108.760 | 10.849 | -84.455 | 9.898 | 0.021 | 7 |
| 98 | Val | 0.164 | 0.016 | 21.973 | 2.073 | -16.604 | 2.602 | 0.021 | 8 |

**Table S8. Order parameters** of the $\alpha_{1B}$-AR-B1D1 binding **ρ-TIA**. Data was recorded at **320 K** at 700 MHz. Standard errors of order parameters ($S^2_{axis}$) were estimated using 1 000 Monte Carlo samplings based on the η and the $\tau_c$ standard errors. Amino acid assignments that are not fully unambiguous are marked with a star. Ile assignments in parentheses indicate less likely alternative assignments. Ile assignments separated by slashes are equally likely. The degree of freedom (df) indicates if data points were excluded to fit η and δ. A df of 9 means that all data points (11) were included, 8 means that one point and 7 means that two data points were excluded from the fit.

| Peak Nr. | Residue | Amino Acid | $S^2_{axis}$ | $S^2_{axis}$ SE | η [$s^{-1}$] | η SE [$s^{-1}$] | δ [$s^{-1}$] | δ SE [$s^{-1}$] | RSE | df |
|---|---|---|---|---|---|---|---|---|---|---|
| 1 | 42 | Ile | 0.194 | 0.024 | 26.018 | 3.178 | -26.333 | 4.553 | 0.025 | 9 |
| 2 | 46 | Ile | 0.462 | 0.057 | 61.742 | 7.772 | -48.851 | 8.181 | 0.030 | 8 |
| 3 | 56 | Ile | 1.053 | 0.161 | 140.916 | 21.487 | -90.905 | 17.374 | 0.038 | 8 |
| 4 | 60 | Ile | 0.904 | 0.125 | 120.911 | 16.970 | -81.891 | 14.527 | 0.035 | 8 |
| 5 | 67 | Ile | 0.662 | 0.128 | 88.570 | 16.454 | -57.224 | 13.671 | 0.051 | 9 |
| 6 | 120 | Ile | 0.345 | 0.040 | 46.178 | 5.180 | -34.267 | 5.546 | 0.028 | 8 |
| 7 | 133 (219B) | Ile | 0.993 | 0.169 | 132.805 | 22.519 | -73.429 | 16.404 | 0.047 | 8 |
| 8 | 141 | Ile | 0.727 | 0.183 | 97.240 | 24.267 | -86.386 | 25.690 | 0.053 | 9 |
| 9 | 145 | Ile | 0.427 | 0.103 | 57.129 | 13.739 | -73.989 | 21.456 | 0.040 | 8 |
| 10 | 176 | Ile | 0.413 | 0.057 | 55.197 | 7.439 | -45.230 | 7.834 | 0.033 | 9 |
| 11 | 178 | Ile | 0.209 | 0.022 | 27.981 | 2.874 | -23.919 | 3.613 | 0.024 | 9 |
| 12 | 214 | Ile | 0.434 | 0.026 | 58.110 | 3.313 | -43.607 | 3.395 | 0.014 | 8 |
| 13 | 219A | Ile | 0.842 | 0.078 | 112.630 | 10.033 | -91.432 | 9.972 | 0.020 | 8 |
| 14 | 228 | Ile | 0.477 | 0.089 | 63.858 | 11.862 | -68.073 | 15.639 | 0.036 | 8 |
| 15 | 298 | Ile | 0.387 | 0.045 | 51.812 | 5.836 | -30.801 | 5.775 | 0.030 | 7 |
| 16 | 304 | Ile | 0.615 | 0.160 | 82.321 | 20.976 | -57.012 | 18.512 | 0.067 | 9 |
| 17 | 312 | Ile | 0.431 | 0.118 | 57.600 | 15.378 | -67.217 | 23.068 | 0.047 | 7 |
| 18 | 346 | Ile | 0.383 | 0.054 | 51.232 | 7.150 | -46.983 | 8.505 | 0.031 | 8 |
| 19 | 84/139 | Ile | 0.732 | 0.158 | 97.883 | 20.749 | -72.370 | 19.497 | 0.051 | 8 |
| 20 | | Ile | 0.506 | 0.055 | 67.707 | 7.680 | -40.581 | 6.567 | 0.032 | 8 |
| 21 | | Ile | 0.772 | 0.181 | 103.237 | 23.618 | -71.214 | 20.922 | 0.058 | 8 |
| 22 | | Leu | 0.154 | 0.005 | 20.630 | 0.518 | -9.521 | 0.552 | 0.008 | 9 |
| 23 | | Leu | 0.565 | 0.064 | 75.623 | 8.793 | -26.385 | 5.410 | 0.043 | 8 |
| 24 | | Leu | 0.360 | 0.032 | 48.104 | 4.242 | -47.049 | 5.452 | 0.018 | 8 |
| 25 | | Leu | 0.644 | 0.039 | 86.098 | 5.081 | -57.407 | 4.727 | 0.015 | 7 |
| 26 | | Leu | 0.155 | 0.037 | 20.793 | 5.028 | -14.252 | 6.268 | 0.061 | 9 |
| 27 | | Leu | 0.375 | 0.050 | 50.134 | 6.681 | -24.455 | 5.070 | 0.043 | 9 |
| 28 | | Leu | 0.637 | 0.070 | 85.239 | 9.118 | -68.170 | 9.254 | 0.025 | 8 |
| 29 | | Leu | 0.455 | 0.067 | 60.932 | 8.842 | -38.434 | 7.583 | 0.041 | 9 |
| 30 | | Leu | 0.593 | 0.133 | 79.296 | 17.671 | -83.907 | 23.424 | 0.041 | 7 |
| 31 | | Leu | 0.297 | 0.035 | 39.768 | 4.654 | -29.080 | 4.806 | 0.030 | 9 |
| 32 | | Leu | 0.338 | 0.042 | 45.202 | 5.670 | -42.139 | 6.788 | 0.028 | 9 |
| 33 | | Leu | 0.189 | 0.009 | 25.270 | 1.123 | -11.729 | 1.075 | 0.014 | 9 |
| 34 | | Leu* | 0.457 | 0.055 | 61.092 | 7.185 | -47.496 | 7.171 | 0.029 | 9 |
| 35 | | Leu | 0.388 | 0.048 | 51.967 | 6.464 | -43.588 | 6.989 | 0.030 | 9 |
| 36 | | Leu | 0.655 | 0.117 | 87.601 | 16.029 | -103.091 | 22.225 | 0.031 | 8 |
| 37 | | Leu | 0.113 | 0.007 | 15.114 | 0.941 | -14.096 | 1.582 | 0.012 | 9 |
| 38 | | Leu | 0.353 | 0.031 | 47.258 | 4.045 | -18.675 | 2.848 | 0.030 | 8 |
| 39 | | Leu | 0.259 | 0.042 | 34.705 | 5.568 | -20.696 | 5.277 | 0.046 | 9 |
| 40 | | Leu | 0.241 | 0.016 | 32.255 | 2.083 | -24.724 | 2.345 | 0.016 | 9 |
| 41 | | Leu | 0.701 | 0.085 | 93.791 | 10.904 | -90.666 | 13.156 | 0.023 | 7 |
| 42 | | Leu | 0.372 | 0.016 | 49.713 | 1.912 | -45.314 | 2.217 | 0.009 | 9 |
| 43 | | Leu | 0.703 | 0.100 | 93.997 | 14.180 | -81.321 | 15.132 | 0.033 | 8 |
| 44 | | Leu | 0.277 | 0.024 | 37.005 | 3.133 | -29.565 | 3.493 | 0.021 | 9 |
| 45 | | Leu | 0.266 | 0.031 | 35.617 | 4.118 | -28.868 | 4.679 | 0.028 | 9 |
| 46 | | Leu | 0.580 | 0.115 | 77.581 | 15.037 | -81.173 | 18.620 | 0.038 | 9 |
| 47 | | Leu | 0.698 | 0.128 | 93.372 | 17.515 | -65.031 | 15.830 | 0.048 | 8 |
| 48 | | Leu | 0.395 | 0.018 | 52.787 | 2.128 | -43.377 | 2.260 | 0.010 | 9 |
| 49 | | Leu | 0.558 | 0.136 | 74.604 | 18.715 | -43.557 | 13.949 | 0.063 | 8 |
| 50 | | Leu | 0.071 | 0.002 | 9.447 | 0.215 | -5.967 | 0.427 | 0.004 | 9 |
| 51 | | Leu | 0.412 | 0.045 | 55.091 | 5.928 | -45.684 | 6.586 | 0.025 | 8 |
| 52 | | Leu | 0.314 | 0.031 | 42.054 | 4.073 | -24.373 | 3.882 | 0.027 | 8 |
| 53 | | Leu | 0.288 | 0.025 | 38.555 | 3.161 | -39.915 | 4.210 | 0.017 | 9 |
| 54 | | Leu | 0.264 | 0.025 | 35.287 | 3.176 | -16.767 | 2.526 | 0.027 | 8 |
| 55 | | Leu | 0.308 | 0.038 | 41.227 | 4.916 | -33.465 | 5.409 | 0.029 | 9 |
| 56 | | Leu | 0.246 | 0.017 | 32.885 | 2.273 | -24.766 | 2.516 | 0.017 | 9 |

| | | | | | | | | | |
|---|---|---|---|---|---|---|---|---|---|
| 57 | Leu | 0.499 | 0.020 | 66.799 | 2.472 | -27.886 | 1.715 | 0.013 | 8 |
| 58 | Leu | 0.407 | 0.049 | 54.472 | 6.492 | -39.084 | 6.210 | 0.031 | 9 |
| 59 | Leu | 0.333 | 0.056 | 44.577 | 7.327 | -40.327 | 8.594 | 0.037 | 9 |
| 60 | Leu | 0.519 | 0.103 | 69.485 | 13.803 | -49.998 | 12.785 | 0.052 | 9 |
| 61 | Leu* | 0.080 | 0.003 | 10.740 | 0.408 | -5.628 | 0.703 | 0.008 | 9 |
| 62 | Leu | 0.275 | 0.026 | 36.778 | 3.686 | -24.708 | 3.681 | 0.027 | 9 |
| 63 | Leu | 0.472 | 0.046 | 63.127 | 6.140 | -54.196 | 6.591 | 0.023 | 9 |
| 64 | Leu | 0.268 | 0.014 | 35.800 | 1.873 | -26.049 | 1.981 | 0.013 | 9 |
| 65 | Leu | 0.667 | 0.075 | 89.219 | 10.088 | -94.585 | 12.783 | 0.021 | 8 |
| 66 | Leu | 0.590 | 0.114 | 78.882 | 14.700 | -53.765 | 12.879 | 0.050 | 9 |
| 67 | Leu | 0.345 | 0.023 | 46.206 | 2.973 | -30.388 | 2.802 | 0.016 | 7 |
| 68 | Leu | 0.599 | 0.131 | 80.175 | 17.857 | -51.794 | 15.615 | 0.061 | 8 |
| 69 | Leu | 0.322 | 0.018 | 43.086 | 2.267 | -23.419 | 1.907 | 0.016 | 9 |
| 70 | Leu | 0.502 | 0.054 | 67.171 | 7.334 | -47.537 | 6.738 | 0.029 | 9 |
| 71 | Leu | 0.362 | 0.041 | 48.463 | 5.538 | -42.528 | 6.263 | 0.026 | 9 |
| 72 | Leu | 0.909 | 0.116 | 121.645 | 15.422 | -82.796 | 13.241 | 0.031 | 8 |
| 73 | Leu | 0.378 | 0.036 | 50.608 | 4.919 | -42.650 | 5.360 | 0.023 | 9 |
| 74 | Leu | 0.340 | 0.058 | 45.483 | 7.480 | -28.176 | 6.717 | 0.047 | 9 |
| 75 | Leu | 0.849 | 0.242 | 113.630 | 31.544 | -121.069 | 39.209 | 0.049 | 8 |
| 76 | Leu | 0.497 | 0.048 | 66.509 | 6.310 | -48.476 | 6.558 | 0.024 | 7 |
| 77 | Val | 0.457 | 0.057 | 61.174 | 7.384 | -55.074 | 8.570 | 0.027 | 8 |
| 78 | Val | 0.890 | 0.157 | 119.046 | 19.729 | -105.304 | 20.887 | 0.034 | 8 |
| 79 | Val | 0.614 | 0.053 | 82.101 | 6.860 | -44.967 | 5.347 | 0.025 | 8 |
| 80 | Val | 0.770 | 0.108 | 103.055 | 14.441 | -83.647 | 14.118 | 0.032 | 9 |
| 81 | Val | 0.644 | 0.087 | 86.188 | 11.158 | -59.032 | 10.069 | 0.034 | 8 |
| 82 | Val | 0.757 | 0.168 | 101.286 | 21.633 | -87.906 | 22.954 | 0.046 | 8 |
| 83 | Val | 0.957 | 0.256 | 127.997 | 35.029 | -112.001 | 36.473 | 0.055 | 8 |
| 84 | Val | 0.564 | 0.059 | 75.476 | 7.931 | -80.011 | 10.220 | 0.020 | 8 |
| 85 | Val | 0.622 | 0.024 | 83.255 | 2.614 | -56.134 | 2.257 | 0.008 | 9 |
| 86 | Val | 0.585 | 0.063 | 78.269 | 8.415 | -45.321 | 6.558 | 0.032 | 9 |
| 87 | Val | 0.561 | 0.074 | 75.067 | 9.891 | -50.420 | 8.978 | 0.035 | 8 |
| 88 | Val | 0.865 | 0.068 | 115.661 | 8.913 | -64.442 | 6.665 | 0.022 | 8 |
| 89 | Val | 0.641 | 0.072 | 85.818 | 9.291 | -50.630 | 7.252 | 0.031 | 9 |
| 90 | Val | 0.555 | 0.150 | 74.289 | 19.716 | -74.550 | 23.654 | 0.054 | 9 |
| 91 | Val | 0.845 | 0.238 | 113.080 | 33.090 | -89.551 | 31.430 | 0.066 | 9 |
| 92 | Val | 0.641 | 0.138 | 85.686 | 18.926 | -56.022 | 15.924 | 0.060 | 9 |
| 93 | Val | 0.636 | 0.107 | 85.041 | 13.976 | -83.679 | 16.305 | 0.033 | 9 |
| 94 | Val | 0.734 | 0.167 | 98.136 | 23.165 | -88.457 | 24.802 | 0.050 | 9 |
| 95 | Val | 0.702 | 0.058 | 93.897 | 7.483 | -50.991 | 5.661 | 0.024 | 8 |
| 96 | Val | 0.566 | 0.052 | 75.752 | 6.899 | -74.031 | 8.648 | 0.018 | 7 |
| 97 | Val | 0.971 | 0.195 | 129.866 | 26.652 | -108.508 | 26.115 | 0.043 | 9 |
| 98 | Val | 0.450 | 0.024 | 60.192 | 2.984 | -46.912 | 2.990 | 0.012 | 9 |
| 99 | Val | 0.130 | 0.015 | 17.386 | 1.949 | -14.281 | 2.881 | 0.025 | 9 |
| 100 | Val | 0.863 | 0.074 | 115.503 | 9.757 | -82.987 | 8.788 | 0.020 | 8 |
| 101 | Val | 0.743 | 0.252 | 99.334 | 34.903 | -105.232 | 43.737 | 0.064 | 8 |
| 102 | Val | 1.078 | 0.265 | 144.181 | 34.715 | -91.964 | 27.744 | 0.060 | 8 |
| 103 | Val | 0.224 | 0.024 | 29.989 | 3.002 | -15.605 | 3.777 | 0.023 | 7 |

**Table S9. Order parameters** of the $\alpha_{1B}$-AR-B1D1 binding **tamsulosin**. Data was recorded at **320 K** at 700 MHz. Standard errors of order parameters ($S^2_{axis}$) were estimated using 1 000 Monte Carlo samplings based on the η and the $\tau_c$ standard errors. Amino acid assignments that are not fully unambiguous are marked with a star. Ile assignments in parentheses indicate less likely alternative assignments. The degree of freedom (df) indicates if data points were excluded to fit η and δ. A df of 9 means that all data points (11) were included, 8 means that one point and 7 means that two data points were excluded from the fit.

| Peak Nr. | Residue | Amino Acid | $S^2_{axis}$ | $S^2_{axis}$ SE | η [$s^{-1}$] | η SE [$s^{-1}$] | δ [$s^{-1}$] | δ SE [$s^{-1}$] | RSE | df |
|---|---|---|---|---|---|---|---|---|---|---|
| 1 | 42 | Ile | 0.205 | 0.015 | 27.382 | 1.866 | -22.505 | 2.305 | 0.016 | 9 |
| 2 | 46 | Ile | 0.558 | 0.119 | 74.697 | 15.610 | -58.199 | 15.242 | 0.051 | 9 |
| 3 | 56 | Ile | 0.842 | 0.146 | 112.614 | 19.203 | -54.659 | 13.154 | 0.052 | 8 |
| 4 | 60A | Ile | 0.835 | 0.136 | 111.659 | 18.527 | -77.281 | 15.823 | 0.041 | 9 |
| 5 | 60B | Ile | 0.897 | 0.105 | 119.972 | 14.336 | -73.357 | 11.390 | 0.032 | 8 |
| 6 | 67 | Ile | 0.589 | 0.066 | 78.770 | 8.598 | -54.307 | 7.894 | 0.029 | 8 |
| 7 | I133A (219D) | Ile | 0.756 | 0.175 | 101.127 | 24.126 | -79.112 | 22.936 | 0.056 | 9 |
| 8 | 141 | Ile | 0.683 | 0.175 | 91.402 | 23.522 | -67.631 | 22.305 | 0.063 | 8 |
| 9 | 145 | Ile | 0.513 | 0.119 | 68.677 | 16.118 | -81.339 | 22.999 | 0.041 | 8 |
| 10 | 163 | Ile | 0.332 | 0.094 | 44.417 | 12.448 | -45.460 | 16.060 | 0.058 | 9 |
| 11 | 176 | Ile | 0.687 | 0.078 | 91.957 | 10.893 | -62.958 | 9.742 | 0.031 | 8 |
| 12 | 178A | Ile | 0.143 | 0.032 | 19.165 | 4.198 | -15.154 | 5.821 | 0.051 | 9 |
| 13 | 178B | Ile | 0.244 | 0.044 | 32.638 | 6.093 | -29.093 | 7.754 | 0.042 | 8 |
| 14 | 214 | Ile | 0.573 | 0.067 | 76.710 | 9.094 | -60.339 | 8.921 | 0.029 | 9 |
| 15 | 219B (133B) | Ile | 0.842 | 0.213 | 112.658 | 28.474 | -97.289 | 29.740 | 0.053 | 8 |
| 16 | 219C (133C) | Ile | 0.994 | 0.199 | 132.910 | 26.853 | -104.138 | 25.436 | 0.044 | 8 |
| 17 | 228 | Ile | 0.501 | 0.067 | 66.956 | 8.642 | -64.807 | 10.156 | 0.027 | 9 |
| 18 | 298 | Ile | 0.575 | 0.176 | 76.944 | 22.688 | -79.600 | 28.596 | 0.057 | 8 |
| 19 | 304A | Ile | 0.695 | 0.212 | 92.926 | 28.159 | -92.873 | 33.833 | 0.059 | 8 |
| 20 | 312 | Ile | 0.502 | 0.054 | 67.113 | 7.221 | -45.513 | 6.720 | 0.029 | 8 |
| 21 | 346A | Ile | 0.392 | 0.051 | 52.428 | 6.815 | -37.593 | 6.559 | 0.034 | 9 |
| 22 | 346B | Ile | 0.487 | 0.089 | 65.127 | 12.044 | -61.628 | 14.322 | 0.040 | 8 |
| 23 | | Ile | 0.556 | 0.115 | 74.355 | 15.310 | -46.219 | 12.629 | 0.059 | 9 |
| 24 | | Leu | 0.387 | 0.044 | 51.741 | 5.784 | -28.001 | 4.632 | 0.034 | 9 |
| 25 | | Leu | 0.563 | 0.076 | 75.340 | 9.359 | -46.536 | 7.535 | 0.035 | 8 |
| 26 | | Leu | 0.344 | 0.030 | 46.004 | 4.023 | -34.256 | 4.074 | 0.022 | 9 |
| 27 | | Leu | 0.397 | 0.051 | 53.062 | 6.649 | -30.435 | 5.850 | 0.036 | 8 |
| 28 | | Leu | 0.424 | 0.036 | 56.748 | 4.715 | -21.497 | 3.024 | 0.030 | 9 |
| 29 | | Leu | 0.367 | 0.041 | 49.037 | 5.492 | -42.227 | 6.112 | 0.026 | 9 |
| 30 | | Leu | 0.402 | 0.049 | 53.783 | 6.604 | -42.556 | 6.800 | 0.030 | 9 |
| 31 | | Leu | 0.804 | 0.113 | 107.535 | 14.829 | -73.692 | 12.617 | 0.035 | 9 |
| 32 | | Leu | 0.324 | 0.015 | 43.360 | 1.917 | -36.576 | 2.464 | 0.010 | 7 |
| 33 | | Leu | 0.436 | 0.029 | 58.363 | 3.802 | -60.207 | 4.730 | 0.011 | 7 |
| 34 | | Leu | 0.494 | 0.108 | 66.149 | 14.367 | -53.494 | 14.636 | 0.053 | 9 |
| 35 | | Leu | 0.199 | 0.007 | 26.598 | 0.813 | -19.487 | 0.949 | 0.008 | 9 |
| 36 | | Leu | 0.351 | 0.033 | 47.009 | 4.209 | -34.119 | 4.422 | 0.023 | 8 |
| 37 | | Leu | 0.430 | 0.095 | 57.534 | 12.536 | -35.219 | 10.637 | 0.063 | 9 |
| 38 | | Leu | 0.412 | 0.072 | 55.054 | 9.524 | -69.715 | 14.119 | 0.029 | 8 |
| 39 | | Leu | 0.255 | 0.046 | 34.095 | 5.986 | -22.366 | 6.027 | 0.048 | 9 |
| 40 | | Leu | 0.295 | 0.035 | 39.412 | 4.695 | -29.809 | 4.970 | 0.030 | 9 |
| 41 | | Leu | 0.201 | 0.010 | 26.856 | 1.250 | -9.252 | 1.042 | 0.016 | 9 |
| 42 | | Leu | 0.535 | 0.058 | 71.589 | 7.464 | -49.723 | 6.988 | 0.027 | 8 |
| 43 | | Leu | 0.362 | 0.081 | 48.425 | 10.864 | -28.525 | 9.312 | 0.066 | 9 |
| 44 | | Leu | 0.497 | 0.088 | 66.500 | 11.927 | -41.327 | 10.479 | 0.050 | 8 |
| 45 | | Leu | 0.398 | 0.063 | 53.196 | 8.186 | -38.545 | 7.918 | 0.040 | 9 |
| 46 | | Leu | 0.594 | 0.159 | 79.457 | 22.010 | -69.627 | 23.462 | 0.062 | 9 |
| 47 | | Leu | 0.474 | 0.087 | 63.437 | 11.324 | -57.041 | 12.088 | 0.035 | 8 |
| 48 | | Leu | 0.449 | 0.058 | 60.037 | 7.612 | -45.716 | 7.497 | 0.032 | 9 |
| 49 | | Leu | 0.767 | 0.091 | 102.626 | 12.987 | -71.330 | 11.227 | 0.032 | 9 |
| 50 | | Leu | 0.347 | 0.051 | 46.472 | 6.876 | -41.955 | 8.173 | 0.033 | 8 |
| 51 | | Leu | 0.359 | 0.036 | 48.041 | 4.802 | -47.783 | 5.990 | 0.021 | 9 |
| 52 | | Leu | 0.565 | 0.101 | 75.536 | 13.204 | -57.866 | 13.173 | 0.043 | 8 |
| 53 | | Leu | 0.381 | 0.034 | 50.986 | 4.223 | -32.970 | 3.804 | 0.023 | 9 |
| 54 | | Leu* | 0.558 | 0.075 | 74.585 | 9.998 | -50.543 | 9.141 | 0.036 | 8 |
| 55 | | Leu | 0.575 | 0.094 | 76.960 | 12.908 | -86.908 | 17.497 | 0.030 | 8 |
| 56 | | Leu | 0.354 | 0.031 | 47.381 | 3.964 | -27.382 | 3.375 | 0.025 | 9 |
| 57 | | Leu | 0.199 | 0.015 | 26.611 | 1.849 | -23.165 | 2.388 | 0.016 | 9 |

| | | | | | | | | | |
|---|---|---|---|---|---|---|---|---|---|
| 58 | Leu | 0.652 | 0.078 | 87.256 | 10.326 | -47.914 | 7.648 | 0.036 | 9 |
| 59 | Leu | 0.263 | 0.037 | 35.131 | 4.980 | -31.953 | 6.145 | 0.032 | 9 |
| 60 | Leu | 0.481 | 0.099 | 64.385 | 13.190 | -60.765 | 15.242 | 0.044 | 9 |
| 61 | Leu | 0.460 | 0.066 | 61.479 | 8.395 | -54.172 | 9.226 | 0.031 | 9 |
| 62 | Leu | 0.681 | 0.108 | 91.145 | 14.664 | -70.891 | 14.016 | 0.039 | 9 |
| 63 | Leu | 0.480 | 0.058 | 64.252 | 8.033 | -40.718 | 7.194 | 0.035 | 8 |
| 64 | Leu | 0.587 | 0.081 | 78.512 | 10.714 | -52.748 | 9.291 | 0.037 | 9 |
| 65 | Leu | 0.179 | 0.015 | 23.966 | 1.973 | -17.076 | 2.358 | 0.021 | 9 |
| 66 | Leu | 0.648 | 0.056 | 86.631 | 6.892 | -58.490 | 5.828 | 0.018 | 7 |
| 67 | Leu | 0.764 | 0.170 | 102.232 | 22.591 | -67.722 | 19.446 | 0.057 | 8 |
| 68 | Leu | 0.800 | 0.110 | 106.972 | 14.235 | -96.370 | 15.963 | 0.027 | 7 |
| 69 | Leu | 0.458 | 0.095 | 61.321 | 12.547 | -29.697 | 9.638 | 0.065 | 8 |
| 70 | Leu | 0.086 | 0.006 | 11.508 | 0.770 | -7.104 | 1.312 | 0.014 | 9 |
| 71 | Leu | 0.447 | 0.064 | 59.791 | 8.650 | -45.627 | 8.537 | 0.037 | 9 |
| 72 | Leu | 0.169 | 0.008 | 22.604 | 1.058 | -6.106 | 0.913 | 0.017 | 9 |
| 73 | Leu | 0.241 | 0.021 | 32.187 | 2.776 | -29.825 | 3.542 | 0.019 | 9 |
| 74 | Leu | 0.564 | 0.128 | 75.422 | 17.364 | -69.127 | 20.742 | 0.048 | 7 |
| 75 | Leu | 0.354 | 0.042 | 47.327 | 5.519 | -36.440 | 5.692 | 0.029 | 9 |
| 76 | Leu | 0.103 | 0.005 | 13.771 | 0.580 | -7.666 | 0.846 | 0.010 | 9 |
| 77 | Leu | 0.210 | 0.017 | 28.113 | 2.196 | -22.855 | 2.672 | 0.019 | 9 |
| 78 | Leu | 0.753 | 0.210 | 100.722 | 28.575 | -96.509 | 32.884 | 0.056 | 8 |
| 79 | Leu | 0.174 | 0.018 | 23.316 | 2.376 | -14.502 | 2.685 | 0.028 | 9 |
| 80 | Leu | 0.342 | 0.036 | 45.809 | 4.806 | -31.034 | 4.567 | 0.028 | 9 |
| 81 | Leu | 0.521 | 0.128 | 69.707 | 17.373 | -77.000 | 24.253 | 0.045 | 7 |
| 82 | Leu | 0.148 | 0.013 | 19.838 | 1.677 | -17.089 | 2.395 | 0.019 | 9 |
| 83 | Leu | 0.310 | 0.037 | 41.407 | 4.842 | -27.763 | 5.003 | 0.031 | 8 |
| 84 | Leu | 0.476 | 0.063 | 63.699 | 8.411 | -38.851 | 7.362 | 0.037 | 8 |
| 85 | Leu | 0.143 | 0.015 | 19.079 | 1.973 | -10.728 | 2.360 | 0.028 | 9 |
| 86 | Leu | 0.390 | 0.035 | 52.236 | 4.598 | -42.666 | 4.870 | 0.021 | 9 |
| 87 | Leu | 0.264 | 0.027 | 35.302 | 3.490 | -23.378 | 3.770 | 0.026 | 8 |
| 88 | Leu | 0.444 | 0.074 | 59.443 | 9.813 | -68.619 | 13.922 | 0.030 | 8 |
| 89 | Leu | 0.427 | 0.040 | 57.142 | 5.338 | -22.357 | 3.474 | 0.033 | 9 |
| 90 | Leu | 0.328 | 0.034 | 43.828 | 4.349 | -21.375 | 3.427 | 0.032 | 9 |
| 91 | Leu | 0.593 | 0.103 | 79.351 | 13.612 | -60.744 | 13.027 | 0.042 | 9 |
| 92 | Leu/Val | 0.495 | 0.039 | 66.161 | 5.000 | -42.186 | 4.473 | 0.021 | 8 |
| 93 | Val | 0.864 | 0.224 | 115.563 | 29.681 | -87.697 | 27.887 | 0.059 | 8 |
| 94 | Val | 0.952 | 0.174 | 127.411 | 22.415 | -81.918 | 18.325 | 0.044 | 8 |
| 95 | Val | 0.633 | 0.137 | 84.724 | 17.456 | -97.211 | 23.762 | 0.036 | 8 |
| 96 | Val | 0.635 | 0.090 | 84.889 | 11.817 | -49.118 | 9.498 | 0.040 | 8 |
| 97 | Val | 0.976 | 0.132 | 130.576 | 17.631 | -77.428 | 13.535 | 0.036 | 8 |
| 98 | Val | 0.643 | 0.112 | 85.976 | 15.063 | -54.833 | 13.607 | 0.047 | 7 |
| 99 | Val | 0.585 | 0.125 | 78.220 | 15.982 | -54.205 | 14.743 | 0.053 | 8 |
| 100 | Val | 0.149 | 0.018 | 19.879 | 2.352 | -15.663 | 3.203 | 0.028 | 9 |
| 101 | Val | 0.386 | 0.125 | 51.700 | 16.582 | -76.159 | 29.305 | 0.048 | 8 |
| 102 | Val | 0.729 | 0.068 | 97.587 | 9.389 | -62.478 | 7.665 | 0.026 | 9 |
| 103 | Val | 0.705 | 0.130 | 94.302 | 17.715 | -90.407 | 20.043 | 0.038 | 9 |
| 104 | Val | 0.686 | 0.103 | 91.775 | 13.666 | -78.626 | 14.115 | 0.033 | 9 |
| 105 | Val | 0.534 | 0.041 | 71.484 | 5.315 | -42.997 | 4.306 | 0.022 | 9 |
| 106 | Val | 0.700 | 0.165 | 93.615 | 22.414 | -86.235 | 24.538 | 0.050 | 9 |
| 107 | Val | 0.893 | 0.168 | 119.480 | 21.990 | -87.524 | 19.505 | 0.044 | 9 |
| 108 | Val | 0.531 | 0.043 | 71.057 | 5.664 | -44.179 | 4.701 | 0.023 | 9 |
| 109 | Val | 0.298 | 0.028 | 39.892 | 3.599 | -27.098 | 3.540 | 0.024 | 9 |
| 110 | Val | 0.477 | 0.033 | 63.756 | 4.268 | -46.688 | 4.471 | 0.017 | 7 |
| 111 | Val | 0.785 | 0.163 | 104.978 | 21.396 | -75.690 | 19.554 | 0.050 | 8 |
| 112 | Val | 0.883 | 0.236 | 118.159 | 31.005 | -87.334 | 28.461 | 0.061 | 8 |
| 113 | Val | 0.229 | 0.029 | 30.605 | 3.702 | -29.871 | 4.957 | 0.026 | 9 |
| 114 | Val | 0.967 | 0.153 | 129.329 | 20.640 | -81.403 | 15.568 | 0.039 | 8 |
| 115 | Val | 0.624 | 0.116 | 83.413 | 15.770 | -50.333 | 13.076 | 0.053 | 8 |
| 116 | Val | 0.614 | 0.067 | 82.165 | 9.026 | -79.981 | 10.466 | 0.022 | 9 |
| 117 | Val | 0.288 | 0.026 | 38.538 | 3.427 | -25.227 | 3.324 | 0.024 | 9 |
| 118 | Val | 0.799 | 0.232 | 106.823 | 30.715 | -118.381 | 38.139 | 0.048 | 8 |
| 119 | Val | 0.867 | 0.137 | 116.048 | 17.700 | -89.424 | 16.823 | 0.035 | 8 |
| 120 | Val | 0.503 | 0.062 | 67.228 | 8.225 | -68.806 | 10.429 | 0.024 | 8 |
| 121 | Val | 0.685 | 0.097 | 91.580 | 13.075 | -52.618 | 10.339 | 0.041 | 8 |
| 122 | Val | 0.747 | 0.110 | 99.931 | 14.345 | -110.171 | 18.211 | 0.025 | 9 |
| 123 | Val | 1.047 | 0.161 | 140.037 | 21.574 | -92.958 | 16.939 | 0.036 | 8 |
| 124 | Val | 0.731 | 0.208 | 97.779 | 26.961 | -98.884 | 32.545 | 0.052 | 8 |
| 125 | Val | 0.352 | 0.035 | 47.035 | 4.546 | -37.263 | 4.791 | 0.024 | 9 |

**Table S10. Relative side chain dynamics** of the $\alpha_{1B}$-AR-B1D1 binding **prazosin**. Data was recorded at **298 K** at 700 MHz. Assignments and ambiguities are mentioned under "Remark". Ile assignments in parentheses indicate less likely alternative assignments. Ile assignments separated by slashes are equally likely. The degree of freedom (df) indicates if data points were excluded to fit η and δ. A df of 5 means that all data points (7) were included, 4 means that one point and 3 means that two data points were excluded from the fit.

| Peak Nr. | Amino Acid | Remark | η [$s^{-1}$] | η SE [$s^{-1}$] | δ [$s^{-1}$] | δ SE [$s^{-1}$] | RSE | df |
|---|---|---|---|---|---|---|---|---|
| 1 | Ile | I304/I133A | 198.337 | 8.145 | -137.538 | 6.713 | 0.008 | 5 |
| 2 | Ile | I42 | 61.867 | 8.297 | -68.467 | 11.231 | 0.021 | 5 |
| 3 | Ile | I219A | 214.473 | 69.531 | -203.960 | 78.739 | 0.044 | 3 |
| 4 | Ile | I145 | 147.292 | 26.648 | -135.124 | 28.411 | 0.030 | 5 |
| 5 | Ile | I178B | 64.673 | 17.049 | -50.180 | 17.502 | 0.053 | 5 |
| 6 | Ile | I163 | 84.334 | 18.759 | -53.474 | 15.826 | 0.051 | 5 |
| 7 | Ile | | 226.436 | 6.476 | -229.606 | 7.584 | 0.004 | 3 |
| 8 | Ile | I219C (I133B) | 214.932 | 56.209 | -164.423 | 53.567 | 0.044 | 3 |
| 9 | Ile | | 247.953 | 69.380 | -227.834 | 72.681 | 0.037 | 4 |
| 10 | Ile | I46 | 149.398 | 43.472 | -166.538 | 54.876 | 0.040 | 5 |
| 11 | Ile | | 191.556 | 66.788 | -158.641 | 65.865 | 0.058 | 4 |
| 12 | Ile | I178A | 66.100 | 15.130 | -50.142 | 15.212 | 0.047 | 5 |
| 13 | Ile | I67 | 219.413 | 66.343 | -187.404 | 64.538 | 0.047 | 5 |
| 14 | Ile | I56 | 251.983 | 54.411 | -200.291 | 49.174 | 0.034 | 5 |
| 15 | Ile | I312 | 126.947 | 7.258 | -98.606 | 6.811 | 0.011 | 5 |
| 16 | Ile | | 153.053 | 15.735 | -100.917 | 12.746 | 0.022 | 5 |
| 17 | Ile | | 144.429 | 25.661 | -126.976 | 28.798 | 0.030 | 3 |
| 18 | Ile | | 184.822 | 54.660 | -176.717 | 59.415 | 0.044 | 5 |
| 19 | Ile | I214 | 146.281 | 14.063 | -124.525 | 14.078 | 0.017 | 5 |
| 20 | Leu | | 204.413 | 46.904 | -182.136 | 48.822 | 0.035 | 4 |
| 21 | Leu | | 166.494 | 59.312 | -160.831 | 65.542 | 0.055 | 5 |
| 22 | Leu | | 95.697 | 2.395 | -80.326 | 2.456 | 0.005 | 5 |
| 23 | Leu | Val? | 196.154 | 60.747 | -174.845 | 61.859 | 0.048 | 5 |
| 24 | Leu | | 166.077 | 27.192 | -116.288 | 22.940 | 0.033 | 5 |
| 25 | Leu | | 110.639 | 10.666 | -82.076 | 9.767 | 0.020 | 5 |
| 26 | Leu | | 67.165 | 5.890 | -48.203 | 5.676 | 0.018 | 5 |
| 27 | Leu | | 113.182 | 7.175 | -76.182 | 6.078 | 0.014 | 5 |
| 28 | Leu | | 112.119 | 17.472 | -49.274 | 13.151 | 0.040 | 3 |
| 29 | Leu | | 143.443 | 22.815 | -96.316 | 18.856 | 0.034 | 5 |
| 30 | Leu | | 123.211 | 13.043 | -79.968 | 10.651 | 0.023 | 5 |
| 31 | Leu | | 111.604 | 22.336 | -105.737 | 24.977 | 0.034 | 5 |
| 32 | Leu | | 195.468 | 12.517 | -148.039 | 11.106 | 0.012 | 5 |
| 33 | Leu | | 108.966 | 14.074 | -75.916 | 12.291 | 0.028 | 5 |
| 34 | Leu | | 121.473 | 29.473 | -92.421 | 27.301 | 0.048 | 5 |
| 35 | Leu | | 130.604 | 5.506 | -55.341 | 3.314 | 0.012 | 5 |
| 36 | Leu | | 64.919 | 3.807 | -38.387 | 3.262 | 0.014 | 5 |
| 37 | Leu | | 119.298 | 8.168 | -114.290 | 9.174 | 0.011 | 5 |
| 38 | Leu | | 72.933 | 8.985 | -71.817 | 10.818 | 0.021 | 5 |
| 39 | Leu | | 105.059 | 19.154 | -97.106 | 21.078 | 0.032 | 5 |
| 40 | Leu | | 143.977 | 23.508 | -83.099 | 17.300 | 0.038 | 5 |
| 41 | Leu | | 123.795 | 23.805 | -71.876 | 17.878 | 0.045 | 5 |
| 42 | Leu | | 96.648 | 14.132 | -68.447 | 12.671 | 0.031 | 5 |
| 43 | Leu | | 180.465 | 46.470 | -139.699 | 42.267 | 0.047 | 5 |
| 44 | Leu | | 146.374 | 23.153 | -115.703 | 21.789 | 0.030 | 5 |
| 45 | Leu | | 205.919 | 31.397 | -98.873 | 21.802 | 0.035 | 3 |
| 46 | Leu | | 93.003 | 22.220 | -92.891 | 26.368 | 0.040 | 5 |
| 47 | Leu | | 131.767 | 56.603 | -171.624 | 82.774 | 0.053 | 5 |
| 48 | Leu | | 175.023 | 16.644 | -156.244 | 17.118 | 0.015 | 5 |
| 49 | Leu | | 87.913 | 4.213 | -69.296 | 4.151 | 0.010 | 5 |
| 50 | Leu | | 67.285 | 13.797 | -47.094 | 13.064 | 0.044 | 5 |
| 51 | Leu | | 149.452 | 32.400 | -119.849 | 30.805 | 0.040 | 5 |
| 52 | Leu | | 162.687 | 20.448 | -127.817 | 18.987 | 0.023 | 5 |
| 53 | Leu | | 92.956 | 5.198 | -72.796 | 5.060 | 0.011 | 5 |
| 54 | Leu | | 39.608 | 1.730 | -28.106 | 2.004 | 0.009 | 5 |
| 55 | Leu | | 55.464 | 14.930 | -58.978 | 19.965 | 0.043 | 5 |
| 56 | Leu | | 171.460 | 58.478 | -147.366 | 58.317 | 0.058 | 5 |
| 57 | Leu | | 139.729 | 13.013 | -111.146 | 12.355 | 0.018 | 5 |
| 58 | Leu | | 37.150 | 1.341 | -21.864 | 1.476 | 0.008 | 5 |
| 59 | Leu | | 131.123 | 25.005 | -104.583 | 23.914 | 0.036 | 5 |

| | | | | | | | | |
|---|---|---|---|---|---|---|---|---|
| 60 | Leu | | 127.390 | 32.899 | -124.845 | 37.503 | 0.042 | 5 |
| 61 | Leu | | 65.677 | 3.308 | -36.651 | 2.726 | 0.012 | 5 |
| 62 | Leu | | 149.254 | 31.834 | -122.568 | 33.779 | 0.037 | 3 |
| 63 | Leu | | 106.853 | 41.247 | -86.165 | 42.421 | 0.074 | 4 |
| 64 | Leu | | 72.423 | 6.872 | -39.144 | 5.397 | 0.023 | 5 |
| 65 | Leu | | 50.365 | 2.388 | -52.231 | 3.201 | 0.008 | 5 |
| 66 | Leu | | 94.240 | 12.136 | -58.618 | 11.740 | 0.028 | 3 |
| 67 | Leu | | 196.206 | 63.481 | -126.596 | 49.421 | 0.066 | 5 |
| 68 | Leu | | 170.450 | 49.901 | -143.131 | 52.824 | 0.049 | 3 |
| 69 | Leu | | 201.563 | 60.077 | -172.217 | 58.865 | 0.048 | 5 |
| 70 | Leu | | 41.971 | 10.511 | -31.663 | 12.233 | 0.048 | 5 |
| 71 | Val | | 194.232 | 20.719 | -109.904 | 14.580 | 0.024 | 5 |
| 72 | Val | | 186.075 | 59.240 | -227.663 | 84.198 | 0.035 | 3 |
| 73 | Val | | 203.522 | 31.592 | -147.641 | 26.954 | 0.029 | 5 |
| 74 | Val | | 249.271 | 29.770 | -158.449 | 23.064 | 0.022 | 4 |
| 75 | Val | | 189.665 | 54.166 | -177.746 | 61.601 | 0.042 | 3 |
| 76 | Val | | 191.501 | 30.422 | -126.800 | 24.232 | 0.032 | 5 |
| 77 | Val | | 77.670 | 7.308 | -72.933 | 8.400 | 0.017 | 5 |
| 78 | Val | | 231.519 | 44.796 | -147.470 | 33.967 | 0.038 | 5 |
| 79 | Val | | 38.345 | 6.115 | -27.992 | 7.303 | 0.031 | 5 |
| 80 | Val | | 172.522 | 13.009 | -152.823 | 13.306 | 0.012 | 5 |
| 81 | Val | | 194.864 | 31.996 | -158.601 | 30.163 | 0.028 | 5 |
| 82 | Val | | 173.213 | 18.424 | -247.417 | 30.145 | 0.010 | 3 |
| 83 | Val | Leu? | 175.725 | 37.147 | -117.486 | 30.051 | 0.043 | 5 |
| 84 | Val | | 150.237 | 48.888 | -99.930 | 39.941 | 0.069 | 5 |
| 85 | Val | | 201.741 | 41.813 | -125.108 | 31.464 | 0.043 | 5 |
| 86 | Val | | 198.433 | 34.262 | -182.523 | 35.819 | 0.026 | 5 |
| 87 | Val | | 231.587 | 65.476 | -274.250 | 84.669 | 0.030 | 5 |
| 88 | Val | | 230.220 | 74.951 | -125.639 | 52.343 | 0.070 | 4 |
| 89 | Val | | 280.712 | 69.402 | -199.914 | 56.558 | 0.040 | 5 |
| 90 | Val | Leu? | 192.281 | 41.176 | -187.423 | 46.509 | 0.031 | 4 |
| 91 | Val | | 260.939 | 18.689 | -145.941 | 13.704 | 0.014 | 3 |
| 92 | Val | | 155.878 | 25.262 | -94.238 | 19.103 | 0.036 | 5 |
| 93 | Val | | 241.088 | 46.193 | -198.878 | 43.266 | 0.029 | 5 |
| 94 | Val | | 229.856 | 34.697 | -240.632 | 42.159 | 0.018 | 3 |
| 95 | Val | | 215.080 | 46.775 | -204.467 | 52.935 | 0.030 | 3 |
| 96 | Val | | 75.826 | 2.947 | -60.647 | 3.000 | 0.008 | 5 |

**Table S11. Relative side chain dynamics** of the **apo** $\alpha_{1B}$-AR-B1D1. Data was recorded at **298 K** at 700 MHz. Assignments and ambiguities are mentioned under "Remark". Ile assignments separated by slashes are equally likely. The degree of freedom (df) indicates if data points were excluded to fit η and δ. A df of 5 means that all data points (7) were included, 4 means that one point and 3 means that two data points were excluded from the fit.

| Peak Nr. | Amino Acid | Remark | η [s$^{-1}$] | η SE [s$^{-1}$] | δ [s$^{-1}$] | δ SE [s$^{-1}$] | RSE | df |
|---|---|---|---|---|---|---|---|---|
| 1 | Ile | | 125.793 | 28.891 | -90.450 | 25.502 | 0.047 | 5 |
| 2 | Ile | I312 | 142.067 | 29.849 | -109.681 | 27.616 | 0.040 | 5 |
| 3 | Ile | I176/I214 | 140.129 | 30.556 | -119.718 | 31.896 | 0.038 | 4 |
| 4 | Ile | I178 | 47.013 | 10.081 | -24.039 | 9.122 | 0.051 | 5 |
| 5 | Ile | I67 | 196.898 | 34.245 | -155.040 | 31.369 | 0.030 | 5 |
| 6 | Ile | I42 | 51.972 | 9.049 | -45.471 | 10.676 | 0.032 | 5 |
| 7 | Ile | | 131.647 | 27.572 | -62.797 | 19.053 | 0.054 | 4 |
| 8 | Ile | | 239.141 | 35.200 | -207.838 | 36.513 | 0.021 | 3 |
| 9 | Ile | I145 | 141.268 | 38.101 | -135.437 | 42.314 | 0.044 | 5 |
| 10 | Ile | I56 | 270.742 | 65.089 | -254.524 | 67.634 | 0.030 | 5 |
| 11 | Leu | | 80.791 | 20.057 | -82.331 | 24.535 | 0.041 | 5 |
| 12 | Leu | | 160.927 | 29.751 | -96.535 | 22.296 | 0.041 | 5 |
| 13 | Leu | | 156.588 | 24.820 | -129.350 | 24.102 | 0.028 | 5 |
| 14 | Leu | | 54.506 | 3.794 | -31.831 | 3.430 | 0.016 | 5 |
| 15 | Leu | | 182.790 | 58.420 | -167.049 | 61.111 | 0.050 | 5 |
| 16 | Leu | | 88.941 | 21.104 | -83.295 | 23.837 | 0.042 | 5 |
| 17 | Leu | | 133.153 | 47.393 | -117.769 | 49.343 | 0.062 | 5 |
| 18 | Leu | | 31.753 | 4.159 | -13.651 | 4.579 | 0.029 | 5 |
| 19 | Leu | | 121.552 | 18.054 | -52.400 | 11.917 | 0.040 | 4 |
| 20 | Leu | | 82.861 | 23.420 | -66.093 | 23.471 | 0.056 | 5 |
| 21 | Leu | | 149.250 | 48.735 | -104.216 | 41.402 | 0.067 | 5 |
| 22 | Leu | | 124.423 | 24.043 | -92.787 | 21.870 | 0.039 | 5 |
| 23 | Leu | | 115.094 | 12.095 | -96.684 | 12.201 | 0.020 | 5 |
| 24 | Leu | | 110.152 | 24.713 | -84.612 | 23.275 | 0.045 | 5 |
| 25 | Leu | | 74.229 | 9.196 | -30.711 | 6.667 | 0.032 | 3 |
| 26 | Leu | | 127.725 | 18.388 | -88.742 | 15.775 | 0.030 | 5 |
| 27 | Leu | Val? | 187.734 | 37.412 | -146.746 | 35.278 | 0.035 | 4 |
| 28 | Leu | | 59.879 | 2.662 | -35.734 | 2.353 | 0.010 | 5 |
| 29 | Leu | | 112.416 | 10.279 | -76.319 | 8.770 | 0.020 | 5 |
| 30 | Leu | | 146.041 | 35.884 | -116.856 | 34.117 | 0.046 | 5 |
| 31 | Leu | | 78.603 | 7.756 | -50.859 | 6.725 | 0.022 | 5 |
| 32 | Leu | | 212.906 | 66.943 | -134.941 | 55.586 | 0.061 | 3 |
| 33 | Leu | | 103.174 | 6.640 | -99.550 | 8.399 | 0.010 | 3 |
| 34 | Leu | | 65.413 | 4.331 | -35.520 | 3.516 | 0.016 | 5 |
| 35 | Leu | | 39.014 | 0.967 | -27.219 | 1.119 | 0.005 | 5 |
| 36 | Leu | | 192.018 | 46.085 | -151.457 | 45.710 | 0.041 | 3 |
| 37 | Leu | | 97.390 | 15.720 | -73.836 | 15.644 | 0.032 | 4 |
| 38 | Leu | | 77.330 | 8.165 | -52.117 | 7.308 | 0.023 | 5 |
| 39 | Leu | | 55.403 | 16.111 | -44.750 | 17.685 | 0.056 | 5 |
| 40 | Leu | | 99.053 | 3.980 | -60.981 | 3.206 | 0.009 | 5 |
| 41 | Leu | | 166.099 | 43.452 | -153.285 | 49.558 | 0.041 | 3 |
| 42 | Leu | | 58.355 | 4.079 | -34.480 | 3.618 | 0.016 | 5 |
| 43 | Leu | | 66.912 | 7.477 | -51.671 | 7.593 | 0.023 | 5 |
| 44 | Leu | | 171.111 | 49.391 | -132.096 | 45.017 | 0.053 | 5 |
| 45 | Leu | | 105.832 | 5.836 | -65.833 | 4.695 | 0.013 | 5 |
| 46 | Leu | | 95.182 | 21.044 | -83.300 | 22.334 | 0.041 | 5 |
| 47 | Leu | | 125.173 | 43.406 | -116.837 | 47.518 | 0.059 | 5 |
| 48 | Leu | | 98.491 | 12.086 | -104.496 | 15.029 | 0.019 | 5 |
| 49 | Leu | | 121.624 | 15.401 | -84.017 | 13.213 | 0.027 | 5 |
| 50 | Val | | 135.871 | 50.395 | -133.764 | 59.275 | 0.058 | 4 |
| 51 | Val | | 132.761 | 25.589 | -117.877 | 29.309 | 0.032 | 3 |
| 52 | Val | | 36.138 | 5.016 | -17.851 | 5.250 | 0.031 | 5 |
| 53 | Val | | 168.971 | 18.536 | -96.962 | 13.368 | 0.025 | 5 |
| 54 | Val | Leu? | 168.203 | 28.447 | -85.103 | 21.029 | 0.040 | 3 |
| 55 | Val | | 192.961 | 9.739 | -123.519 | 7.546 | 0.010 | 5 |
| 56 | Val | | 177.957 | 62.216 | -153.410 | 62.027 | 0.058 | 5 |
| 57 | Val | Leu? | 254.051 | 22.569 | -143.916 | 16.787 | 0.017 | 3 |
| 58 | Val | Leu? | 140.444 | 27.957 | -75.199 | 22.416 | 0.046 | 3 |
| 59 | Val | | 165.055 | 22.545 | -108.697 | 18.122 | 0.029 | 5 |

| | | | | | | | |
|---|---|---|---|---|---|---|---|
| 60 | Val | 185.589 | 31.466 | -177.238 | 34.158 | 0.025 | 5 |
| 61 | Val | 245.792 | 58.877 | -233.631 | 62.278 | 0.032 | 5 |
| 62 | Val | 221.391 | 40.650 | -111.319 | 25.820 | 0.042 | 5 |
| 63 | Val | 162.949 | 36.679 | -136.269 | 35.879 | 0.039 | 5 |
| 64 | Val | 238.514 | 66.827 | -244.977 | 77.348 | 0.034 | 4 |
| 65 | Val | 182.487 | 18.206 | -100.305 | 12.615 | 0.023 | 5 |
| 66 | Val | 157.192 | 11.969 | -89.274 | 8.627 | 0.018 | 5 |
| 67 | Val | 154.315 | 19.155 | -112.699 | 16.823 | 0.024 | 5 |
| 68 | Val | 231.596 | 17.133 | -147.281 | 14.061 | 0.014 | 3 |
| 69 | Val | 175.067 | 26.325 | -129.740 | 23.154 | 0.028 | 5 |

**Table S12. Rotational correlation times** of $\alpha_{1B}$-AR-B1D1 binding **prazosin**. Standard errors for the rotational correlation times were estimated using 1 000 Monte Carlo samplings based on the standard errors of the α- and β-rates. Only values that were used to calculate the global rotational correlation time are listed (see Appendix A section 3 for details).

| Peak Nr. | $\tau_c$ [ns] | SE $\tau_c$ [ns] | $R_\alpha$ [$s^{-1}$] | $R_\alpha$ SE [$s^{-1}$] | $R_\beta$ [$s^{-1}$] | $R_\beta$ SE [$s^{-1}$] |
|---|---|---|---|---|---|---|
| 1 | 23.908 | 4.303 | 43.766 | 3.564 | 95.554 | 8.395 |
| 2 | 34.857 | 3.495 | 18.936 | 1.275 | 94.182 | 7.488 |
| 3 | 44.764 | 3.641 | 20.218 | 0.617 | 116.735 | 7.890 |
| 4 | 34.896 | 5.089 | 22.115 | 1.617 | 97.446 | 10.944 |
| 5 | 31.001 | 1.809 | 16.919 | 0.803 | 83.897 | 3.737 |
| 6 | 40.856 | 5.606 | 20.438 | 1.869 | 108.562 | 12.058 |
| 7 | 44.193 | 5.286 | 22.003 | 1.374 | 117.292 | 11.524 |
| 8 | 34.747 | 2.612 | 28.102 | 1.661 | 103.114 | 5.487 |
| 9 | 44.700 | 3.889 | 27.866 | 1.244 | 124.245 | 8.460 |
| 10 | 37.785 | 4.588 | 29.500 | 2.510 | 111.031 | 9.459 |
| 11 | 31.758 | 6.026 | 22.227 | 1.815 | 90.828 | 12.706 |
| 12 | 40.201 | 8.656 | 14.680 | 3.458 | 101.397 | 18.680 |
| 13 | 29.208 | 3.970 | 29.805 | 3.045 | 92.940 | 8.413 |
| 14 | 50.911 | 4.667 | 17.564 | 0.954 | 127.287 | 10.022 |
| 15 | 40.941 | 4.007 | 35.452 | 3.074 | 123.757 | 8.290 |
| 16 | 37.674 | 3.236 | 18.384 | 1.308 | 99.676 | 6.493 |
| 17 | 30.599 | 4.390 | 22.858 | 1.691 | 88.973 | 9.459 |
| 18 | 21.304 | 1.310 | 18.192 | 0.814 | 64.417 | 2.736 |
| 19 | 35.895 | 2.368 | 22.221 | 0.889 | 99.694 | 5.048 |
| 20 | 35.746 | 7.156 | 24.475 | 2.134 | 101.629 | 15.167 |
| 21 | 48.218 | 7.121 | 23.664 | 1.829 | 127.600 | 15.364 |
| 22 | 36.031 | 2.252 | 22.068 | 0.924 | 99.835 | 4.707 |
| 23 | 43.939 | 4.956 | 24.081 | 1.331 | 118.826 | 10.240 |
| 24 | 44.680 | 2.714 | 22.976 | 1.525 | 119.313 | 5.572 |
| 25 | 49.673 | 11.075 | 24.115 | 2.843 | 131.178 | 22.918 |
| 26 | 36.978 | 3.364 | 16.889 | 1.745 | 96.687 | 6.762 |
| 27 | 34.063 | 4.327 | 21.072 | 1.131 | 94.616 | 9.219 |
| 28 | 35.754 | 7.488 | 23.952 | 2.163 | 101.123 | 15.492 |
| 29 | 40.700 | 2.989 | 17.754 | 1.384 | 105.543 | 6.439 |
| 30 | 34.246 | 2.002 | 25.006 | 1.233 | 98.942 | 4.079 |
| 31 | 38.684 | 2.441 | 20.309 | 1.010 | 103.769 | 5.089 |
| 32 | 25.385 | 2.379 | 22.022 | 1.017 | 76.971 | 4.908 |
| 33 | 43.894 | 4.867 | 15.305 | 1.612 | 109.953 | 10.197 |
| 34 | 29.264 | 4.814 | 25.185 | 1.418 | 88.439 | 10.467 |
| 35 | 33.134 | 5.553 | 40.187 | 3.967 | 111.738 | 10.972 |
| 36 | 31.204 | 1.357 | 19.370 | 0.942 | 86.783 | 2.774 |
| 37 | 48.317 | 8.427 | 9.442 | 2.988 | 113.592 | 18.468 |
| 38 | 41.630 | 3.800 | 20.353 | 1.282 | 110.139 | 7.817 |
| 39 | 40.400 | 9.074 | 15.805 | 1.456 | 102.950 | 19.081 |
| 40 | 33.495 | 2.625 | 21.636 | 0.931 | 93.961 | 5.569 |
| 41 | 35.607 | 2.285 | 20.332 | 1.244 | 97.188 | 4.704 |
| 42 | 34.708 | 2.294 | 23.052 | 0.946 | 97.979 | 4.803 |
| 43 | 26.209 | 11.620 | 49.966 | 6.392 | 106.678 | 24.039 |
| 44 | 38.649 | 1.331 | 16.345 | 0.793 | 99.729 | 2.750 |
| 45 | 39.825 | 2.498 | 23.466 | 0.864 | 109.375 | 5.206 |
| 46 | 33.076 | 2.829 | 24.461 | 1.170 | 95.889 | 5.919 |
| 47 | 51.758 | 8.911 | 14.649 | 2.739 | 126.193 | 18.224 |
| 48 | 38.806 | 3.006 | 23.523 | 1.543 | 107.244 | 6.506 |
| 49 | 23.428 | 3.242 | 21.436 | 1.539 | 72.198 | 6.494 |
| 50 | 37.623 | 5.566 | 21.028 | 1.685 | 102.211 | 12.065 |
| 51 | 32.963 | 6.004 | 20.897 | 2.390 | 92.082 | 12.488 |
| 52 | 43.743 | 6.205 | 24.502 | 1.472 | 118.826 | 13.311 |
| 53 | 33.399 | 4.269 | 21.768 | 1.417 | 93.887 | 9.067 |
| 54 | 40.149 | 11.359 | 32.863 | 5.303 | 119.469 | 23.202 |
| 55 | 34.819 | 2.446 | 17.689 | 0.611 | 92.854 | 5.209 |
| 56 | 39.974 | 2.373 | 21.401 | 0.892 | 107.630 | 5.028 |
| 57 | 44.888 | 1.343 | 19.298 | 0.900 | 116.082 | 2.809 |
| 58 | 34.148 | 5.587 | 22.509 | 1.856 | 96.234 | 11.300 |
| 59 | 30.200 | 2.405 | 15.462 | 0.863 | 80.723 | 5.199 |
| 60 | 33.379 | 2.547 | 23.737 | 0.961 | 95.814 | 5.295 |
| 61 | 35.960 | 10.649 | 19.844 | 2.979 | 97.457 | 22.731 |

| | | | | | | |
|---|---|---|---|---|---|---|
| 62 | 25.133 | 6.537 | 16.049 | 2.248 | 70.458 | 13.884 |
| 63 | 37.322 | 2.466 | 15.101 | 0.787 | 95.639 | 5.206 |
| 64 | 26.731 | 1.842 | 22.967 | 1.388 | 80.797 | 3.558 |
| 65 | 34.223 | 3.204 | 20.452 | 0.817 | 94.339 | 6.840 |
| 66 | 43.975 | 3.945 | 16.143 | 1.155 | 110.966 | 8.520 |
| 67 | 30.382 | 3.610 | 19.678 | 1.749 | 85.328 | 7.509 |
| 68 | 44.175 | 1.442 | 15.161 | 0.489 | 110.412 | 3.123 |
| 69 | 28.771 | 5.151 | 27.619 | 2.987 | 89.816 | 10.682 |
| 70 | 43.797 | 5.357 | 25.928 | 1.506 | 120.367 | 10.826 |
| 71 | 46.867 | 5.035 | 14.195 | 1.228 | 115.228 | 11.032 |
| 72 | 40.280 | 7.714 | 18.819 | 1.614 | 105.706 | 16.966 |
| 73 | 46.806 | 4.776 | 20.703 | 1.340 | 121.607 | 10.147 |
| 74 | 31.496 | 8.820 | 28.932 | 3.766 | 96.970 | 18.311 |
| 75 | 48.098 | 3.010 | 20.618 | 0.945 | 124.297 | 6.702 |
| 76 | 30.190 | 3.269 | 20.594 | 1.437 | 85.832 | 6.938 |
| 77 | 38.225 | 4.566 | 22.631 | 1.307 | 105.106 | 9.451 |
| 78 | 43.689 | 4.458 | 22.497 | 1.389 | 116.706 | 9.098 |
| 79 | 41.318 | 4.625 | 18.290 | 1.511 | 107.405 | 9.798 |
| 80 | 32.325 | 9.124 | 16.935 | 2.147 | 86.752 | 19.508 |
| 81 | 26.059 | 1.050 | 21.378 | 0.495 | 77.768 | 2.335 |
| 82 | 39.302 | 2.608 | 20.004 | 0.890 | 104.791 | 5.472 |
| 83 | 47.916 | 6.829 | 16.040 | 2.204 | 119.328 | 14.684 |
| 84 | 34.162 | 3.233 | 19.604 | 1.191 | 93.360 | 6.652 |
| 85 | 32.320 | 3.025 | 20.006 | 1.614 | 89.812 | 6.224 |
| 86 | 30.344 | 2.267 | 26.633 | 0.952 | 92.203 | 4.888 |
| 87 | 44.683 | 6.216 | 23.727 | 2.023 | 120.069 | 13.808 |
| 88 | 38.740 | 6.525 | 28.938 | 3.509 | 112.517 | 14.053 |
| 89 | 41.074 | 6.508 | 25.126 | 3.323 | 113.718 | 13.594 |
| 90 | 38.655 | 6.498 | 24.229 | 1.900 | 107.627 | 14.374 |
| 91 | 26.541 | 1.582 | 20.733 | 0.748 | 78.155 | 3.265 |
| 92 | 39.404 | 1.610 | 21.138 | 0.922 | 106.145 | 3.370 |
| 93 | 35.812 | 2.733 | 18.035 | 0.743 | 95.330 | 5.951 |
| 94 | 30.605 | 5.036 | 20.585 | 2.039 | 86.712 | 10.961 |
| 95 | 36.460 | 1.963 | 18.963 | 0.627 | 97.650 | 4.194 |
| 96 | 41.382 | 7.620 | 18.063 | 1.363 | 107.316 | 15.556 |
| 97 | 52.544 | 7.678 | 18.919 | 3.667 | 132.153 | 16.203 |
| 98 | 49.118 | 4.862 | 22.175 | 1.653 | 128.046 | 10.251 |
| 99 | 26.495 | 3.486 | 25.415 | 2.073 | 82.739 | 7.116 |
| 100 | 41.850 | 1.812 | 18.369 | 0.831 | 108.627 | 3.896 |
| 101 | 27.456 | 9.506 | 20.345 | 1.975 | 79.726 | 19.730 |
| 102 | 47.743 | 6.745 | 22.020 | 1.820 | 124.936 | 14.390 |
| 103 | 30.240 | 4.742 | 16.935 | 2.255 | 82.280 | 9.983 |
| 104 | 48.443 | 5.922 | 25.246 | 2.952 | 129.666 | 12.454 |
| 105 | 34.322 | 4.225 | 19.625 | 1.525 | 93.725 | 8.858 |
| 106 | 44.694 | 6.075 | 21.349 | 1.556 | 117.715 | 13.367 |
| 107 | 32.104 | 1.905 | 24.155 | 0.751 | 93.497 | 4.118 |
| 108 | 25.476 | 6.342 | 41.717 | 4.061 | 96.860 | 12.734 |
| 109 | 37.161 | 3.735 | 16.608 | 2.019 | 96.798 | 7.884 |
| 110 | 38.350 | 3.176 | 19.841 | 0.972 | 102.585 | 6.753 |
| 111 | 36.900 | 2.898 | 20.052 | 1.019 | 99.683 | 6.274 |
| 112 | 32.174 | 4.865 | 18.770 | 1.581 | 88.263 | 10.220 |
| 113 | 27.370 | 9.760 | 23.974 | 5.001 | 83.172 | 20.824 |

**Table S13. Rotational correlation times** of $\alpha_{1B}$-AR-B1D1 binding **ρ-TIA**. Standard errors for the rotational correlation times were estimated using 1 000 Monte Carlo samplings based on the standard errors of the α- and β-rates. Only values that were used to calculate the global rotational correlation time are listed (see Appendix A section 3 for details).

| Peak Nr. | $\tau_c$ [ns] | SE $\tau_c$ [ns] | $R_\alpha$ [$s^{-1}$] | $R_\alpha$ SE [$s^{-1}$] | $R_\beta$ [$s^{-1}$] | $R_\beta$ SE [$s^{-1}$] |
|---|---|---|---|---|---|---|
| 1 | 37.724 | 3.550 | 16.142 | 1.559 | 97.541 | 7.362 |
| 2 | 55.554 | 9.499 | 19.681 | 2.192 | 139.384 | 19.935 |
| 3 | 36.576 | 4.224 | 17.891 | 1.345 | 96.827 | 9.062 |
| 4 | 41.401 | 7.902 | 22.539 | 2.522 | 111.832 | 17.043 |
| 5 | 32.019 | 5.557 | 17.401 | 1.432 | 86.560 | 12.166 |
| 6 | 38.893 | 1.664 | 17.117 | 0.804 | 101.026 | 3.453 |
| 7 | 33.799 | 5.871 | 16.896 | 1.160 | 89.874 | 12.225 |
| 8 | 35.225 | 3.152 | 24.425 | 1.869 | 100.461 | 6.328 |
| 9 | 57.417 | 8.437 | 23.140 | 1.889 | 146.847 | 18.721 |
| 10 | 34.823 | 3.912 | 24.687 | 2.282 | 99.862 | 7.891 |
| 11 | 29.778 | 5.169 | 21.433 | 2.039 | 85.789 | 10.623 |
| 12 | 27.716 | 3.046 | 16.468 | 1.369 | 76.406 | 6.592 |
| 13 | 40.588 | 9.051 | 30.402 | 2.691 | 117.949 | 19.178 |
| 14 | 47.958 | 8.135 | 26.511 | 2.302 | 129.890 | 17.862 |
| 15 | 36.647 | 8.223 | 14.954 | 4.619 | 94.043 | 16.719 |
| 16 | 42.562 | 1.874 | 15.990 | 0.810 | 107.778 | 3.944 |
| 17 | 58.036 | 7.134 | 18.443 | 1.933 | 143.481 | 16.000 |
| 18 | 32.552 | 4.774 | 24.361 | 1.841 | 94.663 | 10.035 |
| 19 | 37.402 | 9.903 | 21.655 | 2.592 | 102.365 | 21.060 |
| 20 | 37.723 | 4.894 | 18.905 | 1.805 | 100.302 | 10.328 |
| 21 | 34.189 | 3.553 | 21.279 | 1.104 | 95.093 | 7.692 |
| 22 | 29.314 | 5.863 | 16.397 | 2.499 | 79.758 | 12.366 |
| 23 | 35.344 | 3.620 | 19.500 | 0.992 | 95.793 | 7.484 |
| 24 | 39.402 | 4.475 | 20.558 | 1.410 | 105.561 | 9.574 |
| 25 | 39.829 | 7.978 | 17.088 | 2.530 | 103.006 | 17.377 |
| 26 | 30.489 | 2.954 | 18.272 | 0.772 | 84.151 | 6.204 |
| 27 | 32.728 | 2.012 | 21.062 | 0.780 | 91.743 | 4.424 |
| 28 | 43.590 | 3.109 | 16.175 | 1.195 | 110.170 | 6.534 |
| 29 | 44.920 | 5.877 | 21.419 | 1.566 | 118.270 | 12.553 |
| 30 | 23.867 | 6.869 | 35.828 | 4.398 | 87.528 | 13.916 |
| 31 | 49.219 | 11.236 | 20.828 | 3.024 | 126.916 | 24.289 |
| 32 | 39.759 | 8.099 | 22.145 | 2.121 | 107.913 | 16.822 |
| 33 | 43.864 | 4.572 | 24.568 | 2.444 | 119.151 | 9.996 |
| 34 | 25.134 | 4.029 | 25.101 | 2.367 | 79.512 | 8.416 |
| 35 | 35.223 | 3.222 | 23.354 | 1.517 | 99.387 | 6.731 |
| 36 | 29.650 | 3.579 | 22.822 | 1.619 | 86.903 | 7.524 |
| 37 | 31.400 | 2.880 | 17.901 | 0.963 | 85.733 | 5.980 |
| 38 | 32.136 | 6.582 | 20.164 | 2.401 | 89.574 | 12.841 |
| 39 | 43.841 | 7.575 | 21.247 | 2.463 | 115.781 | 15.980 |
| 40 | 34.656 | 3.745 | 19.339 | 1.069 | 94.155 | 7.691 |
| 41 | 30.344 | 2.953 | 20.581 | 1.560 | 86.149 | 5.936 |
| 42 | 37.558 | 4.371 | 20.221 | 1.522 | 101.264 | 8.705 |
| 43 | 39.888 | 8.301 | 13.440 | 3.001 | 99.484 | 17.478 |
| 44 | 34.184 | 3.805 | 19.712 | 1.536 | 93.515 | 8.147 |
| 45 | 37.060 | 6.283 | 23.637 | 1.483 | 103.610 | 12.933 |
| 46 | 28.309 | 6.727 | 38.323 | 3.263 | 99.531 | 13.862 |
| 47 | 36.735 | 5.539 | 24.943 | 1.913 | 104.220 | 11.782 |
| 48 | 39.981 | 6.311 | 26.276 | 2.738 | 112.521 | 13.782 |
| 49 | 36.403 | 2.332 | 20.909 | 1.560 | 99.474 | 4.749 |
| 50 | 39.606 | 1.966 | 19.321 | 0.993 | 104.762 | 4.083 |
| 51 | 43.719 | 5.010 | 18.492 | 1.172 | 112.763 | 10.750 |
| 52 | 38.145 | 9.354 | 20.929 | 3.361 | 103.233 | 19.611 |
| 53 | 34.155 | 2.731 | 20.593 | 1.066 | 94.335 | 5.623 |
| 54 | 27.699 | 11.889 | 28.416 | 6.532 | 88.318 | 24.484 |
| 55 | 33.219 | 5.302 | 23.443 | 1.810 | 95.177 | 11.216 |
| 56 | 36.884 | 3.926 | 17.781 | 1.529 | 97.377 | 8.423 |
| 57 | 31.564 | 10.287 | 15.561 | 2.228 | 83.746 | 22.442 |
| 58 | 41.653 | 6.505 | 22.242 | 2.541 | 112.077 | 13.787 |
| 59 | 33.383 | 3.534 | 17.861 | 1.230 | 89.946 | 7.525 |
| 60 | 42.559 | 5.921 | 22.268 | 1.468 | 114.050 | 12.851 |
| 61 | 49.216 | 4.571 | 17.882 | 0.819 | 123.962 | 9.705 |

| | | | | | | |
|---|---|---|---|---|---|---|
| 62 | 28.530 | 4.424 | 27.766 | 2.809 | 89.449 | 8.684 |
| 63 | 49.813 | 11.497 | 16.426 | 2.753 | 123.790 | 24.056 |
| 64 | 38.847 | 2.066 | 19.665 | 0.715 | 103.476 | 4.361 |
| 65 | 20.601 | 2.447 | 20.322 | 1.500 | 65.047 | 5.003 |
| 66 | 23.737 | 1.182 | 18.629 | 0.920 | 70.053 | 2.429 |
| 67 | 29.250 | 2.587 | 20.296 | 1.169 | 83.521 | 5.356 |
| 68 | 37.771 | 3.082 | 22.169 | 1.571 | 103.669 | 6.369 |
| 69 | 33.834 | 3.110 | 22.761 | 1.475 | 95.813 | 6.586 |
| 70 | 32.301 | 8.117 | 25.710 | 2.300 | 95.474 | 17.570 |
| 71 | 31.818 | 6.039 | 24.966 | 2.091 | 93.694 | 12.925 |
| 72 | 35.758 | 3.973 | 27.783 | 1.821 | 104.964 | 8.585 |
| 73 | 30.494 | 5.843 | 20.388 | 2.885 | 86.278 | 11.966 |
| 74 | 33.134 | 3.141 | 23.702 | 1.487 | 95.253 | 6.562 |
| 75 | 38.675 | 4.928 | 28.074 | 1.436 | 111.514 | 10.834 |
| 76 | 32.280 | 8.901 | 27.762 | 3.613 | 97.481 | 18.454 |
| 77 | 34.496 | 5.334 | 22.980 | 1.348 | 97.451 | 11.024 |
| 78 | 44.855 | 8.520 | 18.450 | 2.167 | 115.162 | 18.618 |
| 79 | 45.759 | 8.615 | 22.605 | 2.236 | 121.260 | 18.336 |
| 80 | 34.234 | 6.041 | 21.242 | 2.180 | 95.152 | 12.471 |
| 81 | 46.037 | 4.956 | 18.338 | 1.895 | 117.589 | 10.571 |
| 82 | 27.980 | 5.765 | 22.886 | 3.252 | 83.390 | 11.790 |
| 83 | 40.524 | 4.935 | 24.160 | 1.809 | 111.570 | 10.462 |
| 84 | 20.735 | 4.585 | 13.353 | 2.658 | 58.364 | 9.251 |
| 85 | 32.267 | 6.515 | 23.460 | 2.059 | 93.150 | 13.645 |
| 86 | 42.991 | 10.552 | 20.319 | 2.132 | 113.029 | 22.406 |
| 87 | 24.832 | 9.218 | 21.822 | 2.769 | 75.587 | 19.668 |
| 88 | 34.273 | 4.381 | 22.318 | 2.651 | 96.313 | 9.212 |
